# A cuffed CRISPR guide RNA for microRNA activity-dependent genome editing

**DOI:** 10.64898/2026.08.08.743703

**Authors:** Arman Adel, Yutaro Shuto, Shunsuke Kawasaki, Yonggang Lu, Hiroki Ono, Satoshi N. Omura, Ryoya Nakagawa, Alima Suleimenova, Meghan Robinson, Tabea L. Stephan, Hideto Mori, Brett Kiyota, Sanchit Chopra, Nemekhbayar Baatartsogt, Morisada Hayakawa, Yuji Kashiwakura, Tsukasa Ohmori, Ryan Flannigan, T. Michael Underhill, Pamela A. Hoodless, Hiroyuki Aburatani, Hirohide Saito, Osamu Nureki, Nozomu Yachie

## Abstract

Cells in multicellular eukaryotic systems are diverse biological units, with characteristics and functions determined by their molecular profiles. CRISPR–Cas9 genome editing has been widely used across biology to modulate gene expression and study gene function. However, there is currently no versatile and scalable method for editing a cell’s genome in response to endogenous cellular signals. Here, we report the engineering of a CRISPR guide RNA that efficiently confers genome editing in response to the catalytic activity of a target microRNA (miRNA) within a cell. miRNAs are short non-coding RNAs that are widely conserved across eukaryotes and can cleave their target RNA through almost perfect base pairing. In mammals, miRNAs are largely involved in development and homeostasis as well as disease progression and developmental disorders. To leverage these properties for genome editing, we developed a cuffed guide RNA (cgRNA) which is composed of a permutated order of sequence domains from the commonly used single guide RNA (sgRNA). These permutated domains were then concatenated with a miRNA target sequence, yielding a warped guide RNA that is inactive until cleaved by a complementary miRNA. We demonstrated that cgRNA enabled efficient miRNA activity-dependent genome editing in human and mouse cell lines. Biochemical and structural analyses revealed three stages of inhibition of the CRISPR genome-editing pathway for unprocessed cgRNA. Utilizing a lentiviral library of cgRNAs containing miRNA targets covering mouse genome-wide miRNAs, we identified miRNA cleavage activities and their sequence specificities in mouse embryonic stem cells and during smooth muscle cell differentiation. Furthermore, we showed that endogenous mRNA expression could be irreversibly recorded into a DNA sequence using a cgRNA targeted by a synthetic miRNA repeat. cgRNA is a simple, robust, miRNA activity-gated genome editing system that could facilitate the development of cell state-specific genome editing, the mapping of miRNA activity and gene expression landscapes, and the recording of molecularly determined cell states during the long-term progression of multicellular systems.

## INTRODUCTION

### No current genome editing tool is scalably responsive to cellular contexts

Clustered regularly interspaced short palindromic repeat (CRISPR)–Cas9 genome editing has sparked a revolution in the life sciences^1^. Its ongoing and potential applications span from the engineering of cell and animal models^2^ to genome-scale phenotypic surveys of gene knockouts^3^, overexpression^4^, and epigenetic changes^5^, as well as the correction of genetic disorders for therapeutic purposes^6,7^. Furthermore, many recent molecular event recording methods propose the use of CRISPR-Cas9 genome editing tools for the recording of developmental cell lineages and gene expressions into a “DNA tape” encoded in a cell’s genome^8^. The advancement of these applications has pioneered diverse approaches for temporal or spatial control of genome editing based on stimuli such as small molecules^9,10^, light of different wavelengths^11,12^, and exogenous genetic promoters^13,14^ or enhancers responsive to the molecular context of a cell^15^.

However, there is currently no scalable and efficient genome editing system that triggers editing in direct response to a target cellular event. Such a system would enable genetic perturbation studies of complex multicellular systems based on prior knowledge of their molecular context. Additionally, several studies in synthetic biology have shown that the precise output control of genetic circuits can be achieved by the programmed alteration of DNA sequences^13,14,16^, as opposed to gene expression control-based approaches which often suffer from signal leakage. As such, the direct conversion of a biological signal to genome editing would hold great promise in diverse applications, including cell engineering and therapeutics.

### Genome editing for DNA event recording

Conversion of a biological signal into genome editing is a particularly intriguing challenge in the emerging field of molecular event recording. The field continues to envision the recording of temporal cellular and molecular information in stable biomolecules within cells to facilitate the readout of dynamic behaviors in living systems^13,15,17–22^. Current scalable measurement methods for cells only capture a snapshot of molecules in complex multicellular systems due to the requirement of sample destruction for observation. Molecular event recording aims to read out stored information and reconstruct system dynamics retrospectively to overcome this limitation.

This field was spearheaded by the development of cell lineage tracing, which uses CRISPR-Cas9 genome editing to generate semi-random mutations in a DNA tape as cells proliferate ^23,24^. It was proposed that such systems could capture cell division histories by reconstructing their lineage from the mutation patterns recorded within each cell’s DNA tape at the time of observation, in a manner similar to evolutionary tree reconstruction from species’ genome sequences. Furthermore, several approaches have demonstrated that transcription of DNA tape sequences enables simultaneous readout of the transcriptomic state and developmental cell lineage of single cells by single-cell RNA sequencing (scRNA-seq)^25–27^.

In addition to lineage recording, molecular DNA event recording could theoretically be applied to the reconstruction of past cell state transitions if these states are represented by a set of molecular activities that can also be recorded in corresponding DNA tapes and captured alongside cell lineage information^24^. Currently, ENGRAM^15^, a method for the recording of *cis*-regulatory element activity using Prime Editing, is the only method that has demonstrated the potential to record molecular activities in DNA at scale. However, ENGRAM requires the addition of transgene reporters, and its recordings do not directly measure endogenous gene expression under a genomic context.

Orthogonal to the use of DNA sequence as an information storage medium, numerous recording regimes have been developed to directly store high-content molecular profiles of cells using proteins^17,18^, protein cargoes^19,21^, RNAs^20^, and epigenetic modifications^22^. However, these recording modalities do not enable stable long-term information storage like DNA tapes, highlighting the need for scalable and longitudinal methods to record diverse cellular events.

### Converting cellular microRNA signals into DNA sequences

MicroRNAs (miRNAs) are short, non-coding RNAs that play major roles in regulating gene expression through translational repression and post-transcriptional degradation of target RNA transcripts^28,29^. They are widely conserved across diverse multicellular eukaryotes from nematodes to humans^30^. miRNA-mediated gene silencing is carried out by the RNA-induced silencing complex (RISC) and operates through two modes: endonucleolytic cleavage of a target messenger RNA (mRNA) or translational repression coupled with mRNA destabilization and decay. Direct cleavage results from nearly perfect annealing between the RNA target and its corresponding miRNA^31^. Efficient translational repression followed by degradation can result from interactions as limited as base pairing between the 6-mer seed sequence of the miRNA and the target RNA^29^. As post-transcriptional regulators of RNAs, miRNAs have also been observed to have cell state or tissue specificity in their expression patterns^32–35^, play a major role in modulating gene expression patterns within those cells^36–38^, and their evolutionarily conserved mechanism is considered to have a wide impact on developmental processes^39^.

Around 1,900 and 1,200 miRNAs have been reported in the human and mouse genomes, respectively, in the latest version of the miRNA database (miRbase v22)^40^ and the most recent CLIP-seq and PAR-CLIP meta-analysis found 1,433,920 human and 181,606 mouse miRNA interactions with target mRNA sequences^41^. Additionally, many congenital developmental disorders have been linked to mutations in miRNA^42–50^. Accordingly, scalable miRNA activity-dependent genome editing will be of great interest in the broader life sciences, including therapeutics, biotechnology, molecular biology, cell biology, and developmental biology. Furthermore, miRNA signal-gated genome editing could serve as a significant modality in DNA-based molecular event recording if it can be scaled effectively.

At present, only two approaches translate miRNA activity into a genome editing outcome using the CRISPR-Cas9 system. The first system modulates Cas9 abundance by repressing either Cas9 mRNA or an anti-CRISPR (Acr) mRNA by a target miRNA of interest^51,52^. However, since the regulation of editing outcomes is at the Cas9 protein level, the number of miRNA activities that can be simultaneously recorded in a cell (multiplexability) by this system is limited by the number of orthogonal Cas9 or Acr proteins. The second system processes a mature CRISPR single-guide RNA (sgRNA) from an RNA polymerase II (Pol2) transcript (inactive as sgRNA) in response to a specific miRNA that cleaves off both the 5′ cap and polyadenylation (poly(A)) tail through perfect base pairing^53^. This system, called MICR (miRNA-induced CRISPR-Cas9 platform), requires a long Pol2 promoter to transcribe the inactive form of the sgRNA and two miRNA target sequences to remove the upstream and downstream Pol2 transcript modifications. Therefore, it is not suitable to construct a condensed circuit that responds to many miRNA activities or a large-scale library of single miRNA activity-dependent genome editing circuits.

### A cuffed guide RNA for miRNA activity-responsible genome editing

Here, we report a cuffed CRISPR-Cas9 guide RNA (cgRNA), which transitions to a functional state after release of its circular configuration to a linear configuration by targeted RNA cleavage. The sgRNA commonly used in genome editing is a fusion of the mature bacterial CRISPR RNA (crRNA) that encodes a spacer sequence and direct repeat and the trans-activating crRNA (tracrRNA) (**Figures 1A** and **S1A**). In the native bacterial CRISPR–Cas9 immune system, the guide RNA consists of crRNA and tracrRNA, which associate through complementary base pairing to form the secondary structure required for Cas9 binding (**Figures 1A** and **S1B**). The guide RNA then recruits Cas9 to the target DNA through sequence complementarity between the spacer and target DNA. Inspired by the natural architecture of a permuted transfer RNA we previously discovered in the red algae species *Cyanidioschyzon merolae*^54^, we swapped the order of the sgRNA domains from 5′-crRNA-tracrRNA-3′ to 5′-tracrRNA-crRNA-3′ and then inserted a miRNA target (miRT) between the permutated domains, resulting in 5′-tracrRNA-miRT-crRNA-3′ (**Figures 1B** and **S1C**). A miRNA loaded into an Argonaute 2 (AGO2)-containing RISC directs sequence-specific cleavage of a target RNA sequence through nearly perfect base pairing^31^ and is capable of cleaving within a 30-nucleotide (nt) RNA loop^55^. We hypothesized that the circular form of cgRNA is unable to confer genome editing via Cas9 without cleavage of its loop by a corresponding miRNA to restore the functional linear crRNA:tracrRNA (**Figure 1B**).

**Figure 1.**
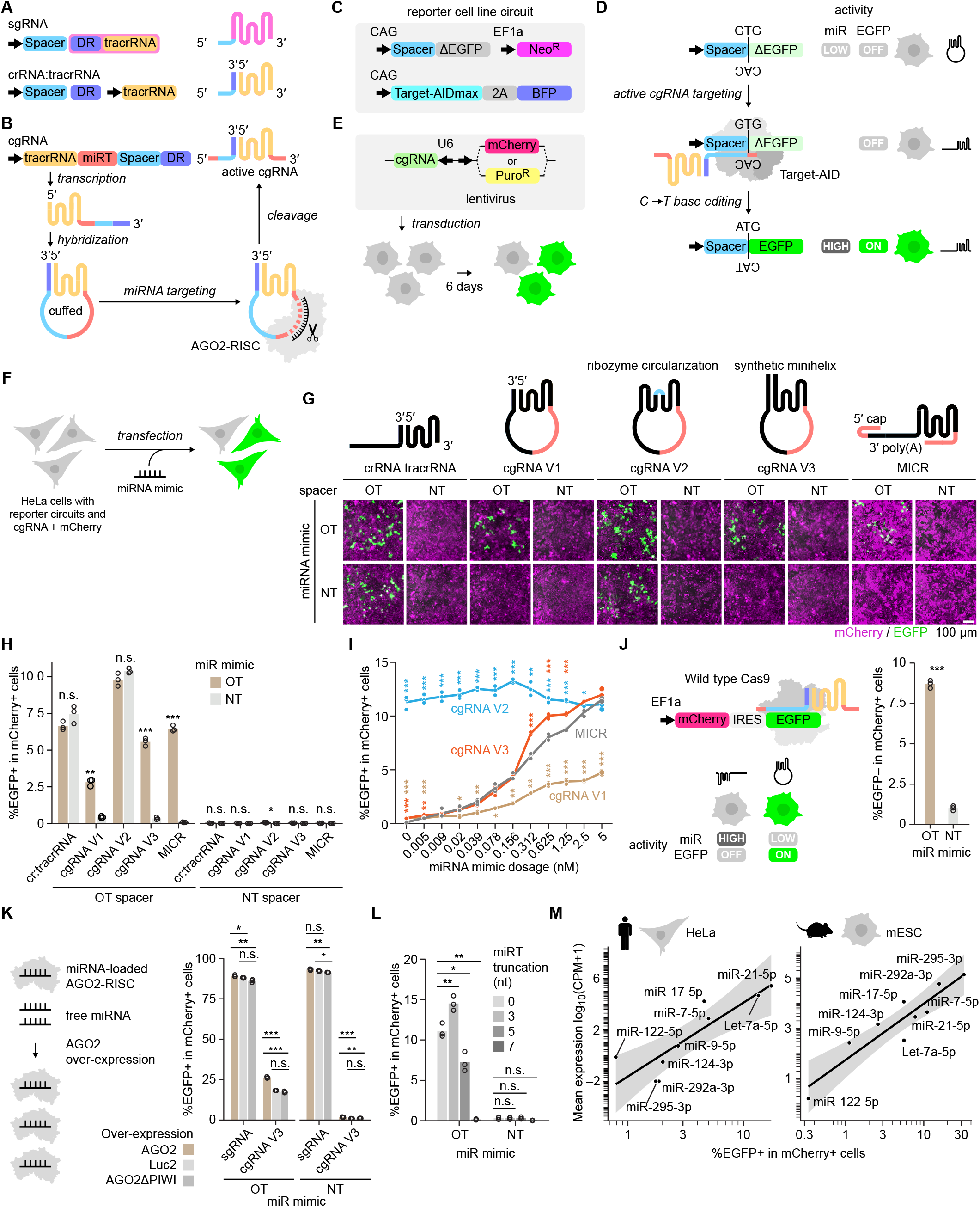
Cuffed CRISPR guide RNA enables miRNA activity-dependent genome editing. (A) The architecture of sgRNA and crRNA:tracrRNA. (B) Schematic of cgRNA and its activation by miRNA-mediated cleavage. (C) Common C→T base-editing reporter circuit used throughout this study. (D) Restoration of EGFP expression by Target-AIDmax following miRNA-dependent activation of cgRNA. (E) Lentiviral delivery of cgRNA together with either mCherry or a puromycin-resistance marker. Cells were analyzed 6 days after transduction. (F– H) Comparison of guide RNA architectures for miRNA mimic-dependent base editing. Toolkit HeLa cells carrying guide RNAs with either a reporter-targeting or non-targeting spacer were transfected with miR-122 or a non-targeting miRNA mimic. (G) Representative fluorescence micrographs. Scale bar, 100 μm. (H) Percentage of EGFP+ cells among mCherry+ cells. Welch’s t-test was used to compare miR-122 and non-targeting mimic conditions (n = 3 replicates). (I) Dose-dependent base editing by cgRNA V1–V3 and MICR following transfection with increasing concentrations of miR-122 mimic. Welch’s t-test was used to compare each cgRNA version with MICR at each dose (n = 3 replicates). (J) miRNA-dependent genome editing by cgRNA coupled to wild-type SpCas9. Following transfection with miR-122 or a non-targeting mimic, the percentage of EGFP− cells among mCherry+ cells was quantified by flow cytometry. Welch’s t-test was used for comparison (n = 3 replicates). (K) Effect of AGO2 abundance on cgRNA activity. Toolkit HeLa cells expressing AGO2, firefly luciferase (Luc2), or catalytically inactive AGO2ΔPIWI were transduced with either sgRNA or miR-122-targeted cgRNA and subsequently transfected with miR-122 or a non-targeting mimic. The percentage of EGFP+ cells among mCherry+ cells was quantified by flow cytometry. Welch’s t-test was used to compare AGO2 with the Luc2 and AGO2ΔPIWI controls (n = 3 replicates). (L) Effect of 5′ truncation of the miR-122 target sequence on cgRNA-mediated base editing. Welch’s t-test was used to compare each truncation with the untruncated target under the miR-122 mimic condition (n = 3 replicates). (M) Relationship between cgRNA editing efficiency and cellular miRNA abundance measured by AQ-seq in HeLa cells and mESCs. Each point represents the mean of three replicate measurements for one miRNA. Lines indicate linear regression fits, and gray shading indicates the 95% confidence interval. n.s., not significant; \**P* < 0.05; \*\**P* < 0.01; \*\*\**P* < 0.001.

In this report, we first develop, characterize, and optimize conditional genome editing using cgRNA in response to a given miRNA input (**Figure 1**), and then provide insights into its switching-state dynamics through biochemical and cryogenic electron microscopy (cryo-EM)-based structural analyses of a cgRNA with Cas9 and its corresponding target DNA (**Figure 2**). We then demonstrate the genome-wide characterization of miRNA activities in mouse embryonic stem cells (mESCs) (**Figure 3**), large-scale recording of miRNA activities in stem cell differentiation into smooth muscle cells (SMCs) (**Figure 4**), and mRNA expression recording in combination with a synthetic miRNA cluster (**Figure 5**). Finally, we discuss advancements and limitations of cgRNA and how cgRNA can transform current miRNA biology and molecular event recording fields with preliminary data on *in vivo* miRNA activity-gated genome editing (**Figure 6**).

**Figure 2.**
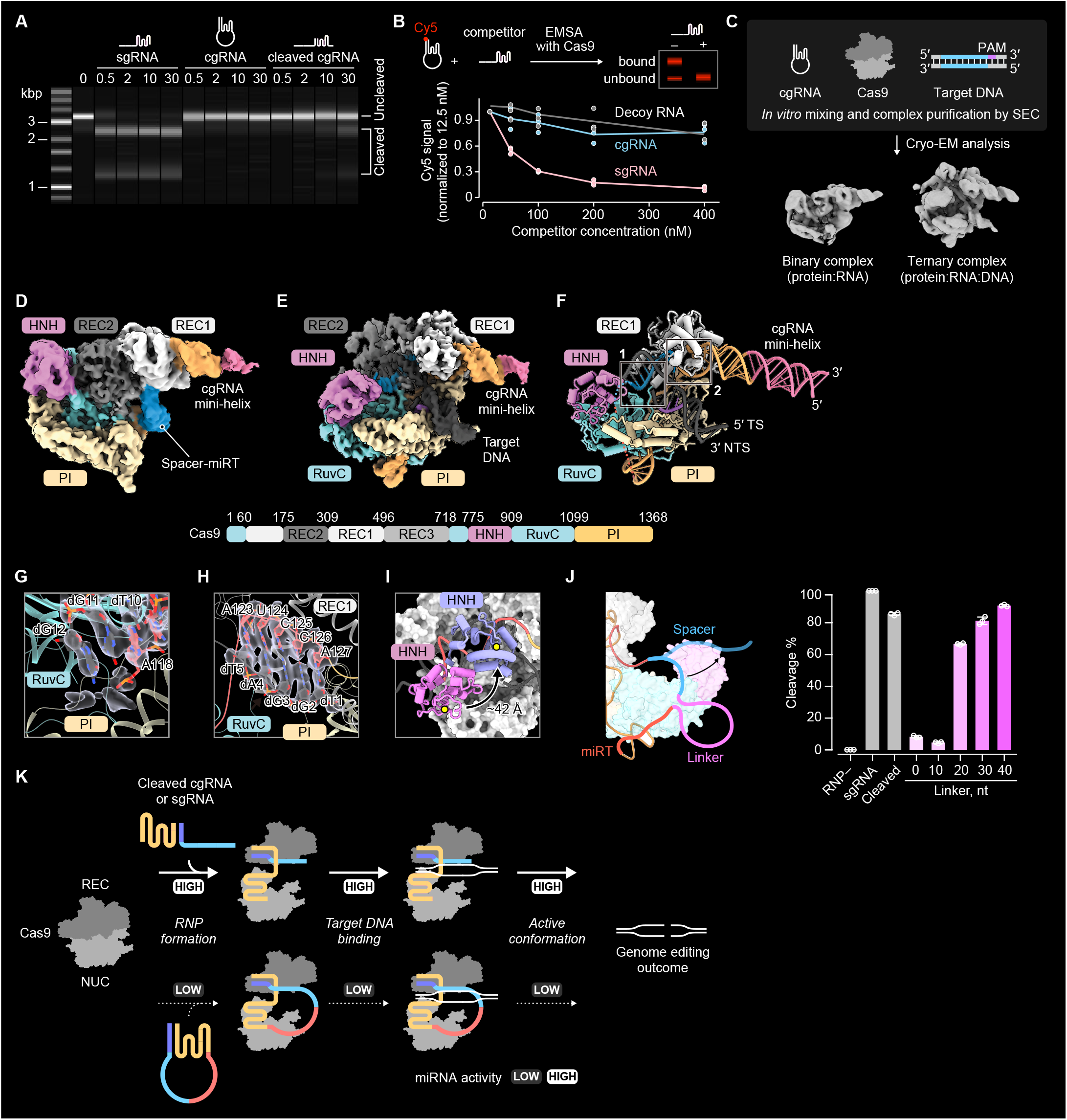
Structural basis of SpCas9 inhibition by cgRNA. (A) Time-resolved in vitro cleavage of a linearized target plasmid by SpCas9 assembled with sgRNA, intact cgRNA, or a cleaved-cgRNA mimic. Reactions were incubated at 37°C for the indicated times and analyzed by microchip electrophoresis. (B) Competitive electrophoretic mobility shift assay measuring SpCas9 binding to Cy5-labeled cgRNA in the presence of increasing concentrations of unlabeled sgRNA, cgRNA, or nonspecific competitor RNA. Signals were normalized to the lowest competitor concentration. (C) Workflow for cryo-EM analysis of SpCas9 assembled with cgRNA and target DNA, yielding binary SpCas9–cgRNA and ternary SpCas9–cgRNA–target DNA complexes. (D) Cryo-EM density map of the SpCas9–cgRNA binary complex. SpCas9 domain organization is shown below. (E, F) Cryo-EM density map (E) and structural model (F) of the SpCas9–cgRNA–target DNA ternary complex containing a partial R-loop. TS, target strand; NTS, non-target strand; PI, PAM-interacting domain. (G, H) Close-up views of the 10-bp spacer–TS heteroduplex at the PAM-distal (G; region 1 in F) and PAM-proximal (H; region 2 in F) regions. Cryo-EM densities are shown as translucent gray surfaces. (I) Comparison of HNH-domain positioning in cgRNA- and sgRNA-bound complexes. The HNH domain of the SpCas9–cgRNA–target DNA complex is shown in pink and is positioned approximately 42 Å from the predicted TS cleavage site. The HNH domain from the catalytically active SpCas9–sgRNA–target DNA complex is shown in light blue (PDB 6O0Y). The remainder of SpCas9 is shown as a gray surface, and the catalytic H840 residues are indicated by yellow stars. (J) *In vitro* DNA cleavage by SpCas9 assembled with sgRNA, a cleaved-cgRNA mimic, or intact cgRNA variants carrying linkers of the indicated lengths between the spacer and miRT. Data are mean ± s.d. (n = 3 replicates). (K) Model of the sequential inhibition imposed by intact cgRNA during SpCas9 ribonucleoprotein formation, target DNA binding, and adoption of the catalytically active conformation.

**Figure 3.**
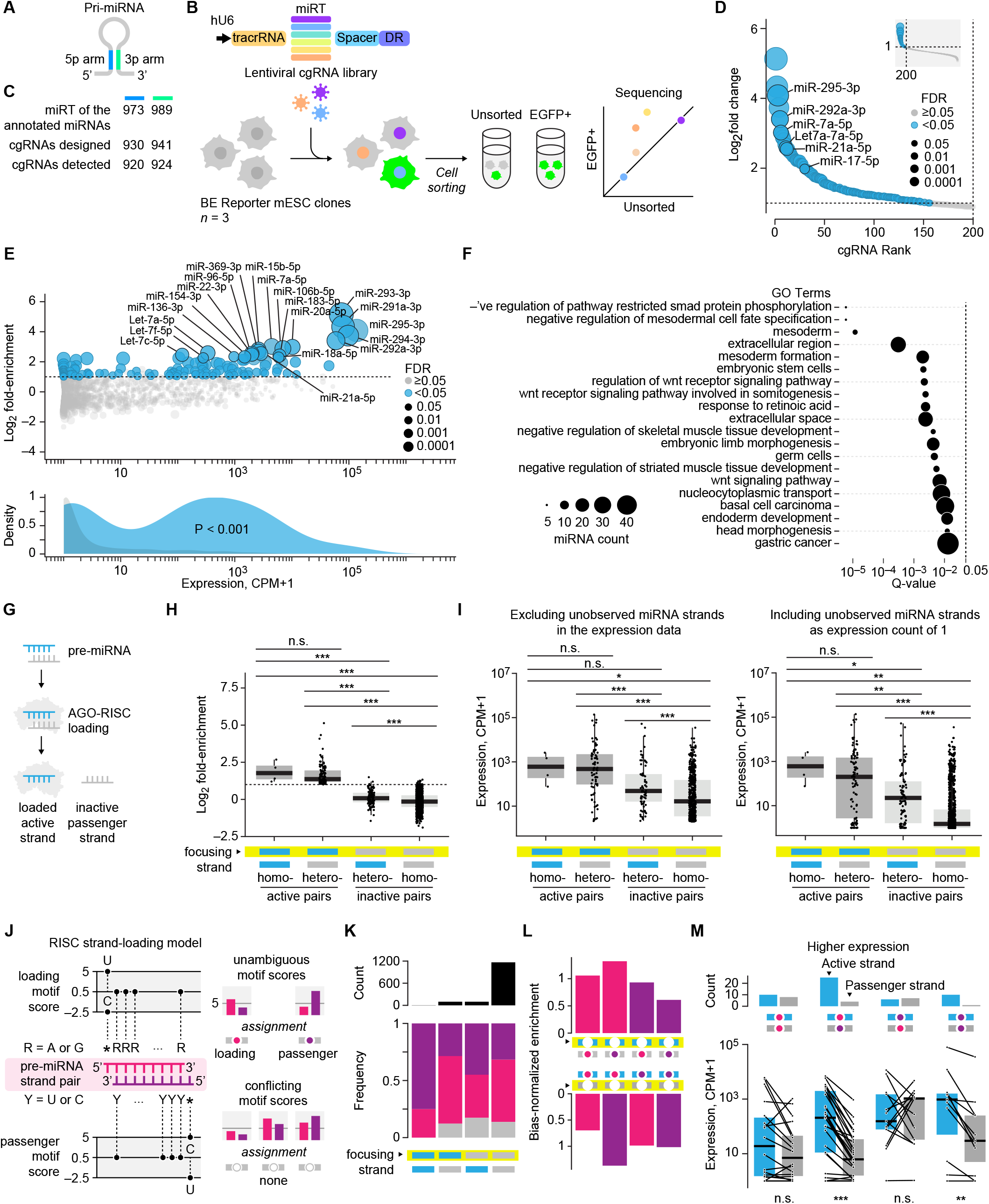
Genome-wide profiling of miRNA cleavage activity in mESCs. (A) Pri-miRNA hairpin encoding mature 5p and 3p miRNAs on opposing arms. (B) Workflow for pooled cgRNA screening of miRNA activity. A lentiviral library of cgRNAs carrying mouse miRNA target sequences was introduced into three independently derived clonal mESC reporter lines (Clones 3, 7, and 10). EGFP+ cells were isolated by flow cytometry and cgRNA abundance in the EGFP+ and unsorted populations was quantified by amplicon sequencing. (C) Numbers of annotated mouse 5p and 3p miRNAs, cgRNAs represented in the library, and cgRNAs detected across the unsorted populations. (D) Ranked mean log_2_ fold enrichment of cgRNAs in EGFP+ relative to unsorted cells. Hits were defined by log_2_ fold enrichment ≥ 1.0 and FDR < 0.05. Blue indicates FDR < 0.05, and point size reflects the FDR. (E) Relationship between cgRNA enrichment and corresponding miRNA abundance measured by AQ-seq. Significant hits are shown in blue. The lower panel compares the miRNA-abundance distributions of screen hits and non-hits; Mann–Whitney two-sided U-test. (F) Gene set enrichment analysis of miRNAs identified as screen hits. The dashed line denotes Q value = 0.05. (G) Schematic of active strand loading into RISC and exclusion of the inactive passenger strand. (H) cgRNA enrichment for, from left to right, active strands from homoactive pairs, active strands from heteroactive pairs, inactive strands from heteroactive pairs, and inactive strands from homoinactive pairs. The dashed line denotes log_2_ fold enrichment = 1.0. Mann–Whitney two-sided U-test. (I) AQ-seq abundance for the same strand categories, either excluding undetected miRNA strands or assigning them an expression count of 1. Mann–Whitney two-sided U-test. (J) Simplified sequence heuristic for assigning loading and passenger sequence motifs. R, A or G; Y, C or U. (K) Numbers and frequencies of loading, passenger, and unassigned motif classifications across the four strand-activity categories shown in H and I. (L) Conditional motif enrichment within heteroactive pairs. The focusing strand is outlined in yellow. Bars show the bias-normalized enrichment of loading and passenger motif assignments for the focusing strand, conditioned on the activity and motif assignment of the opposite strand. The upper row focuses on the active strand, whereas the lower row focuses on the inactive strand. (M) Paired AQ-seq abundances of active and inactive strands from heteroactive pairs, grouped by their combinations of loading and passenger motif assignments. Lines connect strands derived from the same pre-miRNA, and the upper bars summarize the numbers of pairs in which the active or inactive strand exhibited higher expression. Wilcoxon signed-rank test. n.s., not significant; \**P* < 0.05; \*\**P* < 0.01; \*\*\**P* < 0.001.

**Figure 4.**
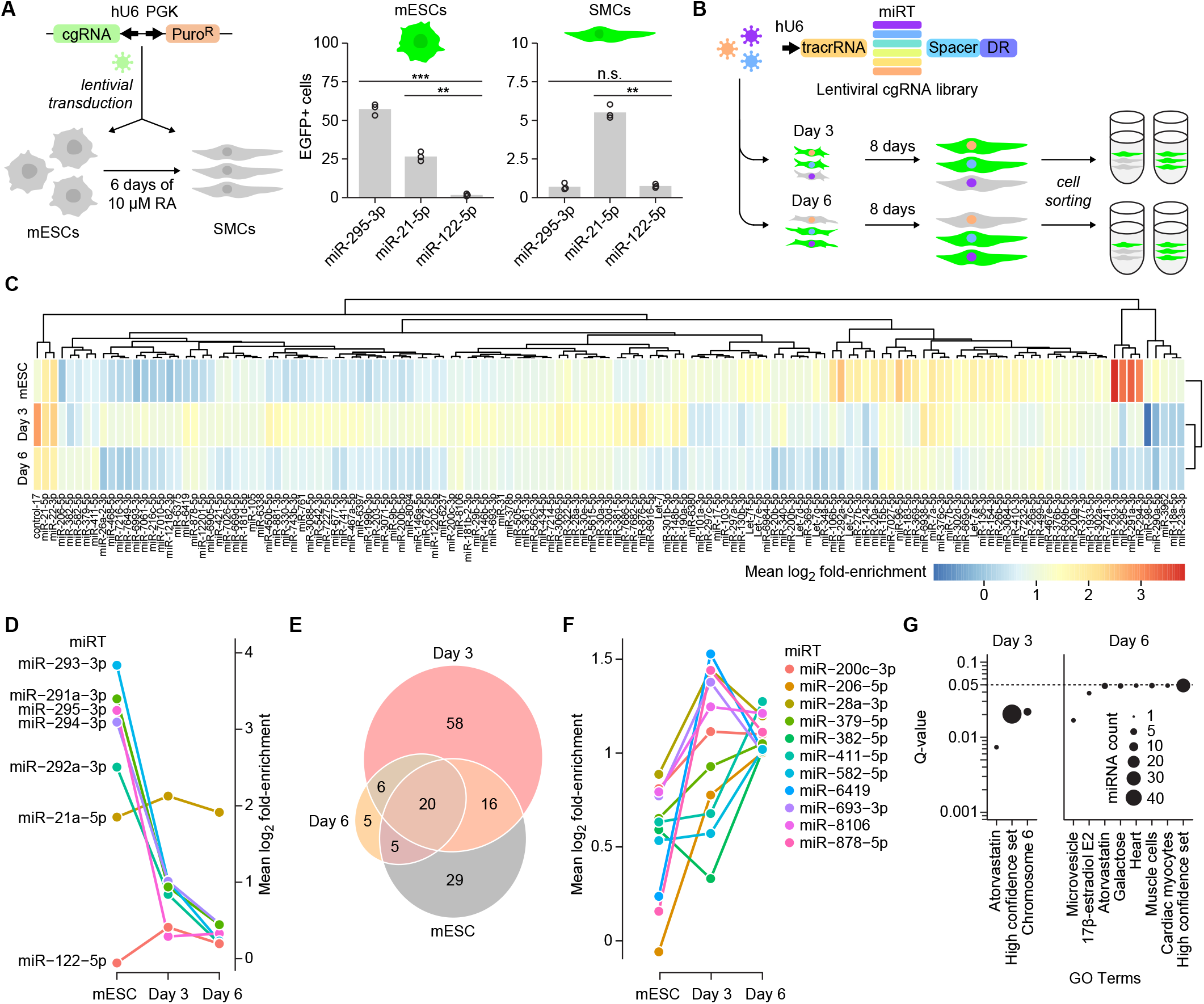
Genome-wide recording of miRNA activity during smooth muscle cell differentiation. (A) miRNA-dependent base editing in mESCs and smooth muscle cells (SMCs). Toolkit mESCs harboring the C→T base-editing reporter and intronic Target-AIDmax were maintained in the mESC state or differentiated for 6 days with 10 μM all-trans retinoic acid (RA) and then transduced with cgRNAs targeting miR-295-3p, miR-21-5p, or the negative-control miR-122-5p. The percentage of EGFP+ cells was quantified by flow cytometry. Welch’s t-test was used to compare the miR-295-3p- and miR-21-5p-targeted cgRNAs with the miR-122-5p-targeted control within each cell state (n = 3 replicates). (B) Workflow for genome-wide cgRNA screening during SMC differentiation. Clone 7 reporter mESCs were transduced with the pooled mouse miRNA-targeted cgRNA library in the mESC state or on Day 3 or Day 6 of differentiation. EGFP+ cells were isolated 8 days after transduction, and cgRNA abundance in the EGFP+ and unsorted populations was quantified by amplicon sequencing (n = 3 replicates per state). (C) Heatmap of mean log_2_ fold enrichment for the union of miRNA targets identified as hits in the mESC, Day 3, and Day 6 screens. Hits were defined by log_2_ fold enrichment > 1.0 and FDR < 0.05, and miRNA targets were ordered by hierarchical clustering. (D) Temporal enrichment profiles of selected mESC-associated miRNAs, including members of the miR-290–295 cluster, together with the broadly active miR-21a-5p and the negative-control miR-122-5p. (E) Overlap of miRNA hits identified across the mESC, Day 3, and Day 6 screens. (F) Temporal enrichment profiles of miRNAs identified as differentiation-specific hits at Day 3 and/or Day 6 but not in the mESC state. (G) Gene set enrichment analysis of miRNAs identified as hits at Day 3 and Day 6. Bubble size represents the number of miRNAs associated with each term, and the dashed line denotes Q = 0.05. n.s., not significant; \**P* < 0.05; \*\**P* < 0.01; \*\*\**P* < 0.001.

**Figure 5.**
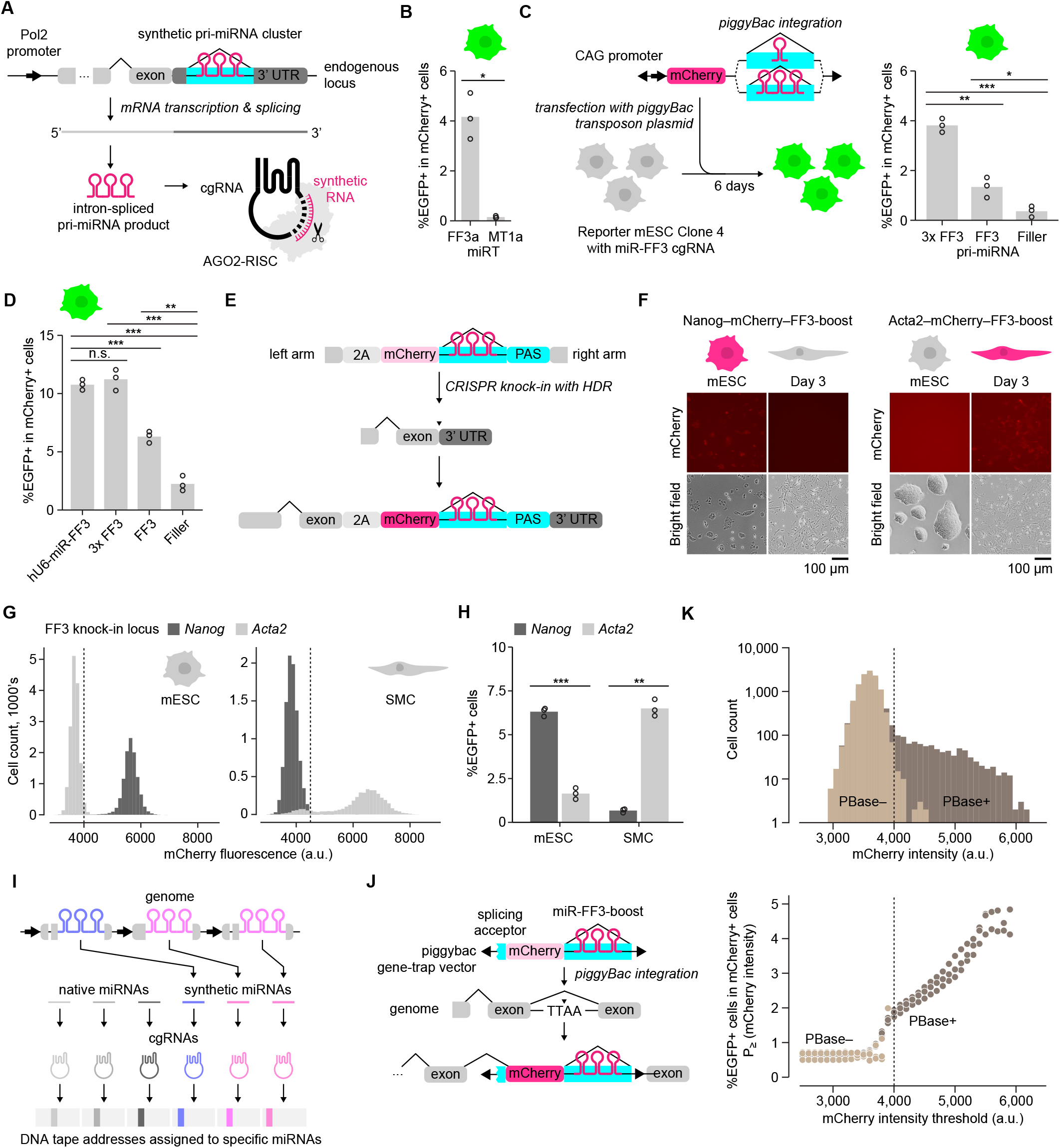
Recording gene expression through cgRNA-mediated genome editing. (A) Strategy for coupling Pol2 transcription to cgRNA activation through an intron-encoded synthetic pri-miRNA. Transcription and splicing release the pri-miRNA, which is processed into a mature synthetic miRNA that activates its cognate cgRNA. (B) Activation of miR-FF3-targeted cgRNA by hU6-driven pri-miR-FF3 in HeLa reporter cells. A cgRNA targeted by a non-expressed control miRNA (miR-MT1) was used to assess target specificity. Welch’s t-test; n = 3 replicates. (C) cgRNA activation by CAG-driven Pol2 transcripts containing a synthetic intron encoding a single pri-miR-FF3, a tandem cluster of three pri-miR-FF3 copies (pri-miR-FF3-boost), or a filler sequence in Clone 4 mESC reporter cells. Welch’s t-test for the indicated comparisons; n = 3 replicates. (D) Comparison of hU6-driven pri-miR-FF3 with CAG-driven single and tandem pri-miR-FF3 constructs in polyclonal mESC reporter cells. Welch’s t-test for the indicated comparisons; n = 3 replicates. (E) CRISPR–Cas9-mediated knock-in of a 2A–mCherry–pri-miR-FF3-boost cassette at the 3′ end of the endogenous *Nanog* or *Acta2* coding sequence. (F) Representative fluorescence and bright-field micrographs of *Nanog*–mCherry–FF3-boost and *Acta2*–mCherry–FF3-boost cells in the mESC state and on day 3 of SMC differentiation. Scale bars, 100 μm. (G) Flow-cytometric distributions of mCherry expression in the knock-in cells shown in F. Dashed lines indicate the thresholds used to define mCherry+ cells. (H) cgRNA-mediated recording of endogenous *Nanog* and *Acta2* expression in the mESC state and on day 6 of SMC differentiation. The percentage of EGFP+ cells was quantified by flow cytometry. Welch’s t-test was used to compare the two knock-in lines within each cell state; n = 3 replicates. (I) Conceptual framework for multiplexed recording of native miRNA activities and endogenous gene-expression states using synthetic pri-miRNA–cgRNA pairs coupled to distinct DNA memory addresses. (J) Gene-trap strategy for coupling expression of randomly targeted endogenous genes to mCherry and pri-miR-FF3-boost production. Following piggyBac integration in the appropriate orientation within an actively transcribed gene body, the payload is expressed through splicing to an upstream endogenous exon. (K) Transcriptional hitchhiking and associated cgRNA-mediated base editing in Clone 4 reporter mESCs transfected with the piggyBac gene-trap vector in the presence or absence of piggyBac transposase. Top, mCherry fluorescence distributions. Bottom, percentage of EGFP+ cells among populations exceeding the indicated mCherry fluorescence thresholds; points represent n = 3 replicates. n.s., not significant; \**P* < 0.05; \*\**P* < 0.01; \*\*\**P* < 0.001.

**Figure 6.**
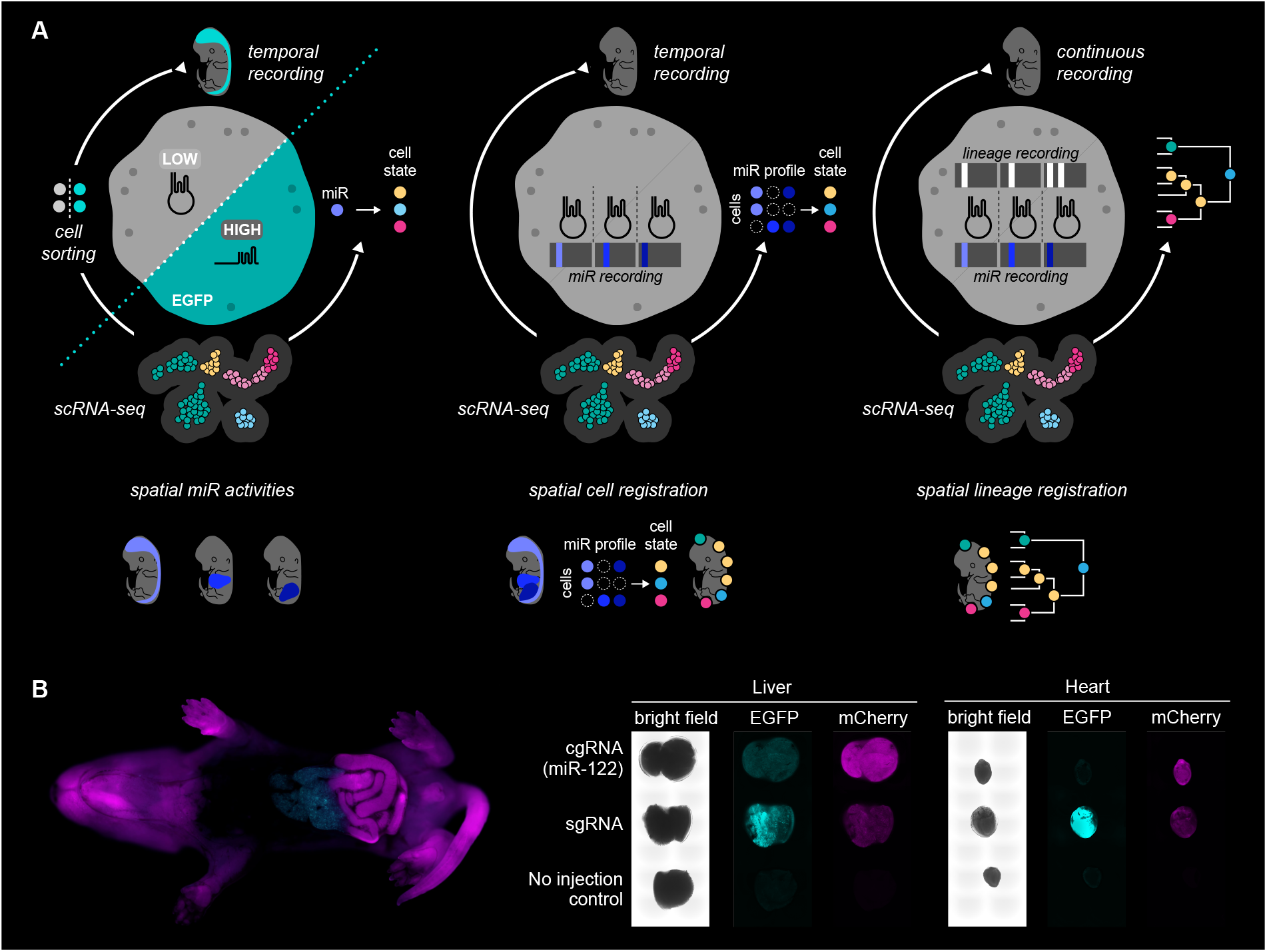
Prospective applications and a preliminary *in vivo* validation of cgRNA-based molecular recording. (A) Prospective applications of cgRNA in single-cell, spatial, and lineage-resolved molecular recording. Top left, temporal recording of an individual miRNA activity, followed by reporter-based cell isolation and scRNA-seq to associate functional miRNA activity with cellular state. Bottom left, organism-scale spatial mapping of individual miRNA activities. Top middle, multiplexed recording of multiple miRNA activities into distinct DNA memory addresses to generate cell-state-associated miRNA activity profiles. Bottom middle, spatial registration of single-cell states using recorded miRNA activity profiles as molecular landmarks. Top right, continuous and simultaneous recording of miRNA activities and cellular lineages. Bottom right, spatial reconstruction of lineage relationships together with recorded molecular-state histories. (B) miR-122 activity-dependent genome editing in mice. Mouse zygotes were pronuclearly injected with the C→T EGFP base-editing reporter, intronic Target-AIDmax, and either sgRNA or miR-122-targeted cgRNA, followed by embryo transfer. Left, representative whole-body fluorescence image of a postnatal mouse receiving the miR-122-targeted cgRNA. Right, representative bright-field, EGFP, and mCherry images of liver and heart tissues from sgRNA-injected, miR-122-targeted cgRNA-injected, and uninjected animals.

## RESULTS

### Development and optimization of cgRNA

We first designed three versions of cgRNA to test for miRNA activity-dependent genome editing. cgRNA V1 utilized the natural helical RNA stem formed between crRNA and tracrRNA of the *Streptococcus pyogenes* Cas9 (SpCas9) guide RNA for circularization (**Figure S1C**). cgRNA V2 instead utilized the second stem of the sgRNA scaffold for circularization, with the stem sealed by ribozyme-mediated splicing^56^ (**Figure S1D**) to potentially create a more stable structure. cgRNA V3 extended the first stem of the sgRNA scaffold with a small synthetic helical structure^57^ (**Figure S1E**). Both cgRNA V2 and V3 were designed to increase the cuffed structure’s stability and genome editing efficacy. RNA circularization and the synthetic minihelix have also been known to facilitate the shuttling of RNA polymerase III (Pol3)-transcribed RNAs to the cytosol, where miRNA activity typically occurs^58^. We hypothesized that these modifications would improve editing efficiency by cgRNA because cleavage of cgRNA by miRNA likely occurs in the cytosol followed by translocation of the cleaved product into the nucleus after loading into cytosolic Cas9.

Throughout the study, we used a common reporter toolkit for each cell line, where a C→T base editing reporter and Target-AID with human codon optimization (Target-AIDmax)^59,60^, both of which were previously established by our group, were integrated via piggyBac transposon (**Figure 1C**). In this reporter system, the start codon of an enhanced green fluorescent protein (EGFP)-encoding gene is mutated to GTG, resulting in no protein expression. Protein production of EGFP can be restored by the editing of the mutated start codon from <u>G</u>TG→<u>A</u>TG by C→T base editing of its antisense strand with a functional guide RNA (**Figure 1D**). In the context of cgRNA, this can be used to report the cleavage activity of a target miRNA. After establishing a toolkit cell line, we introduced cgRNA with either mCherry or the puromycin-resistance gene via lentiviral transduction at an infection rate of 0.1 or lower, ensuring single-copy integration (**Figure 1E**).

We compared the miRNA activity-dependent genome editing efficiencies of the three versions of cgRNA with crRNA:tracrRNA and MICR. We first screened an efficient guide RNA spacer target sequence (US1) in a HeLa reporter cell line (**Figure S1F**) and lentivirally transduced different guide RNA variants with an on-target (OT) or non-target (NT) spacer and an mCherry fluorescent protein gene at an infection rate of <0.1 (**Figure 1E**). As in the previous MICR study^61^, all the cgRNA versions and MICR were designed to be activated by hsa/mmu-miR-122-5p (miR-122), which is not expressed (absent) in HeLa cells. The toolkit HeLa cells with each guide RNA variant were then transfected with a synthesized on-target (OT) miR-122 miRNA mimic or a non-target (NT) miRNA mimic (**Figure 1F**). Activation of the C→T base editing reporter was then observed using fluorescent microscopy imaging (**Figure 1G**) and flow cytometry analysis (**Figure 1H**). EGFP activation was observed in both the control sgRNA (**Figure S1A**) and crRNA:tracrRNA, regardless of the miRNA mimic input. cgRNA V1 and V3, and MICR showed EGFP activation only in response to the OT miRNA mimic. Unexpectedly, cgRNA V2 did not exhibit input miRNA mimic specificity. Of those guide RNAs that activated EGFP in response to the OT miRNA mimic only, cgRNA V3 and MICR showed the two highest OT activations (5.56% and 6.46% EGFP+ cells, respectively), whereas cgRNA V1 showed the lowest activation (2.81% EGFP+ cells). We also observed that cgRNA and MICR genome-editing efficiencies were substantially lower than that of sgRNA (80.13% EGFP+ cells) but comparable to that of crRNA:tracrRNA (6.63% EGFP+ cells). This was expected given that cgRNA was miRNA activity-gated and not a concern (we present several pieces of supporting evidence below that demonstrate the practicality of cgRNA).

To examine how these different miRNA activity-dependent guide RNAs respond to different abundances of a target miRNA and subsequently confer genome editing, we treated cgRNA V1–V3 and MICR cells with increasing doses of OT miRNA mimic and analyzed EGFP activation by flow cytometry (**Figure 1I**). Consistent with the previous result, while cgRNA V2 showed constitutive EGFP activation independent of the miRNA mimic dosage, cgRNA V1 and V3, and MICR showed increasing levels of EGFP activation along with the miRNA mimic dosage. While cgRNA V1 showed the lowest response, cgRNA V3 and MICR showed similar response patterns, where cgRNA V3 showed significantly higher activities in a high dosage range of 0.312 to 5.000 nM. Accordingly, cgRNA V3 performed comparably or better than the current state-of-the-art miRNA activity-gated guide RNA.

We additionally designed cgRNA V4 and V5, alongside V3, to investigate if there are better potential cgRNA configurations. cgRNA V4 employed ribozyme-mediated circularization at the first stem position of the sgRNA (**Figure S1G**). We constructed V3 and V4 cgRNAs to be targeted by hsa/mmu-miR-21-5p (miR-21) (present in HeLa cells)^61^ and miR-122 (absent), introduced them into toolkit HeLa cells via lentiviral transduction (infection rate < 0.1), and observed that they performed similarly in response to miR-21 only. cgRNA V5 was permutated at the second stem position of sgRNA, like cgRNA V2, and the stem was extended with the synthetic minihelix from cgRNA V3. We constructed V3 and V5 cgRNAs targeted by mmu-miR-295-3p (miR-295, present in mESCs) and miR-122 (absent), lentivirally transduced them (infection rate < 0.1) to mESCs harboring the reporter toolkit, and observed that cgRNA V5 diminished miRNA targeting specificity (**Figure S1H**).

We also tested whether increasing the loop size of cgRNA could enhance accessibility to the miRT but hypothesized that this would be at the expense of raising background editing activity by reducing the structural distortion caused by the circularization of the guide RNA. However, we observed the opposite effect in HeLa cells: loop expansions with a 6-to 18-nt linker diminished the editing activity of cgRNA targeted by miR-21 and did not elevate unexpected editing by cgRNAs targeted by a negative control miRNA (absent in HeLa) as a function of the linker size (**Figure S1I**). (Note that we did not test longer linker sizes in this assay, but in the following biochemical assay in **Figure 2**, we found that linkers of 20 nt or greater could restore genome editing activity in the absence of miRNA cleavage *in vitro.*) Taken together, we decided to use cgRNA V3 for the remainder of this study. Hereafter, all references to cgRNA refer to cgRNA V3.

cgRNA could also be coupled with the commonly used wild-type SpCas9 and the Cas9 variant of *Staphylococcus aureus* (SaCas9), in addition to C→T base editing. We engineered a new set of HeLa toolkit cell lines, harboring a piggyBac transposon-integrated SpCas9 or SaCas9 and lentivirally integrated bicistronic mCherry-IRES-EGFP reporter and miR-122-responsive cgRNA with a spacer sequence targeting the EGFP coding sequence (**Figures 1J** and **S1J**). For SaCas9, we engineered its cgRNA using the same design principles as cgRNA V3. In contrast to the C→T base-editing reporter system, successful genome editing with these wild-type Cas9 variants was expected to stochastically induce frameshift mutations, leading to loss of the EGFP signal. Transfection of OT and NT miRNA mimics into the toolkit cells showed miRNA mimic-dependent genome editing, with SpCas9 showing higher editing activity than SaCas9 (8.68% versus 1.35% EGFP+ cells). Accordingly, we demonstrated the broad potential of cgRNA with base editing and wild-type Cas9, at least among Class II Type II CRISPR-Cas systems.

### Involvement of the miRNA cleavage pathway

AGO2, a member of the Argonaute protein family, is the catalytic component of the RISC and mediates target RNA cleavage in miRNA- and small interference (si)RNA-guided silencing. AGO2 abundance has been observed to be a rate-limiting factor in the cleavage pathway of siRNA^62^. Thus, we expected that cellular AGO2 might be of lower abundance than cellular miRNA and that AGO2 overexpression could access this excess miRNA pool and so might increase miRNA-dependent cgRNA activity, supporting the involvement of the miRNA cleavage pathway in cgRNA activation by a target miRNA.

We further engineered the toolkit reporter HeLa cell line (**Figure 1C**) by piggyBac transposon integration of an additional overexpression cassette encoding human codon-optimized AGO2 (hereafter simply referred to as AGO2), control firefly luciferase (Luc2), or AGO2 with a deletion of the catalytic PIWI domain^63^ (AGO2 ΔPIWI). These overexpression HeLa cell lines were then transduced with either a reporter targeting sgRNA or a cgRNA targeted by miR-122 (absent in HeLa) and a puromycin-resistance gene at a high infection rate of 0.5 or more. The established cell lines were then transfected with an OT miR-122 mimic or an NT miRNA mimic (**Figure 1K**). Flow-cytometry analysis showed that EGFP activation by sgRNA did not largely differ across AGO2-, Luc2-, and AGO2 ΔPIWI-overexpression cell lines (89.23–86.08% EGFP+ and 93.07–91.27% EGFP+ cells in the OT and NT miRNA mimic conditions, respectively). In contrast, cgRNA with the OT miR-122 mimic showed the most significant elevation of genome editing activity in the AGO2-overexpression cells (26.31% EGFP+ cells) compared to those of the control conditions (18.27% and 17.31% EGFP+ cells for Luc2 and hAGO2 ΔPIWI, respectively). cgRNA with the NT miRNA mimic condition did not show genome editing activity across the overexpression conditions (0.91–1.82%).

Next, to examine whether AGO2 overexpression can boost cgRNA activity triggered by endogenously expressed miRNA, we lentivirally transduced cgRNAs for miR-21 (present in HeLa) and miR-122 (absent in HeLa) with a puromycin-resistance gene to the overexpression cell lines (infection rate > 0.5). Similar to the miRNA mimic experiments, flow cytometry analysis showed that EGFP activation by sgRNA did not largely differ across AGO2-, Luc2-, and AGO2 ΔPIWI-overexpression cell lines (84.52%-81.90% EGFP+) (**Figure S1K**). In contrast, cgRNA for miR-21 showed the most significant elevation of genome editing activity in the AGO2 overexpression cells (27.34% EGFP+ cells) compared to those of the control conditions (13.64% and 10.94% EGFP+ cells for Luc2 and AGO2 ΔPIWI, respectively). The negative control cgRNA for miR-122 did not show genome editing activity across the overexpression conditions (0.58–1.01%). Overall, these results support the involvement of AGO2 in the miRNA-dependent activation of cgRNA. Furthermore, while the overexpression of AGO2 has previously only been tested in the context of siRNA activity^62^, our results suggest that overall cellular miRNA activity can also be modulated by AGO2 expression levels.

Aside from AGO2 abundance, another factor that affects target cleavage efficiencies is the extent of sequence complementarity and accessibility between the miRNA and its target. This ranges from the length of the target sequence^31,64^, positional mismatches^31,55^, and even adjacent structured RNA regions^55^. It is known that mismatches near the cleavage site (11^th^–12^th^ nt from the 3′ end) of the miRT often reduce target RNA cleavage by AGO2. When programming a mismatch at the 11^th^ nt position of the miRT for hsa/mmu-let-7a-5p (present in mESCs), its genome editing activity in mESCs was diminished compared to that of the fully complementary target sequence (**Figure S1L**).

In contrast, previous studies have also observed that perfect complementarity between a miRNA and its target does not necessarily maximize miRNA cleavage kinetics since it reduces the rate of decoupling of miRNA from the target-bound AGO2–RISC complex^31,64^. Primarily, 2–3 nt mismatches at the 3′ end of miRNA (non-seed region; 5′ end of miRT) increase the silencing activity, whereas 5 nt or greater mismatches can diminish or abolish the miRNA activity in mammalian cell culture^64^. To test whether the same phenomenon can be recapitulated in the context of cgRNA, we progressively truncated the 5′ end of the miRT of cgRNAs targeted by miR-122. Toolkit HeLa cells were transduced with cgRNA truncation variants with an mCherry-coding gene (infection rate < 0.1) and then transfected with an OT miR122 mimic or NT miRNA mimic. The 3-nt miRT truncation resulted in a modest increase in genome editing (14.51% EGFP+ cells) compared to the fully complementary miRT sequence (11.10% EGFP+ cells), while the 5-nt and 7-nt truncations resulted in a reduction (7.25% EGFP+ cells) and loss (0.15% EGFP+ cells) of activity, respectively (**Figure 1L**).

Collectively, these results support the view that the activation of cgRNA in response to its target miRNA involves the AGO2–RISC-mediated miRNA cleavage pathway and provide potential design considerations for further optimizing the system’s sensitivity beyond the cgRNA structure.

### Quantitative measurement of cleavage activities across different miRNAs

Next, we sought to determine whether genome editing by cgRNA can universally report quantitative miRNA cleavage activities across different miRNA species. Prior to this analysis, we tested whether genome editing activity by cgRNA was sensitive to the cgRNA copy number per cell. We transduced toolkit HeLa cells with different doses of lentiviruses encoding cgRNA targeted by miR-21 or by miR-122, along with mCherry. The percent mCherry+ cells was used to approximate the viral infection rate (**Figure S1M**). We then measured the percentage of EGFP+ cells among mCherry+ cells to estimate genome editing activity. For the cgRNA targeted by miR-21, we observed a consistent EGFP+ rate, averaging 12.74%, across viral infection rates below 0.25, which, according to Poisson statistics, indicate single-copy genomic integration of the payload in infected cells. EGFP+ rate then increased along with the viral infection rate up to 0.41, indicating that for higher copy numbers of cgRNA substrates, there is a response to the pool of active RISC-loaded miRNA complexes in the cell. On the other hand, the control cgRNA targeted by miR-122 showed almost no background activation (2.35% EGFP+ cells at the highest multiplicity of infection). We concluded that a single cgRNA expression unit per cell is sufficient for reporting the quantitative activity of a target miRNA in a cell population, but sensitivity can be dialed up by increasing the cgRNA dosage per cell.

Given these results, we constructed cgRNAs targeted by seven arbitrarily selected miRNAs that are conserved across human and mouse but have been reported to be differentially expressed in HeLa cells and mESCs, as well as miR-295 and mmu-miR-293-3p (miR-293) that are specific to the mouse genome. We lentivirally transduced these cgRNAs into both the toolkit HeLa cells and mESCs with single-copy integration (infection rate < 0.1) and measured the percent EGFP+ of the infected cells (**Figures S2A–S2D**). Separately, we performed AQ-Seq^65^, a bias-minimized small RNA sequencing, and quantified these miRNA expression levels in both toolkit cell lines (**Figures S2E–S2H**). The activation rates of the base editing reporter by different cgRNAs were significantly correlated with the expression levels of their corresponding miRNAs in both HeLa cells and mESCs (**Figure 1M**). In mESCs, hsa/mmu-miR-9-5p (miR-9) and hsa/mmu-miR-124-3p (miR-124) demonstrated lower levels of the reporter activation than hsa/mmu-miR-7-5p (miR-7), miR-21, and hsa/mmu-Let-7a-5p (Let-7a), albeit they all showed similar expression levels. These discrepancies between miRNA expression and genome editing level might stem from other cellular factors regulating miRNA activities, as exemplified by miR-124, which is expressed in mESCs but has been observed to be inactive in the mESC state before neural differentiation^66^. At the same time, the overall correlation between the cgRNA activity and its corresponding miRNA expression level in two different cell types suggests that cgRNA activity reflects the miRNA cleavage activity in the cell, and expression might be a major factor in this relationship. Interestingly, in the expression-activity correlation plots of both HeLa cells and mESCs, the miRNA expression difference of around two orders of magnitude could be covered by a cgRNA activity difference of only one order of magnitude. This may reflect the miRNA recycling effect, in which a single miRNA molecule cleaves many target RNA molecules, consistent with the results of the miRT truncation experiments, which showed that a potential fast target releaser yields higher genome editing (**Figure 1L**). Altogether, this dynamic range compression from expression to genome editing activity demonstrates that cgRNA is sensitive enough to respond to lower-abundance miRNAs.

In addition to HeLa cells and mESCs, we also constructed a toolkit human induced pluripotent stem cells (hiPSCs) harboring piggyBac transposon integrated C→T base editing reporter and Target-AIDmax (**Figure 1C**). We lentivirally delivered cgRNAs targeted by hsa-miR-302a-3p and hsa-miR-302b-3p (present in hiPSC^67^) and those targeted by non-expressed hsa/mmu-miR-1a-3p and hsa/mmu-miR-133a-3p (infection rate < 0.1), and found that despite overall lower activity levels, there were higher genome editing activities for miR-302a and b compared to miR-1a or miR-133a (**Figure S2I**).

### Multilayered inhibition of CRISPR–Cas9 genome editing kinetics by cgRNA

To investigate how cgRNA regulates Cas9 cleavage activity, we performed biochemical and structural analyses. We first performed an *in vitro* DNA cleavage assay using purified SpCas9 and *in vitro*-transcribed guide RNA variants, which include sgRNA, cgRNA, and a mimic of AGO2-mediated cleaved cgRNA (a mixture of crRNA and tracrRNA with extended sequences for expected cleaved miRT halves) for targeting of a linearized plasmid DNA containing a guide RNA target sequence with a TGG protospacer adjacent motif (PAM) (**Figure 2A** and **Table S1**). We did not observe detectable DNA cleavage activity of the linearized plasmid DNA by the SpCas9–cgRNA complex. In contrast, both the SpCas9–cleaved-cgRNA-mimic complex and the SpCas9–sgRNA complex successfully cleaved the target DNA, although the SpCas9–cleaved-cgRNA-mimic complex exhibited lower cleavage efficiency than the SpCas9–sgRNA complex, presumably because of the imperfect assembly of crRNA and tracrRNA. Additionally, a competitive electrophoretic mobility shift assay (EMSA) using a mixture of sgRNA and cgRNA showed that both Cy5-labeled cgRNA and sgRNA binding to SpCas9 were markedly reduced in the presence of increasing concentrations of a competitive unlabeled sgRNA (**Figures 2B** and **S3A–C**). On the other hand, assays with a competitive cgRNA showed minimal reduction in binding affinity. These findings therefore indicate that the cgRNA architecture suppresses SpCas9-mediated DNA cleavage, at least in part, by impairing the binding affinity of SpCas9 for the guide RNA.

To further elucidate the molecular mechanism underlying cgRNA-mediated Cas9 regulation, we attempted cryo-EM analysis of the SpCas9–cgRNA–target DNA ternary complex by mixing wild-type SpCas9 with cgRNA (164 bp) and its corresponding target DNA (39 bp) (**Figures 2C** and **S3D–K**). However, despite the ternary complex assembly conditions, our initial 3D classification revealed that the majority of sorted particles corresponded to the SpCas9–cgRNA binary complex (**Figures 2C** and **S3D**). Conversely, previous structural studies using the canonical sgRNA only identified the ternary complex from their datasets^68–72^. Through iterative 3D classification and refinement, we determined the SpCas9– cgRNA binary complex structure at an overall resolution of 3.3 Å (**Figures 2D** and **S3G, S3H, S3I**). The overall architecture of SpCas9–cgRNA binary complex closely resembled that of the previously reported SpCas9–sgRNA binary complex in an open conformation prepared for linear DNA binding (**Figures S4A–S4C**)^73,74^. The cgRNA scaffold showed a canonical sgRNA-like conformation comprising stem loops 1–3 (with stem loop 1 disordered) and a repeat:anti-repeat duplex that included an extended mini-helix segment (**Figures S4A, S4D,** and **S4E**). The miRT region was completely disordered, likely reflecting its intrinsic flexibility. Notably, we observed ambiguous density corresponding to the cgRNA spacer segment between the REC1 and PI domains, where initiation of the spacer–target heteroduplex formation occurs. These structural observations suggest that the cgRNA spacer segment is located in a region proximal to the REC1 and PI domains, thereby preventing initial target DNA engagement and subsequent formation of the SpCas9–cgRNA–target DNA ternary complex. Consistent with these structural observations, an EMSA of Cy5-labeled target DNA using Cas9 in complex with sgRNA or cgRNA demonstrated that cgRNA markedly impairs the target DNA-binding ability of SpCas9 (**Figure S4F**). These results indicate that the cgRNA architecture further suppresses SpCas9-mediated DNA cleavage by interfering with the formation of the SpCas9–cgRNA–target DNA ternary complex through positional constraint of the cgRNA spacer segment between the REC1 and PI domains. The iterative 3D classification, however, also identified a minor population of particles corresponding to the SpCas9–cgRNA–target DNA ternary complex and determined its structure at an overall resolution of 3.1 Å (**Figures 2E, 2F, S3G, S3J, S3K, S4G,** and **S4H**). In this structure, the SpCas9–cgRNA complex failed to form a full R-loop comprising the canonical 20-bp spacer–target heteroduplex, but instead adopted a partial R-loop, in which nucleotides rA118–rA127 of the cgRNA spacer base pair with nucleotides dT10–dT1 in the intact target strand (TS) to form a 10-bp heteroduplex (**Figures 2G, 2H,** and **S4G**). Furthermore, SpCas9 protein adopted an inactive conformation^72^, with the HNH nuclease domain positioned approximately 42 Å away from the TS cleavage site (**Figures 2I** and **S4H–S4J**). Collectively, these structural observations suggest that even when the SpCas9–cgRNA complex engages the target DNA, it does not form the full R-loop conformation, thereby preventing SpCas9 nuclease activation. We hypothesized that this truncated R-loop is likely due to the physical tension imposed by the cuffed architecture of cgRNA. To test this hypothesis, we performed *in vitro* cleavage assays using cgRNA variants designed to enhance spacer mobility, with linkers of varying lengths between the spacer and the miRT region (**Figure 2J**). Although we did not see background base editing activity by cgRNA in HeLa cells in the absence of a target miRNA for the linker size of up to 18 nt (**Figure S1I**), we found that the cleavage activity of SpCas9 was restored with a linker size of 20 nt or greater in this *in vitro* experiment. These results support our model of cgRNA-mediated regulation, where the cuffed architecture restricts spacer mobility, thereby hindering R-loop formation and effectively locking Cas9 in catalytically inactive conformations.

Taken together, we show that applying a cuffed structure to the guide RNA may impair CRISPR genome-editing kinetics in three sequential modes (**Figure 2K**). The competitive EMSA assay for the binding of guide RNA variants to SpCas9 suggests that the cuffed structure diminishes the binding kinetics of guide RNA to Cas9. The enrichment of the SpCas9–cgRNA binary complex in the cryo-EM classification, its spacer positioning in the REC1 and PI domain-proximal region, and the EMSA assay for its binding to DNA collectively show that cgRNA also constrains access to the target DNA. Furthermore, even after target DNA binding, the SpCas9–cgRNA–target DNA ternary complex and the loop extension assay demonstrate further inhibition of Cas9 cleavage through incomplete R-loop formation.

### Genome-wide characterization of miRNA cleavage activities in mESCs

miRNA biogenesis begins with transcription of primary miRNA (pri-miRNA) transcripts in the nucleus^75–77^. Pri-miRNAs form characteristic stem–loop structures, in which the mature 5p and 3p miRNAs are encoded on opposite arms of the hairpin (**Figure 3A**). The Microprocessor complex, composed of a single RNase III enzyme Drosha and two copies of the RNA-binding protein DGCR8, recognizes and cleaves pri-miRNA hairpins to generate ∼55–70 nt precursor miRNAs (pre-miRNAs). These pre-miRNAs are then exported to the cytoplasm by Exportin-5 and further processed by Dicer into ∼22-nt miRNA duplexes corresponding to the 5p and 3p strands. One strand (the guide strand) is preferentially loaded into an Argonaute protein (AGO1–4) to form the RISC, while the non-loaded strand (passenger strand) is ejected from the complex and shuttled for degradation. As discussed in the previous section, the Argonaute protein isoform AGO2 possesses endonucleolytic (“slicer”) activity, and a miRNA can directly cleave its target mRNA when loaded into the AGO2–RISC, provided there is nearly perfect sequence complementarity between the miRNA and its target.

Leveraging the capacity of cgRNA to estimate miRNA cleavage activity, we set up a pooled lentivirus assay to systematically measure the genome-wide miRNA cleavage activity profile of mESCs (**Figure 3B**). We first synthesized an oligo pool of 1,871 miRT sequences perfectly matching the sequences of annotated mouse miRNAs (930 and 941 miRTs for 935 5p- and 951 3p-miRNAs, respectively) and 54 negative control targets (**Figure 3C** and **Table S2**). The target oligo pool was cloned into a common cgRNA lentiviral plasmid backbone targeting the C→T base editing reporter to establish a cgRNA library. Some targets were excluded due to the presence of restriction enzyme sequences used for library cloning, or poly-thymidine stretches in their sequence that were expected to lead to termination of Pol3 RNA transcription.

For the genome-wide activity screen, we generated a new set of clonal reporter toolkit mESC lines for this screen and for subsequent experiments in which we recorded miRNA activities during cell differentiation. Prior to cell line establishment, we incorporated multiple endogenous mouse intron sequences into the Target-AIDmax gene, based on previous reports that incorporating intron sequences into transgene coding regions can prevent transgene silencing^78–81^ (**Figure S5A**). When intronic versions of Target-AIDmax were introduced into mESCs along with the C→T base editing reporter via piggyBac transposon integration, the Target-AIDmax genes with two and three introns showed higher expression levels than canonical Target-AIDmax in mESCs according to the bicistronic BFP reporter signal (**Figure S5B**). Although retrospective analysis of base editing efficiency did not show an improvement in the polyclonal mESC toolkit lines (**Figure S5C**), we had already proceeded to generate three clonal lines from the two-intron Target-AIDmax gene and utilized them for the subsequent miRNA activity screen (Clones 3, 7, and 10). These three clones were selected based on qualitative observations of similar mESC morphology and similar growth rates.

Following virus packaging, the three mESC lines were transduced with the cgRNA library at an infection rate of 0.1 or less, ensuring a single cgRNA integration per cell, cultured for six days, and sorted for EGFP+ cells using flow cytometry (**Figure S5D**). The miRT sequences were then amplified by PCR from the extracted genomic DNA of the EGFP+ and unsorted cell populations and read out by deep sequencing (**Figures S5E**). Finally, the enrichment of every cgRNA in the genome-edited population over the control population was calculated using DESEQ2^82^. Among the synthesized miRTs, 1,844 (920 5p- and 924 3p-miRTs, respectively) were commonly detected in the control unsorted cell populations across the three mESC lines (**Figure 3C**). Active miRNA hits were called based on targets that showed enrichment above an arbitrary enrichment threshold of 2.0-fold compared to the control population and a false discovery rate (FDR) of less than 5%. Based on these criteria, our screen captured 139 miRNA hits (**Figure 3D**), with 3 of the 54 negative control targets being detected as hits and 4 being undetected (**Figure S5F**).

We confirmed that the cgRNA enrichment signals in the pooled assay were significantly correlated with the percent EGFP+ cell values obtained for individual cgRNAs that were transduced independently to mESCs (**Figure 1M** and **Figure S5G**). Additionally, cgRNAs that generated percent EGFP+ values of 2.0% or more in individual assays were all identified as significant hits in the genome-wide screen. Mapping of genome-wide cgRNA editing activities to their corresponding miRNA expression levels showed that miRNA targets enriched in the genome-wide cgRNA screen had a statistically significant positive shift in distribution towards higher expression values, with a median count of 177.19 per million (CPM) compared to non-enriched miRNA targets with a median count of 1.42 (**Figure 3E**). Finally, we tested whether enriched targets could be linked to mESC-related functions using a miRNA-based gene set enrichment analysis^83^. We detected expected terms, such as embryonic stem cells, germ cells, Wnt signaling pathways^84^, negative regulation of pathway-restricted Smad protein phosphorylation^85^, as well as hits for negative regulation of differentiating phenotypes, such as mesoderm formation and skeletal and striated muscle (**Figure 3F**). Altogether, these results suggest that the enriched hits from the genome-wide cgRNA library screen reflect miRNA cleavage activities in mESCs.

It has been observed that one strand of a miRNA duplex, processed from pre-miRNAs by Dicer, is loaded into an Argonaute protein to form the RISC complex while the other strand is excluded for degradation^77^ (**Figure 3G**). Leveraging our comprehensive capture of the miRNA cleavage landscape in mESCs, we next investigated whether this previously described model for guide-strand selection from miRNA duplexes could also be recapitulated globally in our dataset.

106 of the 139 miRNA hits from the genome-wide screen had a miRNA counterpart originating from the same pre-miRNA hairpins, with a target also present in the cgRNA library (102 pairs in total). We categorized these paired targets into two groups, 4 homoactive pairs and 98 heteroactive pairs, depending on whether their miRNA strands were found to be active hits for cleavage (**Figure 3H**). We also defined the remaining 1,166 pairs in the library as homoinactive pairs. Comparing the activity levels (fold-enrichment of cgRNA abundance in the EGFP+ cell population over the control population) of the active strands of the homoactive pairs, the active and inactive strands of the heteroactive pairs, and the inactive strands of the homoinactive pairs, we confirmed that the cleavage activities of all active strands were overall significantly higher than those of all inactive strands and that the inactive strand activities of the heteroactive pairs were comparably low in activity, similar to the activity of homoinactive pairs. Consistently, the expression levels of active strands across active pair types were significantly higher than those of inactive strands (**Figure 3I**). The active strands of heteroactive pairs were expressed at significantly higher levels than their inactive strands, even though both strands originate and are processed from the same expressed pri-miRNA. This trend suggests degradation of the inactive strands, as might be expected given their ejection from the RISC.

It has been previously shown that guide-strand selection during Argonaute loading is influenced by both thermodynamic asymmetry and sequence composition. The predominantly selected loading strand exhibits a bias toward 5′-end uridine and a purine (R)-rich sequence, whereas the opposing passenger strand is enriched for 5′-end cytosine and pyrimidine (Y)-rich sequence^86–88^. To test whether our dataset can capture sequence features that contribute to the mutually exclusive process of loading strand selection in conjunction with miRNA activity, we defined a simplified sequence heuristic and estimated the likelihood of each strand sequence to be loading or passenger by its proximity to the sequence motifs 5′-UR_n_ and 5′-CY_n_, respectively (**Figures 3J, S5H,** and **S5I**). Using this sequence motif model, we independently assigned the loading or passenger sequence motif for each strand sequence of every miRNA pair. We also identified strands that showed ambiguous motif assignments (i.e., loading and passenger strand scores were equal, or both were above or below the motif assignment threshold) and excluded the pairs including strands with these ambiguous motif assignments from the subsequent paired analyses. This resulted in two homoactive pairs, 71 heteroactive pairs, and 433 homoinactive pairs with unambiguous motif assignments (**Figure 3K**). Among the heteroactive pairs, the active strands were enriched for the guide motif (66.20% and 33.80% for the loading and passenger assignments, respectively), whereas the inactive strands were skewed toward the passenger motif (43.66% and 56.34% for the loading and passenger assignments, respectively). We also found that 61.99% of the homoinactive pair strands in mESCs were assigned to the loading context. This could be because those miRNAs are loaded and active in other cell states across the mouse system.

We then focused on heteroactive miRNA pairs and their sequence motif patterns to test whether the paired sequence motifs of a pre-miRNA influence strand selection and cleavage activity. To do this, we assessed the sequence motif enrichment of one strand as a function of its activity status, given the assignment of the opposite strand’s sequence motifs with a different activity status (**Figure 3L**). The motif enrichment scores were normalized to the baseline biases of motif assignment among all heteroactive pairs to account for the observation that miRNA sequences were more frequently assigned as guide sequences. As supported by the model, we observed a global effect: assigning the loading motif to active strands led to enrichment of the passenger motif in inactive strands. Conversely, when inactive strands were assigned the passenger motif, the corresponding active strands showed enrichment for the loading motif. This supports the idea that the pre-miRNA sequence encodes information for guide/passenger selection. Consistent with this observation, the difference in expression level between the active and inactive strands was most prominent when they were assigned as loading and passenger, respectively (**Figure 3M**). Nevertheless, higher expression of active miRNA strands were also observed for the heteroactive pairs where both strands had loading motifs or passenger motifs.

Taken together, the cgRNA-based genome-wide screen provides a comprehensive approach to survey miRNA cleavage activity in a given cell state. In mESCs, we identified miRNAs with cleavage activity and found that miRNA loading into Argonaute is governed by paired-sequence features encoded on both strands of a single pri-miRNA.

### Recording genome-wide miRNA cleavage activities during SMC differentiation

The lentiviral introduction of the cgRNA library for capturing genome-wide mouse miRNA activities is a form of population-level molecular recording. miRNA activities are recorded with the restoration of the EGFP-coding sequence in the C→T base editing reporter and read out as the abundance of cgRNA miRT sequences in EGFP+ cells. We next tested whether this recording form could capture dynamic changes in miRNA activity in cells during state transitions, such as stem cell differentiation.

We chose a previously established smooth muscle cell (SMC) differentiation protocol that supplements 10 µM of all-trans retinoic acid (RA) to a simple serum-containing medium^89^. During this process, miR-295 expression is downregulated within 3 days, whereas miR-21 expression is maintained throughout the mature SMC state^90,91^. We first transduced a reporter toolkit mESC population harboring intronic Target-AIDmax in either the mESC state or after 6 days of SMC differentiation, with cgRNAs targeted by either miR-295 or miR-21 (**Figure 4A**). In the mESC state, cells transduced with the miR-295-targeted cgRNA had the highest EGFP+ rate compared to cells containing the miR-21-targeted cgRNA (57.31% and 26.59%, respectively). Cells transduced with a control cgRNA targeted by miR-122 showed minimal activation (1.67%). In contrast, cells transduced on day 6 of SMC differentiation with the miR-21-targeted cgRNA showed the highest EGFP+ rate (5.51%), while those with the miR-295-targeted cgRNA and the miR-122 control showed similar minimal editing levels (0.70% and 0.75%, respectively). Together, these results demonstrate that cgRNA has the capacity to record miRNA activity during the SMC differentiation process.

We then sought to measure genome-wide miRNA activity profiles during SMC differentiation. In this genome-wide screen, we chose to transduce Clone 7 of the reporter toolkit mESCs harboring intronic Target-AIDmax, since it showed the strongest miRT abundance profile correlations with the other two clones in the mESC analysis of the previous section (**Figures S5D** and **S5E**). We transduced the Clone 7 toolkit mESC line with the cgRNA library in either the mESC state or at different days of RA treatment during the SMC differentiation process (Day 3 and Day 6) (**Figure 4B**). Each cell population was transduced in triplicate at an infection rate of 0.1 or less, cultured for eight days, and sorted for EGFP+ cells. The miRT sequences were then PCR amplified from extracted genomic DNA of the EGFP+ and unsorted cell populations and quantified by deep sequencing as in the previous section. First, we confirmed that there was a significant correlation between the mean cgRNA enrichment profile of the Clone 7 toolkit mESCs in triplicate and the previous triple clone screen (**Figure S6A**). DESEQ2 analysis with an enrichment threshold of 2.0-fold and an FDR cutoff of 5% identified 70 miRNA hits in mESCs, 66 of which were captured in the triple clone screen (**Figure S6B**). Furthermore, consistent with the previous screen, a limited number of negative controls were identified as hits (**Figures S6C** and **S6D**), and gene set enrichment analysis showed enrichment for ESC-specific miRNAs (**Figure S6E**).

Applying the same enrichment parameters, we identified 100 and 36 active miRNAs from Day 3 and Day 6 of SMC differentiation, respectively (**Table S3**). Hierarchical clustering of cgRNA enrichment profiles for the union of the hits across the three differentiation stages revealed distinct patterns of stage-specific miRNA activity profiles (**Figure 4C**). Additionally, the trends observed in the independent cgRNA assays were recapitulated in the library-scale screens (**Figure 4D**). In the mESC state, both miR-295-3p and miR-21a-5p were significantly enriched (log_2_-fold change of 3.25 and 1.85, respectively), whereas miR-122-5p was not (–0.06). In Day 3 and Day 6, no significant enrichment was observed for miR-295-3p (0.29 and 0.32, respectively) or miR-122-5p (0.41 and 0.20, respectively), whereas miR-21a-5p maintained significant enrichment (2.13 and 1.91, respectively). Among the miRNAs specific to the library screen, members of the miR-290–295 cluster, a family of mESC-enriched miRNAs that includes miR-294-3p, miR-293-3p, miR-292a-3p, and miR-291a-3p, exhibited progressively reduced enrichment to varying extents from the mESC state to day 6 of differentiation. The enrichment of miR-292a-3p and miR-291a-3p was not detectable by Day 3 (0.84 and 0.93, respectively), whereas miR-294-3p and mmu-miR-293-3p showed some moderate level of sustained enrichment on Day 3 (1.01 and 1.02, respectively) before fully diminishing at Day 6 (0.19 and 0.20, respectively).

We next investigated miRNAs that showed increased activity during differentiation. Among 139 total miRNA hits across the three differentiation states, 110 were observed in Day 3 or Day 6 of SMC differentiation. 41 of these hits were shared with the mESC state, and 11 were unique to the differentiation states (**Figures 4E** and **4F**). Six out of the 11 SMC differentiation specific miRNA hits were associated with SMC-related processes, including a muscle-enriched microRNA (miR-206-5p), which is typically expressed in skeletal muscle^92^ but is also highly expressed in SMCs during lung development^93^; regulators of SMC quiescence and deproliferation^94–96^ (miR-200c-3p, miR-379-5p, miR-382-5p); a factor promoting proliferation while reducing migration (miR-582-5p) ^97^; and a regulator of striated muscle programming and angiogenesis (miR-411-5p)^98,99^. The remaining 5 have either been observed to be downregulated in other muscle disease phenotypes (miR-693-3p)^100^ or have not been characterized in SMCs (miR-28a-3p, miR-6419, miR-8106, and miR-878-5p). Our observation of their activities suggests their potential involvement in SMC differentiation. Additionally, gene set enrichment analysis of miRNAs enriched at Day 6 revealed terms such as muscle cells, myocytes, cardiac, and heart (**Figure 4G**). This likely reflects both the extensive annotation of miRNA expression in cardiac and skeletal muscle and the partial overlap of these miRNA programs with those in smooth muscle cells (SMCs). In contrast, enrichment analysis at Day 3 did not identify cell type-specific terms, likely due to the heterogeneous and incompletely specified cellular state, resulting in overlapping mESC, SMC, and other lineage signatures.

Accordingly, genome-wide cgRNA library transduction during SMC differentiation enabled temporal recording of miRNA activity across differentiation.

### Recording of coding gene expression with cgRNA editing

As demonstrated in the recording of miRNA activity profiles, cgRNA represents the first synthetic genetic element capable of scalable recording of endogenous molecular activities directly into DNA memory. Since miRNAs are deeply involved in gene regulatory networks, the recording of their activity profiles by a large set of corresponding cgRNAs integrated in a cell could be utilized to decode substantial information about cellular state and identity. However, a long-standing goal in the molecular recording field has been to directly record endogenous mRNA expression profiles as a practical, scalable vector of cell state information. At present, no such system has been established in mammalian cells. While ENGRAM^15^ enabled scalable recording of *cis*-regulatory element (CRE) reporter activities, it does not directly capture endogenous gene expression dynamics. The ENGRAM study also attempted to encode the Prime Editing guide RNA (pegRNA) directly within endogenous gene bodies but observed no genome editing in response to mRNA expression, presumably because the resulting pegRNA was insufficiently expressed. Here, we instead propose encoding synthetic miRNAs within an intron of an expressed mRNA and using the resulting miRNA as a recyclable intermediary that amplifies cgRNA activation.

The majority of miRNAs are encoded in the introns of genes^101,102^, and their pri-miRNAs are processed out by Drosha without interfering with splicing^102^. Additionally, encoding of a single pri-miRNA^103^ or a cluster of pri-miRNAs^104^ in a synthetic intron within the untranslated region (UTR) of synthetic mRNAs has been demonstrated for miRNA-based gene regulatory circuits^103,104^. Therefore, we hypothesized that a synthetic pri-miRNA encoded into an intronic region of an endogenous gene could activate a corresponding cgRNA and thereby enable recording of gene expression in a target DNA memory sequence (**Figure 5A**).

To test whether a synthetically encoded intronic pri-miRNA in a Pol2 transcript can trigger genome editing by its targeting cgRNA, we first transduced toolkit mESCs encoding the C→T base editing reporter and Target-AIDmax (**Figure 1C**) with a cgRNA targeted by a synthetic miRNA, miR-FF3, that has previously been demonstrated to be processed from pri-miR-FF3 encoded in a Pol2-transcribed intron^105^. We independently confirmed that cgRNA could be activated by pri-miR-FF3 expressed under the human U6 Pol3 promoter using the toolkit HeLa cells (**Figure 5B**). After establishing the “FF3 reporter” mESCs, we also established a clonal cell line (Clone 4). We then introduced the synthetic CAG Pol2 promoter (CMV enhancer/chicken β-actin promoter/rabbit β-globin intron)-derived mCherry gene with 3′ UTR encoding pri-miR-FF3 to the FF3 reporter mESCs or Clone 4 and investigated the EGFP+ rate among mCherry+ cells (**Figure 5C**). Because Pol3 promoters typically support rapid and high-level production of small RNA transcripts in comparison with Pol2 promoters^106^, we also tested a tandem repeat of three pri-miR-FF3 (pri-miR-FF3-boost) to enhance the sensitivity of the system. As expected, in both the FF3 reporter mESCs and Clone 4 cells, we observed that the CAG promoter-derived single pri-miR-FF3 and pri-miR-FF3-boost both conferred genome editing. The pri-miR-FF3-boost showed significantly higher reporter activity than the single pri-miR-FF3, whereas the filler sequence control showed negligible reporter activity. We also showed that in the heterogeneous FF3 reporter mESCs, the Pol2 promoter-derived pri-miR-FF3-boost showed activity comparable to that of a control hU6 Pol3 promoter-derived single pri-miR-FF3 (11.2% and 10.76% EGFP+ cells, respectively; **Figure 5D**).

Next, to investigate whether the synthetic pri-miRNA–cgRNA system could record endogenous mRNA expression, we employed CRISPR–Cas9-induced homology-directed repair to introduce a 2A–mCherry– intronic pri-miR-FF3-boost cassette into the 3′ end of the endogenous *Nanog* or *Acta2* coding sequence in wild-type mESCs (**Figure 5E**). Following successful knock-in, the engineered locus becomes a single Pol2 transcriptional unit, in which mCherry is co-translated with the target gene and subsequently separated by 2A peptide-mediated ribosomal skipping, while the pri-miR-FF3-boost cassette is spliced out via its synthetic intron and then processed to generate mature miR-FF3 from the same transcript. We chose *Nanog* and *Acta2* because *Nanog* exhibits mESC-specific expression and is absent in differentiated cells^107^, whereas the smooth muscle actin gene *Acta2* shows early and sustained expression in smooth muscle differentiation and mature smooth muscle cells^89^, while being absent in the mESC state. Successful *Nanog*–mCherry–FF3-boost knock-in cells were readily isolated by flow cytometric sorting of mCherry-positive cells. In contrast, because *Acta2* is not expressed in mESCs, a successful *Acta2*–mCherry–FF3 knock-in cell clone was established through clonal colony isolation followed by genotyping. We confirmed that *Nanog*–mCherry–FF3-boost cells lost mCherry expression by Day 3 of SMC differentiation, whereas *Acta2*–mCherry–FF3-boost cells exhibited no detectable mCherry expression in the mESC state but robustly activated mCherry expression by Day 3 of SMC differentiation (**Figures 5F** and **5G**).

We next stably introduced the Target-AIDmax construct with three introns (**Figure S5A**) together with the C→T base-editing reporter into the *Nanog*–mCherry–FF3-boost and *Acta2*–mCherry–FF3-boost cells via piggyBac transposon-mediated integration. Finally, we lentivirally delivered a miR-FF3-targeted cgRNA to cells either in the mESC state or at Day 6 of SMC differentiation and quantified base editing by %EGFP+ cells using flow cytometry (**Figure 5H**). Consistent with the mCherry expression pattern, *Nanog*–mCherry–FF3-boost cells showed a higher fraction of EGFP+ cells (6.31%) than *Acta2*–mCherry– FF3-boost cells (1.64%). In contrast, at Day 6 of SMC differentiation, *Acta2*–mCherry–FF3-boost cells showed a higher fraction of EGFP+ cells (6.50%) than *Nanog*–mCherry–FF3-boost cells (0.67%). Taken together, we concluded that the synthetic pri-miRNA–cgRNA system enables the recording of a target Pol2-derived gene expression signal into a target DNA sequence by genome editing.

Since each cgRNA couples a miRNA-responsive target sequence to a programmable DNA-targeting protospacer, distinct miRNA activities can be encoded into unique DNA memory addresses. In principle, multiplexed cgRNA architectures could therefore enable large-scale recording of miRNA activity states and endogenous gene expression profiles through corresponding pri-miRNA encoders (**Figure 5I**). While recording miRNA activity profiles represents a relatively direct implementation of the cgRNA framework, extending this concept toward genome-scale recording of endogenous gene expression states presents a substantially more ambitious challenge. Although comprehensive transcriptional recording was beyond the scope of this initial study, we sought to explore a conceptual route toward this goal by adapting principles from a previous gene-trapping strategy^108^.

We constructed a piggyBac plasmid vector encoding a splice acceptor sequence followed by mCherry and a synthetic intron encoding the pri-miR-FF3-boost cassette (**Figure 5J**). Upon random integration into a genomic 5′-TTAA-3′ site, observable expression of the piggyBac payload is activated when the insertion occurs in the correct orientation within an actively transcribed gene body. In this configuration, mCherry is expressed through splicing-mediated fusion to the upstream endogenous exon, while the embedded pri-miR-FF3-boost cassette is simultaneously processed to generate mature miR-FF3. We introduced this vector to the Clone 4 mESC line harboring the miR-FF3-targeted cgRNA, the C→T base editing reporter, and Target-AIDmax. The resulting cell population exhibited a broad distribution of mCherry expression, with 8.20% of cells classified as mCherry+ using an arbitrary threshold, whereas only 0.32% mCherry+ cells were observed for the no piggyBac control (**Figure 5K**). This indicated successful “transcriptional hitchhiking” of the payload into endogenously expressed genes. Furthermore, the fraction of EGFP+ cells from the C→T base editing reporter increased in parallel with increasing mCherry expression thresholds, reaching up to 4.91%, whereas the no piggyBac control exhibited only a background activation rate of 0.63% EGFP+ cells (**Figure 5H**). These results show that a pri-miRNA-encoding sequence randomly inserted in the genome can hitchhike the expression of an endogenous gene and convert the gene expression signal into genome editing.

Recent large-scale genome engineering studies have demonstrated antibiotic selection gene-dosing techniques that can mediate the random integration of approximately one hundred transgenes per cell^109,110^, with a substantial fraction of integrations occurring within intronic regions^110^. Coupling a library of synthetic pri-miRNAs integrated at diverse genomic loci with corresponding cgRNAs harboring distinct DNA memory addresses may therefore provide a framework for recording endogenous gene expression profiles at scale.

## DISCUSSION

Through systematic optimization of cgRNA architecture using synthetic miRNA mimics and endogenous miRNA activity, we demonstrated that cgRNA-mediated genome editing is conferred in a miRNA-dose-dependent manner. While AGO2 is broadly expressed across tissues and cell states^111^, miRNA-dependent genome editing could be further enhanced by AGO2 overexpression, thereby suggesting that RISC loading is a limiting step in cgRNA activation. In parallel, biochemical characterization and cryo-EM structural analyses revealed that the warped conformation of cgRNA inhibits activity through three complementary mechanisms: reduced Cas9 binding, impaired target DNA recognition, and diminished target DNA cleavage. Together, these findings provide a mechanistic framework explaining how endogenous miRNA activity can be translated into programmable genome editing.

Considering its activation dynamics with AGO2, cgRNA activity reflects a composite of mature miRNA abundance, Argonaute loading, target recognition, and miRNA-directed cleavage. A variety of miRNA-responsive ON and OFF reporters based on fluorescent and luciferase protein-coding mRNA have been previously established, where reporter expression is either repressed by miRNA^112,113^ or activated through miRNA cleavage^114,115^. These systems have been successfully applied at both small^112–115^ and large scales^116–118^ to profile miRNA function. However, their readouts are transient expressions and do not readily enable multiplexed recording of numerous miRNA activities into distinct genomic memory addresses within the same cell.

Previously, MICR represented the only approach to directly couple miRNA activity to sgRNA activation through miRNA-mediated cleavage^53^. However, MICR relies on a substantially more complex architecture in which a Pol 2-transcribed sgRNA must undergo two independent miRNA cleavage events to remove both the 5′ cap and 3′ poly(A) tail prior to activation. In contrast, cgRNA is a simple permutation of sgRNA sequence domains with a single miRNA target and retains a compact architecture comparable in size to the original sgRNA itself. This simplicity facilitates scalable library construction and pooled screening, which were not established in the MICR study, enabling genome-wide profiling of miRNA activity states.

As such, we demonstrated robust activation of cgRNA by endogenous miRNAs in both human and mouse cells and throughout stem cell differentiation. Importantly, pooled genome-wide screening revealed that cgRNA can be used to profile miRNA activity at scale, uncovering dynamic changes in functional miRNA programs during differentiation. Beyond miRNA sensing, we also established a proof-of-concept framework to convert endogenous gene expression into genome-editing signals via engineered pri-miRNA intermediates. This result extends the scope of cgRNA from a miRNA-responsive genome editing system to a more general molecular recording framework, in which transcriptional activity can be transformed into durable DNA memory. Notably, recent advances in large-scale piggyBac-mediated genome engineering^109,110^ suggest a potential path toward transcriptome-scale deployment, where diverse endogenous transcriptional programs could be coupled to distinct genomic memory addresses through synthetic pri-miRNA–cgRNA pairs.

To our knowledge, cgRNA represents the first scalable genome-editing platform capable of directly coupling endogenous miRNA activity and endogenous gene expression to programmable DNA recording in mammalian cells.

The high sensitivity, specificity, and tractability of cgRNA and the pervasive role of miRNA-mediated regulation across development, physiology, and disease suggest several immediate avenues for biological and biomedical applications. We envision three directions in which cgRNA may have a substantial impact: (1) miRNA-responsive genetic circuits for therapeutics, (2) spatial and single-cell biology, and (3) molecular event recording of cellular state transitions and differentiation trajectories.

### Application of cgRNA in miRNA-responsive genome editing and cell state engineering

First, since disease-associated and cell state-specific miRNA programs have been extensively characterized across diverse physiological and pathological contexts^42–50^, cgRNA may provide a powerful new framework for therapeutic genome editing and cell engineering. Fluorescent and mRNA regulatory protein-based miRNA-responsive ON and OFF reporter systems have previously been proposed for the construction of synthetic genetic circuits^112–115,119^. However, such systems remain constrained by signal leakage, limited dynamic range, and the transient nature of transcriptional and translational outputs. For example, OFF-switch architectures frequently exhibit residual basal activity, whereas ON-switch architectures often generate insufficient signal amplitudes to robustly propagate information through downstream circuit layers.

By contrast, conversion of molecular signals into DNA-state transitions provides a fundamentally different mode of biological computation. As demonstrated repeatedly across synthetic biology^13,14,16^, DNA modifications can serve as stable, digital, and heritable representations of transient molecular events that enable the construction of complex genetic circuits with improved robustness and scalability. Because cgRNAs can be readily programmed against diverse target miRNAs and coupled to distinct genomic memory addresses, they offer a scalable framework for translating endogenous miRNA activities into programmable DNA outputs. Importantly, cgRNA-mediated genome editing is not limited to memory recording. The resulting DNA modifications could be used to activate, deactivate, or reconfigure a wide range of genetic elements from protein-coding reading frames, site-specific recombination substrates, sgRNAs, and even other cgRNAs. Therefore, cgRNA introduces a novel and versatile platform for molecular programmability in which endogenous miRNA activities serve as inputs to precision genome engineering and synthetic cellular decision-making.

### Utilizing cgRNA to probe spatial and single-cell biology

Second, we propose that cgRNA may substantially expand the study of miRNA biology in single-cell contexts and scale up spatial genomics broadly. Although changes in miRNA expression have been extensively associated with developmental processes and disease progression in mice and humans^42–50^, methods for profiling miRNA activity remain substantially less developed than those for transcriptome analysis. No current technology can comprehensively measure miRNA activity profiles at a scale comparable to scRNA-seq approaches, and integration of miRNA activity with other single-cell genomic modalities remains challenging. Moreover, miRNA expression does not necessarily equate to miRNA function, as mature miRNAs can differ substantially in their regulatory pathways^66,86–88,120–123^.

By coupling miRNA activity to genome editing, cgRNA provides a direct route for converting functional miRNA states into genetically encoded information. As demonstrated in this study, individual miRNA activities can be linked to fluorescent reporter activation to isolate cells. Linkage of specific miRNA activities to downstream scRNA-seq or single-cell multiomics analysis can then be used to connect functional miRNA activity to cellular state (**Figure 6A top-left**). More broadly, sequencing of single cgRNA-associated molecular records alongside transcriptomic profiles, conceptually analogous to the mapping of information in Perturb-seq, could enable scalable interrogation of cell-state-specific miRNA activities across heterogeneous populations. While this described method only observes miRNA activity at a cell-type population level, further expanding the scope of such a method by multiplexing multiple diverse miRNA-targeted cgRNAs that correspond to distinct Pol 2 transcribed genomic memory addresses could potentially enable the reconstruction of high-dimensional miRNA activity profiles at a single-cell resolution (**Figure 6A top-middle**).

cgRNA may also provide a path toward scaling spatial genomics from tissue sections to whole organisms. Although spatial transcriptomic technologies have transformed our ability to study tissue organization, most current approaches remain restricted to two-dimensional sections or relatively small tissue volumes. As a result, comprehensive three-dimensional mapping of molecular activities across entire animal bodies remains an outstanding challenge.

One promising strategy is gene expression cartography, in which spatial information is reconstructed computationally by leveraging molecular landmarks whose spatial distributions are known. Conceptually, this approach resembles geographic cartography: just as the positions of cities can be inferred from recognizable landmarks, the spatial locations of cells can be inferred from characteristic patterns of gene expression. Existing implementations typically rely on reference maps generated using fluorescence *in situ* hybridization (FISH) or related technologies^124^. However, extending such approaches to high-resolution, whole-organism three-dimensional mapping remains technically challenging because volumetric FISH imaging is still limited in scale and throughput.

In addition to the aforementioned results, we preliminarily demonstrated that cgRNA can also mediate endogenous miRNA activity-dependent genome editing in mice. Pronuclear injection of a cgRNA targeting the liver-specific miRNA miR-122 together with the EGFP base-editing reporter and intronic Target-AIDmax into mouse zygotes, followed by embryo transfer, generated postnatal animals exhibiting liver-specific EGFP activation (**Figure 6A bottom-left** and **6B**). These results suggest that cgRNA-based recording may enable organism-scale mapping of endogenous miRNA activities. Looking forward, we envision combining cgRNA recording with tissue clearing, whole-organ imaging, and high-content microscopy to generate three-dimensional maps of multiple miRNA activities throughout the body. If single-cell states can be indexed by their miRNA activity profiles and these profiles can be spatially registered within a common three-dimensional coordinate system, large-scale single-cell genomic datasets may ultimately be projected back onto intact animal bodies (**Figure 6A bottom-middle**). Alternatively, if cgRNA-based recording can be extended to landmark mRNAs that define spatial coordinates, existing gene expression cartography frameworks^124^ could be directly leveraged to reconstruct cellular positions throughout intact organisms. Together, these capabilities raise the possibility of whole-organism molecular cartography, in which cellular states, regulatory activities, and spatial organization are integrated into a unified three-dimensional representation of an animal.

### Molecular event recording of cellular state transitions and differentiation trajectories with cgRNA

Third, cgRNA may substantially expand the scope of developmental recording and lineage tracing. There have been many advances in CRISPR-based lineage recording technologies in recent years. These technologies continue to show rapid progress towards the reconstruction of cellular ancestry across the entire development of the body of a model organism, such as mice. In parallel, single-cell genomics has transformed our ability to characterize cellular states and infer developmental trajectories. However, existing approaches primarily capture lineage relationships alongside terminal molecular states, thereby providing limited access to the dynamic regulatory programs that cells undergo during development. As a result, the molecular histories underlying cell fate decisions remain largely inaccessible.

A unique feature of cgRNA is that it enables the recording of endogenous molecular activities into DNA memory. We demonstrated genome-wide recording of miRNA activity and established a proof-of-principle framework for converting endogenous gene expression into genome-editing signals through engineered pri-miRNA intermediates. These capabilities suggest a path toward molecular recording systems that capture both cellular ancestry and their regulatory states traversed during development. Because miRNA activities and gene expression programs are intimately linked to cell identity and differentiation, their cumulative recording may provide a historical record of developmental progression that cannot be reconstructed from endpoint measurements alone.

Importantly, cgRNA-based recording can be integrated with existing CRISPR lineage recorders without fundamentally altering lineage recording architectures (**Figure 6 top-right**). In such a framework, lineage barcodes would capture patterns of cell division and ancestry, whereas cgRNA-mediated recording would capture dynamic regulatory activities experienced by each lineage. The resulting records would therefore contain complementary information describing where cells originated, how they diversified, and which molecular states they traversed throughout development.

The convergence of lineage recording, miRNA activity recording, gene expression recording, spatial registration, and single-cell genomics raises the possibility of reconstructing developmental processes at unprecedented scale and resolution (**Figure 6C bottom-right**). Under these contexts, future recording systems could potentially retrospectively reconstruct a whole-animal developmental process as a unified spatiotemporal process by linking cellular ancestry, regulatory histories, molecular states, and spatial organization within a common framework.

Taken together, cgRNA establishes a simple and versatile framework for coupling endogenous molecular activities to programmable genome editing and DNA memory. Because cgRNA retains an architecture comparable in simplicity to a canonical sgRNA while interfacing with miRNAs, one of the most pervasive regulatory layers in biology, we anticipate that its utility will extend well beyond the applications explored in this study. As our understanding of miRNA biology and molecular recording continues to expand, cgRNA may provide a broadly applicable foundation for future technologies that connect cellular regulation, genome engineering, and biological memory.

## ACKNOWLEDGMENTS

We thank the current and past members of the Yachie lab at the University of British Columbia, the University of Osaka, and the University of Tokyo for their constructive discussion and feedback. We thank Nika Shakiba, Mikko Taipale, Ross Jones, Peter Zandstra, Fabio Caliendo, and Ron Weiss for their valuable inputs. We also thank Tara Stach, Yvonne Cheung, Stephen Yu, Kaori Shiina, Andy Johnson, and Justin Wong for their technical support. We also thank Martin Kirzywinski for editing figures of this manuscript. This study was funded by the Japan Science and Technology Agency (JST)’s PRESTO Single Cell Analysis Program and CREST Cell Control (yuCell) Program, the Takeda Science Foundation, the Canada Foundation for Innovation (CFI), the Canadian Institutes of Health Research (CIHR) Project Grants, the Allen Distinguished Investigator Award (all to N.Y.), and the Japan Agency for Medical Research and Development (AMED) (to N.Y. and O.N.). The authors were also supported by the Canada Research Chair program (N.Y.), the Ministry of Science and Education (MEXT) Research Scholarship (A.A.), and the Natural Sciences and Engineering Research Council of Canada (NSERC) CGS-M Fellowship (A.A.). This work was partially supported by World Premier International Research Center Initiative (WPI), MEXT, Japan. Part of the high-throughput sequencing data analysis was conducted using the SHIROKANE Supercomputer at the University of Tokyo Human Genome Center.

## AUTHOR CONTRIBUTIONS

A.A. and N.Y. conceived cgRNA and designed the study.

A.A. optimized the cgRNA system, performed the major mammalian cell culture experiments, and conducted computational data analyses.

Y.S., R.N., S.O., and O.N. performed the cryo-EM analysis.

S.K. performed the biochemical assays.

Y.L., N.B., M.H., Y.K., and T.O. performed the *in vivo* assays.

H.O. and H.S. performed the small RNA sequencing.

A.S., B.K., S.C., and T.M.U. supported the plasmid construction and mammalian cell culture assays.

H.M. prepared the QUEEN-compatible GenBank files for the plasmids used in this study.

M.R., T.S., R.F., and P.H. supported the human stem cell-based assays.

H.A. supported the high-throughput sequencing.

A.A. and N.Y. wrote the manuscript.

N.Y. supervised the project.

## AI DISCLOSURE STATEMENT

We disclose that manuscript grammatical editing and formatting were supported by AI-based tools.

## DATA AVAILABILITY

All the raw sequencing data have been deposited in the NCBI Gene Expression Omnibus (GEO) at GSE319043 and GSE310325 for library-scale pooled cgRNA experiments and AQ-seq small RNA sequencing, respectively. Cryo-EM structural data have been deposited in the RCSB Protein Data Bank (PDB) at PDB 24LS and PDB 24LT. Other data, processed data, and associated scripts are available at https://github.com/yachielab/cgRNA. The list of plasmids used in this study can be found in **Table S4**. The backbone plasmid for cloning cgRNAs and other new plasmids necessary to reproduce the work have been deposited at Addgene.

## METHODS

### 1. Molecular cloning and plasmid documentation

Plasmids were constructed by restriction enzyme digestion, PCR or fusion PCR, T4 DNA ligation, Golden Gate assembly, or Gibson assembly. All plasmids used or generated in this study are listed in **Table S4.** The construction history of each newly generated plasmid was documented using QUEEN^125^. QUEEN generates an annotated GenBank (.gbk) file encoding the complete plasmid map and the sequence of operations used for its construction. Each QUEEN-generated .gbk file contains a quine code that enables retrieval of the plasmid construction process and regeneration of the same plasmid map in GenBank format. These QUEEN-generated .gbk files therefore provide a reproducible record of the plasmid construction protocols. Natural-language descriptions of the plasmid construction protocols were also included in the QUEEN .gbk files. Users can retrieve these protocols by executing QUEEN -- protocol_description --input [gbk file] in a QUEEN-installed environment.

All plasmids generated in this study, together with their corresponding QUEEN .gbk files, will be made available through Addgene and the project repository. Detailed natural-language protocols for the construction of individual plasmids are also provided in the detailed molecular cloning methods section below (Section 15). The MICR backbone plasmid PB4 was generously provided by Yangming Wang (Peking University). Newly generated plasmids were verified by Sanger sequencing of the assembled regions or by whole-plasmid sequencing.

### 2. Biochemical and structural analyses

Nucleic acids used in *in vitro* biochemical assays and cryo-EM analysis are listed in **Table S5**.

#### 2.1. Protein and RNA preparation for /in vitro/ cleavage assays, electrophoretic mobility shift assays and structural analyses

The wild-type *Streptococcus pyogenes* Cas9 (SpCas9) coding sequence was synthesized by Eurofins Genomics and assembled into a modified pET vector encoding an N-terminal His_6_–SUMO tag (LifeSensors). Mutations were introduced by a PCR-based method and confirmed by DNA sequencing. SpCas9 was expressed and purified using previously reported protocols^126,127^. Briefly, SpCas9 was expressed in *Escherichia coli* (*E. coli*) Rosetta 2 (DE3) cells. The transformed *E. coli* cells were cultured at 37 °C until the optical density at 600 nm (OD_600 nm_) reached 0.8, and protein expression was induced at 20 °C for 18–20 h by the addition of 1 mM isopropyl-β-d-thiogalactopyranoside (Nacalai Tesque).

The *E. coli* cells were collected by centrifugation and lysed by sonication in buffer A containing 20 mM Tris-HCl, pH 8.0, 1 M NaCl and 20 mM imidazole. The clarified lysate was incubated with Ni-NTA Superflow resin (Qiagen) at 4 °C for 1 h and loaded into an Econo-Column (Bio-Rad). The resin was washed sequentially with buffer A and buffer B containing 20 mM Tris-HCl, pH 8.0, 300 mM NaCl and 20 mM imidazole, and SpCas9 was eluted with buffer C containing 20 mM Tris-HCl, pH 8.0, 300 mM NaCl and 300 mM imidazole. The eluted protein was incubated overnight with SUMO protease produced in-house at 4 °C and then loaded onto a HiTrap Heparin column (GE Healthcare) equilibrated with buffer D containing 20 mM Tris-HCl, pH 8.0, and 300 mM NaCl. SpCas9 was eluted with a linear gradient of 0.3– 2 M NaCl and further purified on a HiLoad 16/600 Superdex 200 pg column (GE Healthcare) equilibrated with buffer E containing 20 mM Tris-HCl, pH 8.0, 500 mM NaCl, 2 mM MgCl_2_ and 1 mM DTT. Peak fractions were collected and stored at –80 °C until use.

The gRNAs were transcribed *in vitro* using T7 RNA polymerase and purified by 10% denaturing polyacrylamide gel electrophoresis containing 7 M urea. Purified gRNAs were stored at –20 °C until use.

#### 2.2. *In vitro* DNA cleavage assays

Purified wild-type SpCas9 and gRNA were mixed at a molar ratio of 1:2 to form the SpCas9−gRNA complex. For the cleaved cgRNA, an equimolar mixture of the crRNA and tracrRNA fragments was used as the gRNA. The SpCas9−gRNA complex (2 µL, final concentration 100 nM) was mixed with a linearized pUC119 plasmid containing a 20-nt target sequence followed by a 5′-TGG-3′ PAM (8 µL, 100 ng) and incubated at 37 °C in 10 µL reaction buffer (20 mM HEPES-KOH pH 7.5, 100 mM NaCl, 2 mM MgCl_2_, 5% glycerol, and 1 mM DTT). The reactions were stopped by the addition of quench buffer, containing EDTA (20 mM final concentration) and proteinase K (42 ng). The reaction products were resolved, visualized, and quantified with a MultiNA microchip electrophoresis device (Shimadzu).

#### 2.3. EMSA for Cas9–guide RNA binding to its corresponding DNA target

For target DNA binding, catalytically dead SpCas9 (dSpCas9)–guide RNA complexes were prepared as described for the *in vitro* cleavage assays. Increasing concentrations of dSpCas9–guide RNA complex (2 µl; 0, 50, 100, 200, 400 and 800 nM) were mixed with an annealed Cy5-labeled 39-bp target DNA (8 µl; 50 nM) containing a 20-nt target sequence and a TGG PAM. The binding reactions were performed at room temperature for 10 min in reaction buffer containing 20 mM HEPES-KOH, pH 7.5, 100 mM NaCl, 5 mM MgCl_2_, 5% glycerol and 1 mM DTT, and were stopped by the addition of quench buffer. The reaction products were resolved on 8% non-denaturing Novex TBE gels (Invitrogen) and visualized using an Amersham Imager 600 (GE Healthcare).

#### 2.4. Competitive EMSA for Cas9–cgRNA RNP complex formation

##### 2.4.1. RNA preparation

Double-stranded DNA fragments were used as templates for *in vitro* transcription. The cgRNA template was purchased from Twist Bioscience and amplified using Platinum SuperFi II PCR Master Mix (Thermo Fisher Scientific, no. 12368010 or 12368050) with the SK001/SK002 primer pair. The templates for sgRNA and nonspecific RNA were generated by annealing complementary oligonucleotides SK003/SK004 and SK005/SK006, respectively, in annealing buffer containing 10 mM Tris-HCl (Nacalai Tesque, no. 35436-01), 1 mM EDTA (Nacalai Tesque, no. 14347-21) and 100 mM NaCl (Nacalai Tesque, no. 690014).

RNAs were transcribed *in vitro* from the respective DNA templates, and template DNA was subsequently removed by DNase treatment using the MEGAshortscript T7 Transcription Kit (Thermo Fisher Scientific, no. AM1354) according to the manufacturer’s protocol. The reaction mixtures were then purified using the Monarch Spin RNA Cleanup Kit with a 50-µg capacity (NEB, no. T2040L). The RNAs were further purified by 8% denaturing polyacrylamide gel electrophoresis containing 8 M urea. Bands corresponding to full-length RNA were excised and eluted overnight in gel-extraction buffer containing 400 mM NaCl (Nacalai Tesque), 25 mM Tris-HCl, pH 7.6 (Nacalai Tesque), and 0.1% SDS (Nippon Gene, no. 311-90271). Eluted RNAs were recovered by phenol:chloroform:isoamyl alcohol extraction (Nacalai Tesque, no. 26058-54), precipitated with ethanol and resuspended in nuclease-free water.

##### 2.4.2. Cy5-labeled RNA preparation

RNA 3′-end labeling was performed in reactions containing 50 pmol RNA, 500 pmol pCp-Cy5 (Jena Bioscience, no. NU-1706-CY5), 1× T4 RNA ligase buffer and 20 U T4 RNA ligase (Thermo Fisher Scientific, no. EL0021). Reactions were incubated at 16 °C for 48 h while protected from light. For sgRNA labeling, DMSO (Sigma-Aldrich, no. D8418-100ML) was added to a final concentration of 10% (v/v). Labeled RNAs were purified using the Monarch RNA Cleanup Kit with a 10-µg capacity (NEB, no. T2030S) and eluted in nuclease-free water.

##### 2.4.3. Competitive EMSA

Unlabeled competitor RNAs at 12.5, 50, 100, 200 or 400 nM were mixed with 10 nM 3′-Cy5-labeled cgRNA or sgRNA in EMSA buffer containing 20 mM Tris-HCl, pH 7.6 (Nacalai Tesque), 150 mM KCl (Sigma-Aldrich, no. 60142-100ML-F), 5 mM MgCl_2_ (Sigma-Aldrich, no. M1028-100ML), 1 mM DTT (Sigma-Aldrich, no. 43816-50ML), 5 mM EDTA (Nacalai Tesque, no. 14347-21), 5% glycerol (Nacalai Tesque, no. 17017-35), 0.01% Tween-20 (Sigma-Aldrich, no. P9416-50ML) and 50 µg ml^−1^ heparin (Nacalai Tesque, no. 17513-41). The RNA mixtures were heated at 95 °C for 5 min and cooled on ice for 10 min to allow refolding.

Recombinant dCas9-NLS (100 nM; Nippon Gene, no. 314-09171) and 2 U murine RNase inhibitor (NEB, no. M0314), prepared in Diluent B (NEB, no. B8002S), were then added to the RNA mixtures, which were incubated at room temperature for 30 min. After incubation, 6× native loading buffer (Orange G; Nippon Gene, no. 317-90251) was added to the dCas9-NLS–RNA mixtures, and the mixtures were resolved on precast 8% native TBE gels (Novex TBE, 15-well; Thermo Fisher Scientific, no. EC62155BOX) at 200 V for 40 min under non-denaturing conditions. The gels were briefly rinsed with nuclease-free water, imaged using the ChemiDoc Touch MP Imaging System (Bio-Rad), and fluorescence intensities corresponding to bound RNA species were quantified.

##### 2.4.4. Quantification of EMSA signal intensities

Fluorescent EMSA images were analyzed using Image Lab software (Bio-Rad). The shifted RNA–protein complex band in each lane was defined as the region of interest, and background-corrected integrated intensities (adjusted volume) were obtained using lane-based background subtraction. For each sample, shift intensities were normalized to those obtained under the condition with the lowest competitor RNA concentration within the same experiment.

#### 2.5. Cryo-EM sample preparation and data collection

The SpCas9–cgRNA–target DNA ternary complex was reconstituted by mixing purified wild-type SpCas9, the 164-nt cgRNA, the 39-nt target strand and the 39-nt non-target strand at a molar ratio of 1:1.1:1.2:1.2. The resulting complex was purified by size-exclusion chromatography on a Superdex 200 Increase 10/300 column (GE Healthcare) equilibrated with buffer F containing 20 mM Tris, 150 mM NaCl, 2 mM MgCl_2_ and 1 mM DTT. The purified complex solution (A_260_ = 4.7) was applied to Au 300-mesh R1.2/1.3 grids (Quantifoil), which were freshly glow-discharged with 3 µl amylamine. The grids were prepared using a Vitrobot Mark IV (FEI) at 4 °C and 100% humidity, with a waiting time of 10 s and a blotting time of 4 s, and were then plunge-frozen in liquid ethane cooled by liquid nitrogen.

Cryo-EM data were collected using a Titan Krios G4 microscope (Thermo Fisher Scientific) operated at 300 kV and equipped with a Gatan Quantum-LS Energy Filter (GIF) and a Gatan K3 Summit direct electron detector operated in electron-counting mode (The University of Tokyo, Japan). Data were recorded using EPU software (Thermo Fisher Scientific) in standard mode, with a total electron dose of approximately 50 e^−^ Å^−2^ fractionated over 48 frames. The dose-fractionated movies were subjected to beam-induced motion correction and dose weighting using Patch Motion Correction, and contrast transfer function (CTF) parameters were estimated using Patch-based CTF estimation in cryoSPARC v4.4.0^128,129^.

#### 2.6. Single-particle cryo-EM data processing

Data were processed using the cryoSPARC v4.4.0 software platform^128^. From 8,625 motion-corrected and dose-weighted micrographs, 7,877,257 particles were automatically picked using Template Picker, followed by several rounds of reference-free 2D classification to select promising particles. The particles were further curated by several rounds of Heterogeneous Refinement using maps derived from Ab-initio Reconstruction as templates, resulting in 1,299,107 particles for the SpCas9–cgRNA binary complex and 678,308 particles for the SpCas9–cgRNA–target DNA ternary complex. Each particle set was subjected to CTF Refinement, Reference-Based Motion Correction and 3D Classification without alignment. Non-Uniform (NU) Refinement of the selected particles yielded the SpCas9–cgRNA binary complex and the SpCas9–cgRNA–target DNA ternary complex with partial R-loop maps at overall resolutions of 3.26 Å and 3.08 Å, respectively^130–132^. All reported resolutions were based on the gold-standard Fourier shell correlation with a cutoff of 0.143, and local resolution was estimated with BlocRes in cryoSPARC. For data-processing details, see **Figure S3.**

#### 2.7. Structural model building, refinement and validation

The models were built using the cryo-EM structures of the SpCas9–sgRNA binary complex (PDB: 7S37) and the SpCas9–sgRNA–target DNA ternary complex (PDB: 4O08), respectively, as reference models. After manual model building with Coot^133,134^, the models were refined against unsharpened half maps using Servalcat^135^, with reference-structure restraints generated by ProSMART^136^. The models were validated using MolProbity^137^. The 3DFSC sphericity was calculated using the 3DFSC processing server (https://3dfsc.salk.edu/upload/info/). Statistics for the 3D reconstruction and model refinement are summarized in **Table S1**. Molecular graphics were prepared using UCSF ChimeraX v1.10.1^138^.

### 3. Animals

B6D2F1 mice, used as wild-type mice in this study, were obtained from Japan SLC or bred in-house by crossing C57BL/6 females with DBA/2 males. Mice were maintained under a 14-h light/10-h dark cycle with *ad libitum* access to food and water. All animal experiments were approved by the Institutional Animal Care and Use Committee of the University of Osaka Medical School (protocol no. 24-020-007) and were conducted in compliance with all applicable guidelines and regulations.

### 4. Mammalian cell culture

#### 4.1. Human cell culture

The HeLa cell line was obtained from the American Type Culture Collection, and the Lenti-X HEK293T cell line was purchased from Takara Bio. Both cell lines were maintained in Dulbecco’s Modified Eagle’s Medium (DMEM; Thermo Fisher Scientific, no. 11965092) supplemented with 10% fetal bovine serum (FBS; Thermo Fisher Scientific, no. A5256701) at 37 °C with 5% CO_2_. Cells were routinely checked for mycoplasma by PCR.

The iPS11-10 neonatal male human induced pluripotent stem cell (hiPSC) line was purchased from Cell Applications. hiPSCs were maintained in mTeSR1 feeder-free medium (STEMCELL Technologies, no. 85850) on tissue-culture surfaces coated with Geltrex (Thermo Fisher Scientific, no. A1413301) at 37 °C with 5% CO_2_. Cells were routinely checked for mycoplasma by PCR.

#### 4.2. Mouse embryonic stem cell (mESC) culture

The mESC line was derived from embryos obtained by crossing 129(+Ter)/SvJcl females with C57BL/6NJcl males, as previously described^139^. mESCs were maintained in low-glucose DMEM (Thermo Fisher Scientific, no. 11885084) supplemented with 15% FBS, 1% GlutaMAX (Thermo Fisher Scientific, no. 35050061), 1% non-essential amino acids (Thermo Fisher Scientific, no. 11140050), 1% sodium pyruvate (Thermo Fisher Scientific, no. 11360070), 0.2 mM β-mercaptoethanol (Sigma-Aldrich, no. M6250-100ML), 3 µM CHIR99021 (Sigma-Aldrich, no. SML1046-5MG), 1 µM PD0325901 (Sigma-Aldrich, no. PZ0162-5MG) and 10^3^ U LIF (R&D Systems, no. 8878-LF-500/CF). Prior to plating mESCs, tissue-culture surfaces were coated with 0.2% gelatin (Sigma-Aldrich, no. G9391) in 1× phosphate-buffered saline (PBS; Thermo Fisher Scientific, no. 10010023) for 30 min at 37 °C with 5% CO_2_. For maintenance, mESC medium was changed every 2 days. Cells were routinely checked for mycoplasma by PCR.

### 5. Differentiation of mESCs to smooth muscle cells (SMCs)

mESCs were differentiated using a previously described all-*trans*-retinoic acid (RA)-based differentiation protocol^89^. mESCs were plated at high density on tissue-culture surfaces at 5 × 10^4^ cells per well of a 6-well plate or 9 × 10^6^ cells per T75 flask. The tissue-culture surfaces were precoated with 0.2% gelatin in 1× PBS for 1 h at 37 °C with 5% CO_2_ prior to cell plating. Differentiation medium consisted of DMEM/F-12 (Thermo Fisher Scientific, no. 11320033) supplemented with 10% FBS, 1 mM GlutaMAX, 0.1 mM non-essential amino acids, 0.1 mM sodium pyruvate, 0.1 mM β-mercaptoethanol and 10 µM RA (STEMCELL Technologies, no. 100-1045). Cultures were maintained at 37 °C with 5% CO_2_. The differentiation medium was replaced daily through day 9 and every 2 days thereafter.

### 6. Lentivirus production and transduction

#### 6.1. Single sgRNA, crRNA:tracrRNA, cgRNA, MICR and mCherry–IRES–EGFP Cas9 cleavage-activity reporter

Lenti-X HEK293T cells were plated 1 day prior to transfection in 6-well tissue-culture plates at a seeding density of 8 × 10^5^ cells per well. The following day, each well was transfected with 1,230 ng of an sgRNA, crRNA:tracrRNA, cgRNA or MICR transfer vector (**Table S4**), or the mCherry–IRES–EGFP transfer vector (pAA054), together with 1,170 ng of psPAX2 vector and 350 ng of pMD2.G vector using 8 µl of 1 mg ml^−1^ PEI MAX (Polysciences, no. 24765). At 16 h after transfection, the culture medium was aspirated and replaced with fresh medium. Three days after medium replacement, viral supernatant was harvested.

Lentivirus encoding mCherry–IRES–EGFP was stored at –80 °C prior to downstream cell-line generation. sgRNA, crRNA:tracrRNA, cgRNA and MICR lentiviral particles were either left unconcentrated or concentrated 10× using Lenti-X Concentrator (Takara Bio, no. 631231) according to the manufacturer’s protocol, resuspended in PBS and stored at –80 °C prior to downstream experiments. Unless otherwise specified, unconcentrated virus was used for HeLa cell experiments, whereas 10×-concentrated virus was used for mESC and hiPSC experiments.

#### 6.2. Large-scale production of pooled mouse cgRNA-library lentivirus

Lenti-X HEK293T cells were seeded in five 10-cm culture dishes at a density of 6 × 10^6^ cells per dish. The following day, each dish was transfected with 7,600 ng of pooled cgRNA-library transfer plasmids, 8,600 ng of psPAX2, 2,590 ng of pMD2.G and 60 µl of 1 mg ml^−1^ PEI MAX per dish. At 16 h after transfection, the culture medium was aspirated and replaced with fresh medium. Lentiviral supernatants were collected daily, pooled and maintained at 4 °C for no longer than 3 days after medium replacement. The pooled lentivirus was concentrated 10× using Lenti-X Concentrator (Takara Bio, no. 631231), resuspended in PBS and stored at –80 °C until use.

#### 6.3. General transduction conditions

Unless otherwise stated, lentiviral transductions were performed in the corresponding culture medium supplemented with 10 µg ml^−1^ polybrene (Sigma-Aldrich, no. TR-1003). At 24 h after transduction, the polybrene-containing medium was replaced with the corresponding basal culture medium or antibiotic-selection medium.

### 7. Mammalian cell line generation

#### 7.1. Generation of toolkit HeLa cell lines

##### 7.1.1. HeLa cell line containing the C→T EGFP base-editing reporter and Target-AIDmax

Transient transfection of plasmid DNA into HeLa cells was performed using FuGENE HD transfection reagent (Promega, no. E2311). HeLa cells were plated 1 day prior to transfection in a single well of a 6-well tissue-culture plate at a density of 2 × 10^5^ cells. The same transfection protocol was used to establish all subsequent HeLa toolkit cell lines.

The following day, cells were transfected with 622.5 ng of PB-INS-CAG-US1-eGFP-EF1a-Neo (pAA026), 1,022.5 ng of PB-INS-CAG-TargetAIDmax-nlsTagBFP2 (pAA028) and 352 ng of pCMV-PBase using 6 µl of FuGENE HD. At 3 days after transfection, cells were resuspended in sorting buffer (PBS containing 10% FBS), and BFP+ cells were isolated using a CytoFLEX SRT cell sorter (Beckman Coulter). Sorted cells were selected and expanded in a 6-well tissue-culture plate in antibiotic-selection medium containing 300 µg ml^−1^ Geneticin (G418 Sulfate; Thermo Fisher Scientific, no. 10131035). This toolkit HeLa cell line and subsequent HeLa cell lines carrying Geneticin resistance were maintained in medium supplemented with 100 µg ml^−1^ Geneticin after expansion.

##### 7.1.2. HeLa cell line containing the mCherry–IRES–EGFP reporter and wild-type SpCas9 or SaCas9

Using the transfection protocol described in Section 7.1.1, SpCas9 and SaCas9 toolkit HeLa cell lines were generated by transient transfection with 1,645.5 ng of PB-INS-CAG-spCas9-nlsTagBFP2 (pAA031) or PB-INS-CAG-saCas9-nlsTagBFP2 (pAA032), respectively, together with 352 ng of pCMV-PBase. At 3 days after transfection, cells were resuspended in sorting buffer, and BFP+ cells were isolated using a CytoFLEX SRT cell sorter. Sorted cells were expanded for 8 days, after which 1 × 10^5^ cells were transduced in suspension in a single well of a 12-well tissue-culture plate with lentiviral particles encoding mCherry–IRES–EGFP (pAA054) at a transduction efficiency of <20%. After 7 days, transduced cells were resuspended in sorting buffer, and mCherry+EGFP+BFP+ cells were isolated using a CytoFLEX SRT cell sorter.

##### 7.1.3. HeLa toolkit cells expressing human codon-optimized AGO2, Luc2 or AGO2ΔPIWI

Using the transfection protocol described in Section 7.1.1, toolkit HeLa cells were subjected to a second round of transfection with 1,645 ng of PB-INS-CAG-hAgo2-EF1-mCherry (pAA035), PB-INS-CAG-Luc2-EF1-mCherry (pAA036) or PB-INS-CAG-hAgo2ΔPIWI-EF1-mCherry (pAA037), together with 352 ng of pCMV-PBase. At 9 days after transfection, cells were resuspended in sorting buffer, and mCherry+BFP+ cells were isolated using a CytoFLEX SRT cell sorter (Beckman Coulter). Sorted cells were expanded in a 6-well tissue-culture plate in antibiotic-selection medium containing 300 µg ml^−1^ Geneticin.

#### 7.2. Generation of toolkit stem cell lines

##### 7.2.1. mESCs carrying the C→T EGFP base-editing reporter and Target-AIDmax variants

mESCs were seeded at 2 × 10^5^ cells per well in 6-well culture plates 1 day before transfection. Cells were transfected with 622.5 ng of PB-INS-CAG-US1-eGFP-EF1a-Neo (pAA026), 1,022.5 ng of PB-INS-CAG-TargetAIDmax-nlsTagBFP2 (pAA028) or its intron-containing variants (pAA029 and pAA030), and 352 ng of pCMV-PBase using 6 µl of Lipofectamine 2000 (Thermo Fisher Scientific, no. 11668027). At 3 days after transfection, cells were resuspended in sorting buffer, and BFP+ cells were isolated using a CytoFLEX SRT cell sorter. Sorted cells were selected and expanded in 6-well tissue-culture plates in antibiotic-selection medium containing 300 µg ml^−1^ Geneticin. These toolkit mESC lines and subsequent toolkit mESC lines carrying Geneticin resistance were maintained in medium containing 100 µg ml^−1^ Geneticin.

The functionality of the intron-containing and intron-free Target-AIDmax variants was confirmed by transduction with a base-editing reporter-targeting sgRNA. Toolkit mESCs, toolkit-2-intron mESCs and toolkit-3-intron mESCs were transduced in suspension at 5 × 10^4^ cells per well in 24-well tissue-culture plates with sgRNA (pAA069) lentiviral particles at a transduction efficiency greater than 40%. At 24 h after transduction, the culture medium was replaced with antibiotic-selection medium containing 3 µg ml^−1^ puromycin (Thermo Fisher Scientific, no. A1113803). At 6 days after transduction, cells were resuspended in flow-cytometry buffer (PBS containing 2% FBS), and the percentage of EGFP+ cells in the BFP+ population was analyzed using a CytoFLEX flow cytometer (Beckman Coulter).

##### 7.2.2. Toolkit mESCs carrying miR-FF3-targeted cgRNA

cgFF3-toolkit mESCs were generated by transducing 3 × 10^5^ toolkit mESCs in suspension in a single well of a 6-well tissue-culture plate with miR-FF3-targeted cgRNA (pAA091) lentiviral particles at a transduction efficiency of less than 20%. At 24 h after transduction, the culture medium was replaced with medium containing 3 µg ml^−1^ puromycin, and the selected population was expanded.

To establish a clonal line, cgFF3-toolkit mESCs were seeded at 5,000 cells per well in 6-well culture plates and cultured for 6 days to allow the formation of spatially separated colonies. A single colony was isolated, dissociated in 50 µl of trypsin–EDTA containing 0.05% trypsin and 0.53 mM EDTA (Thermo Fisher Scientific, no. 25300054), and expanded in one well of a 24-well culture plate.

##### 7.2.3. hiPSCs carrying the C→T EGFP base-editing reporter and Target-AIDmax

Transient transfection of plasmid DNA into hiPSCs was performed using Lipofectamine Stem transfection reagent (Thermo Fisher Scientific, no. STEM00003) according to the manufacturer’s instructions. hiPSCs were seeded at 2 × 10^5^ cells per well in 6-well culture plates 1 day before transfection. The following day, cells were transfected with 622.5 ng of PB-INS-CAG-US1-eGFP-EF1a-Neo (pAA026), 1,022.5 ng of PB-INS-CAG-TargetAIDmax-nlsTagBFP2 (pAA028) and 352 ng of pCMV-PBase using 5 µl of Lipofectamine Stem. At 3 days after transfection, cells were selected in medium containing 300 µg ml^−1^ Geneticin for 2 weeks. BFP+ cells were then isolated using a BD FACSMelody cell sorter (BD Biosciences) and expanded in one well of a 6-well plate.

#### 7.3. Generation of pri-miRNA knock-in toolkit mESCs

##### 7.3.1. Design of homology-directed repair donors and genome-targeting constructs

HDR donors and sgRNAs were designed using a previously described double-cut HDR-donor strategy^140^. sgRNA target sequences were selected in proximity to the stop codon of each target gene. Donors contained 300-bp homology arms corresponding to the genomic regions immediately 5′ and 3′ of the sgRNA cleavage site. Each donor was flanked by two copies of the genomic sgRNA target site to enable Cas9-mediated donor linearization and genomic cleavage using the same sgRNA. Synonymous substitutions were introduced into the residual sgRNA target sequence retained within the HDR donor to prevent self-targeting. A cassette encoding a T2A ribosome-skipping sequence, mCherry, a synthetic intron containing three tandem copies of pri-miR-FF3, and a bovine growth hormone polyadenylation signal was placed between the homology arms.

##### 7.3.2. Generation of Nanog-FF3-toolkit-3-intron mESCs

mESCs were transfected in suspension at 5 × 10^4^ cells per well in 24-well culture plates with 350 ng of PB-INS-CAG-spCas9-nlsTagBFP2-Nanog-sgRNA (pAA040) and 150 ng of pNANOG-HDR-mCherry-3x-miR-FF3 (pAA103) using 1.5 µl of Lipofectamine 2000. The same transfection conditions were also used to generate *Acta2*-FF3 mESCs, as described in Section 7.3.3. At 4 days after transfection, cells were resuspended in sorting buffer, and mCherry+ cells were isolated using a CytoFLEX SRT cell sorter. Following sorting, cells were expanded for 4 days in one well of a 6-well tissue-culture plate.

A portion of the expanded *Nanog*-FF3 mESC population was then subjected to SMC differentiation. At 3 days after the start of differentiation, cells were imaged by fluorescence microscopy using a Keyence BZ-X800 high-resolution imaging system (Keyence) to confirm loss of mCherry expression.

After confirmation of differentiation-associated loss of mCherry expression, 2 × 10^5^ *Nanog*-FF3 mESCs were transfected in suspension in a single well of a 6-well tissue-culture plate with 622.5 ng of PB-INS-CAG-US1-eGFP-EF1a-Neo (pAA026), 1,022.5 ng of the three-intron PB-INS-CAG-TargetAIDmax-nlsTagBFP2 (pAA030) and 352 ng of pCMV-PBase using 6 µl of Lipofectamine 2000. At 3 days after transfection, cells were resuspended in sorting buffer, and mCherry+BFP+ cells were isolated using a CytoFLEX SRT cell sorter. Sorted cells were expanded in medium containing 300 µg ml^−1^ Geneticin. The resulting population was designated *Nanog*-FF3-toolkit-3-intron mESCs.

##### 7.3.3. Generation of Acta2-FF3-toolkit-3-intron mESCs

mESCs were transfected with 350 ng of PB-INS-CAG-spCas9-nlsTagBFP2-Acta2-sgRNA (pAA041) and 150 ng of pACTA2-HDR-mCherry-3x-miR-FF3 (pAA104) using 1.5 µl of Lipofectamine 2000. At 4 days after transfection, cells were resuspended in sorting buffer, and BFP+ cells were isolated using a CytoFLEX SRT cell sorter.

Following sorting, cells were seeded sparsely at 5,000 cells per well in 6-well tissue-culture plates to facilitate the formation of mESC colonies and cultured for 6 days. Individual colonies were then isolated and dissociated in 50 µl of trypsin. For each colony, 25 µl of the dissociated cell suspension was transferred to a well of a 96-well plate for cell-line expansion, and the remaining 25 µl was transferred to a separate 96-well plate for genotyping PCR. Cells allocated for genotyping were expanded for 3 days, and genomic DNA was extracted by NaOH lysis. Cells were lysed in 100 µl of 50 mM NaOH, heated to 95 °C for 15 min and then cooled to 4 °C in a thermocycler (Bio-Rad). The lysates were neutralized with 10 µl of 1 M Tris-HCl, pH 8.0 (Thermo Fisher Scientific, no. 15568025).

For genotyping PCR, 3 µl of extracted genomic DNA from each clone was amplified using primer pair AA120/AA121 and Phusion High-Fidelity DNA Polymerase (NEB, no. M0530L) according to the PCR conditions described in **Supplementary Note 1**, with an annealing temperature of 64 °C and an extension time of 30 s. Clones showing the expected knock-in bands were expanded from 96-well to 6-well tissue-culture plates.

Genotype-positive clones were then subjected to SMC differentiation. At 3 days after the start of differentiation, cells were imaged by fluorescence microscopy using a Keyence BZ-X800 high-resolution imaging system to confirm differentiation-induced mCherry expression. A single clone showing high mCherry expression after 3 days of differentiation was selected for downstream engineering and experiments.

Following validation, 2 × 10^5^ *Acta2*-FF3 mESCs were transfected in suspension in a single well of a 6-well tissue-culture plate with 622.5 ng of PB-INS-CAG-US1-eGFP-EF1a-Neo (pAA026), 1,022.5 ng of the three-intron PB-INS-CAG-TargetAIDmax-nlsTagBFP2 (pAA030) and 352 ng of pCMV-PBase using 6 µl of Lipofectamine 2000. BFP+ cells were isolated using a CytoFLEX SRT cell sorter and expanded in medium containing 300 µg ml^−1^ Geneticin. The resulting cell line was designated *Acta2*-FF3-toolkit-3-intron mESCs.

### 8. cgRNA singleton activity assays

#### 8.1. piggyBac transfection-based assessment of genome editing by single cgRNAs and sgRNAs in mESCs

mESCs were plated 1 day prior to transfection in 48-well tissue-culture plates at a density of 1 × 10^4^ cells per well. The following day, cells were transfected with 82 ng of an all-in-one reporter–Target-AID vector (pAA014 or pAL001–pAL016), 82 ng of a cgRNA (pAL035–pAL053) or sgRNA (pAL018–pAL034) vector, and 68.76 ng of pCMV-PBase using 0.75 µl of Lipofectamine 2000. At 3 days after transfection, 50% of the cells in each transfected well were passaged into mESC medium supplemented with 5 µg ml^−1^ blasticidin (Thermo Fisher Scientific, no. A1113903). At 3 days after passage and selection, cells were resuspended in flow-cytometry buffer, and the percentage of EGFP+ cells in the BFP+ population was analyzed using a CytoFLEX flow cytometer.

#### 8.2. Lentivirus-based miRNA-activity-dependent genome editing by single cgRNAs in mammalian cell culture

##### 8.2.1. Comparison of cgRNA with sgRNA, crRNA:tracrRNA and MICR in HeLa toolkit cells

Toolkit HeLa cells were transduced in suspension at 5 × 10^4^ cells per well in 24-well plates with lentiviral particles encoding sgRNA (pAA065 and pAA066), crRNA:tracrRNA (pAA063 and pAA064), MICR (pAA055 and pAA059) or cgRNA (pAA056–pAA058 or pAA060–pAA062) at a transduction efficiency of 10% or less. At 24 h after transduction, the culture medium was replaced, and transduced cells were passaged and expanded in 6-well tissue-culture plates for 7 days. After expansion, cells were resuspended in sorting buffer, and mCherry+ cells were isolated using a CytoFLEX SRT cell sorter and expanded in 6-well tissue-culture plates.

After expansion, sgRNA-, crRNA:tracrRNA-, MICR- and cgRNA-transduced cell lines were plated at 4 × 10^4^ cells per well in 24-well tissue-culture plates. The following day, cells were transfected with 0.156 nM hsa-miR-122-5p mimic (Dharmacon, 5′-UGGAGUGUGACAAUGGUGUUUG-3′) or non-targeting control mimic (Dharmacon, 5′-UCACAACCUCCUAGAAAGAGUAGA-3′) using 0.5 µl of Lipofectamine RNAiMAX (Thermo Fisher Scientific, no. 13778030). At 6 days after mimic transfection, cells were either resuspended in flow-cytometry buffer, and the percentage of EGFP+ cells in the mCherry+BFP+ population was analyzed using a CytoFLEX flow cytometer, or imaged by fluorescence microscopy using a Keyence BZ-X800 high-resolution imaging system. Prior to imaging, cells were fixed with 4% paraformaldehyde (PFA; Sigma-Aldrich, no. P6148-500G) for 20 min. Following fixation, cells were washed twice with PBS to remove PFA and maintained in PBS for imaging.

MICR (pAA055)- and cgRNA (pAA056–pAA058)-transduced cell lines containing base-editing reporter-targeting spacer sequences were also tested across a range of miRNA mimic concentrations to generate a dose–response curve. These MICR- and cgRNA-transduced cell lines were plated at 4 × 10^4^ cells per well in 24-well tissue-culture plates. The following day, cells were transfected with 5, 2.5, 1.25, 0.625, 0.312, 0.156, 0.078, 0.039, 0.020, 0.009, 0.005 or 0 nM hsa-miR-122-5p mimic using 0.5 µl of Lipofectamine RNAiMAX. At 6 days after mimic transfection, cells were resuspended in flow-cytometry buffer and analyzed using the same parameters as in the single-dose experiment above with a CytoFLEX flow cytometer.

##### 8.2.2. Effect of AGO2 overexpression on cgRNA activity

AGO2, Luc2 or AGO2ΔPIWI toolkit HeLa cells were transduced in suspension at 1 × 10^5^ cells per well in 24-well tissue-culture plates with lentiviral particles encoding sgRNA (pAA083), miR-21-targeted cgRNA (pAA084) or miR-122-targeted cgRNA (pAA085) at a transduction efficiency of 40% or higher. At 24 h after transduction, the culture medium was replaced with antibiotic-selection medium containing 3 µg ml^−1^ puromycin. At 6 days after selection, cells were resuspended in flow-cytometry buffer, and the percentage of EGFP+ cells in the mCherry+BFP+ population was analyzed using a CytoFLEX flow cytometer.

A portion of the toolkit cell lines transduced with sgRNA or miR-122-targeted cgRNA was independently expanded and replated at 4 × 10^4^ cells per well in 24-well tissue-culture plates. The following day, cells were transfected with 5 nM hsa-miR-122-5p mimic or non-targeting control mimic using 0.5 µl of Lipofectamine RNAiMAX. At 6 days after mimic transfection, cells were resuspended in flow-cytometry buffer and analyzed using the same parameters as described above with a CytoFLEX flow cytometer.

##### 8.2.3. Effect of miRNA target truncation on cgRNA activity

Toolkit HeLa cells were transduced in suspension at 5 × 10^4^ cells per well in 24-well tissue-culture plates with cgRNA (pBK009 or pAA048–pAA050) lentiviral particles at a transduction efficiency of 20% or less. At 24 h after transduction, the culture medium was replaced, and transduced cells were passaged and expanded in 6-well tissue-culture plates for 9 days post-transduction.

After expansion, cgRNA-transduced cell lines were plated at 4 × 10^4^ cells per well in 24-well tissue-culture plates. The following day, cells were transfected with 5 nM hsa-miR-122-5p mimic or non-targeting control mimic using 0.5 µl of Lipofectamine RNAiMAX. At 6 days after mimic transfection, cells were resuspended in flow-cytometry buffer, and the percentage of EGFP+ cells in the mCherry+BFP+ population was analyzed using a CytoFLEX flow cytometer.

##### 8.2.4. Evaluation of cgRNA activity with wild-type SpCas9 and SaCas9

For miRNA mimic experiments, SpCas9 and SaCas9 toolkit HeLa cells were transduced in suspension at 3 × 10^5^ cells per well in 6-well tissue-culture plates with lentiviral particles encoding a miR-122-targeted SpCas9 cgRNA (pAA093) for the SpCas9 toolkit cell line or a miR-122-targeted SaCas9 cgRNA (pAA096) for the SaCas9 toolkit cell line, at transduction efficiencies of 40% or higher. At 24 h after transduction, the culture medium was replaced with antibiotic-selection medium containing 3 µg ml^−1^ puromycin, and cells were expanded to confluence. Transduced cells were then plated at 4 × 10^4^ cells per well in 24-well tissue-culture plates. The following day, cells were transfected with 5 nM hsa-miR-122-5p mimic or non-targeting control mimic using 0.5 µl of Lipofectamine RNAiMAX. At 12 days after mimic transfection, cells were resuspended in flow-cytometry buffer, and the percentage of EGFP+ cells in the mCherry+BFP+ population was analyzed using a CytoFLEX flow cytometer.

For endogenous miRNA experiments, SaCas9 toolkit HeLa cells were transduced in suspension at 3 × 10^5^ cells per well in 6-well tissue-culture plates with lentiviral particles encoding miR-21- or miR-122-targeted SaCas9 cgRNAs (pAA095 and pAA096, respectively) at a transduction efficiency of 40% or higher. At 24 h after transduction, the culture medium was replaced with puromycin-containing medium as described above, and cells were expanded to confluence. At 12 days after transduction, cells were resuspended in flow-cytometry buffer and analyzed using the same parameters as described above with a CytoFLEX flow cytometer.

##### 8.2.5. Effect of lentiviral transduction efficiency on cgRNA activity

Prior to transduction, 10×-concentrated cgRNA lentiviral particles (pBK003 and pBK009) were serially diluted twofold for a total of 11 dilution steps. Toolkit HeLa cells were transduced in suspension at 2 × 10^4^ cells per well in 48-well tissue-culture plates with each dilution of the cgRNA lentiviral particles. At 24 h after transduction, the culture medium was replaced. At 6 days after transduction, cells were resuspended in flow-cytometry buffer, and the percentage of EGFP+ cells in the mCherry+BFP+ population was analyzed using a CytoFLEX flow cytometer.

##### 8.2.6. Activity of endogenous miRNA-targeted cgRNAs in HeLa cells and mESCs

Toolkit HeLa cells and toolkit mESCs were transduced in suspension at 2 × 10^4^ cells per well in 48-well tissue-culture plates with lentiviral particles encoding either sgRNA (pAA042) or cgRNA (pBK001– pBK009) at a transduction efficiency of 20% or less. At 24 h after transduction, the culture medium was replaced. At 6 days after transduction, cells were resuspended in flow-cytometry buffer, and the percentage of EGFP+ cells in the mCherry+BFP+ population was analyzed using a CytoFLEX flow cytometer.

##### 8.2.7. Activity of endogenous miRNA-targeted cgRNAs in hiPSCs

Toolkit hiPSCs were plated 1 day prior to transduction at 3 × 10^5^ cells per well in 6-well tissue-culture plates and then transduced in their respective culture medium without polybrene supplementation with lentiviral particles encoding either sgRNA (pAA083) or cgRNA (pAA087–pAA090) at a transduction efficiency of 40% or less. At 48 h after transduction, the culture medium was replaced with antibiotic-selection medium containing 1 µg ml^−1^ puromycin. At 6 days after transduction, cells were resuspended in phenol red-free DMEM (Thermo Fisher Scientific, no. 31053028), and the percentage of EGFP+ cells in the BFP+ population was analyzed using a BD Fortessa I analyzer (BD Biosciences).

##### 8.2.8. Evaluation of cgRNA V2 and V3 circularization

Toolkit HeLa cells were transduced in suspension at 5 × 10^4^ cells per well in 24-well tissue-culture plates with lentiviral particles encoding either cgRNA V3 (pBK003 and pBK009) or cgRNA V4 (pAA106 and pAA107) at a transduction efficiency of 20% or less. At 24 h after transduction, the culture medium was replaced. At 6 days after transduction, cells were resuspended in flow-cytometry buffer, and the percentage of EGFP+ cells in the mCherry+BFP+ population was analyzed using a CytoFLEX flow cytometer.

Toolkit mESCs were transduced in suspension at 2 × 10^4^ cells per well in 48-well tissue-culture plates with lentiviral particles encoding either cgRNA V3 (pBK001 and pBK009) or cgRNA V5 (pAA045 and pAA046) at a transduction efficiency of 20% or less. At 24 h after transduction, the culture medium was replaced. At 6 days after transduction, cells were resuspended in flow-cytometry buffer and analyzed using the same parameters as described above with a CytoFLEX flow cytometer.

##### 8.2.9. Evaluation of cgRNA loop length on genome-editing activity

Toolkit HeLa cells were transduced in suspension at 2 × 10^4^ cells per well in 48-well tissue-culture plates with cgRNA lentiviral particles containing increasing insertion lengths between the 3′ end of the gRNA scaffold and the 5′ end of the miRNA target (pAA070, pAA071 or pAA072–pAA081) at a transduction efficiency of 40% or less. At 24 h after transduction, the culture medium was replaced with antibiotic-selection medium containing 3 µg ml^−1^ puromycin. At 6 days after transduction, cells were resuspended in flow-cytometry buffer, and the percentage of EGFP+ cells in the BFP+ population was analyzed using a CytoFLEX flow cytometer.

##### 8.2.10. Endogenous miRNA-dependent cgRNA activity in SMCs

Toolkit-2-intron mESCs were either subjected to SMC differentiation as described in Section 5 or maintained as mESCs in 6-well tissue-culture plates. On day 6 of differentiation, SMCs were transduced with cgRNA lentiviral particles (pAA084–pAA086) at a transduction efficiency of 20% or less. mESCs were plated 1 day prior to transduction at 3.5 × 10^4^ cells per well in 24-well tissue-culture plates and then transduced with cgRNA lentiviral particles (pAA084–pAA086) at a transduction efficiency of 20% or less. At 24 h after transduction, the culture medium for both SMCs and mESCs was replaced with antibiotic-selection medium containing 3 µg ml^−1^ puromycin. At 6 days after selection, cells were resuspended in flow-cytometry buffer, and the percentage of EGFP+ cells in the BFP+ population was analyzed using a CytoFLEX flow cytometer.

### 9. Engineered pri-miRNA and gene-expression recording with cgRNA

#### 9.1. Expression of cgRNA by hU6-driven pri-miR-FF3 in HeLa toolkit cells

Toolkit HeLa cells were co-transduced in suspension at 1 × 10^5^ cells per well in 12-well tissue-culture plates with lentiviral particles encoding miR-FF3- or miR-MT1-targeted cgRNA (pAA091 or pAA092, respectively) and pri-miR-FF3 (pAA052) at a transduction efficiency of 40% or higher. At 24 h after transduction, the culture medium was replaced with antibiotic-selection medium containing 3 µg ml^−1^ puromycin. At 6 days after transduction, cells were resuspended in flow-cytometry buffer, and the percentage of EGFP+ cells in the mCherry+BFP+ population was analyzed using a CytoFLEX flow cytometer.

#### 9.2. Expression of cgRNA by U6- and CAG-driven pri-miR-FF3 in mESCs

cgFF3-toolkit mESCs and cgFF3 Clone 4 toolkit mESCs were plated 1 day prior to transfection in 24-well tissue-culture plates at a density of 5 × 10^4^ cells per well. The following day, cells were transfected with 400 ng of CAG promoter-driven pri-miR-FF3 (pAA098 and pAA099) or an empty-intron control (pAA097), together with 70 ng of pCMV-PBase, using 1 µl of Lipofectamine 2000. At 6 days after transfection, cells were resuspended in flow-cytometry buffer, and the percentage of EGFP+ cells in the mCherry+BFP+ population was analyzed using a CytoFLEX flow cytometer.

Additionally, cgFF3-toolkit mESCs were transduced in suspension at 5 × 10^4^ cells per well in 24-well tissue-culture plates with pri-miR-FF3 (pAA052) lentiviral particles at a transduction efficiency of 40% or less. At 6 days after transduction, cells were resuspended in flow-cytometry buffer and analyzed using the same parameters as described above with a CytoFLEX flow cytometer.

#### 9.3. Recording endogenous *Nanog* and *Acta2* expression during SMC differentiation

*Nanog*-FF3 mESCs and *Acta2*-FF3 mESCs were either subjected to SMC differentiation as described in Section 5 or maintained as mESCs in 6-well tissue-culture plates. On day 6 of differentiation, SMCs were transduced with miR-FF3-targeted cgRNA (pAA091) lentiviral particles at a transduction efficiency of 40% or less. mESCs were plated 1 day prior to transduction at 3.5 × 10^4^ cells per well in 24-well tissue-culture plates and then transduced with miR-FF3-targeted cgRNA (pAA091) lentiviral particles at a transduction efficiency of 40% or less. At 24 h after transduction, the culture medium for both SMCs and mESCs was replaced with antibiotic-selection medium containing 3 µg ml^−1^ puromycin. At 6 days after transduction, cells were resuspended in flow-cytometry buffer, and mCherry expression and the percentage of EGFP+ cells in the BFP+ population were analyzed using a CytoFLEX flow cytometer.

#### 9.4. Random integration of pri-miR-FF3 gene-trap cassettes

cgFF3 Clone 4 toolkit mESCs were plated 1 day prior to transfection in 12-well tissue-culture plates at a density of 1 × 10^5^ cells per well. The following day, cells were transfected with 800 ng of PB-geneTrap-3x-miR-FF3 (pAA100), either together with 140 ng of pCMV-PBase or without pCMV-PBase, using 2 µl of Lipofectamine 2000. At 7 days after transfection, cells were resuspended in flow-cytometry buffer, and the percentage of EGFP+ cells in the mCherry+BFP+ population was analyzed across different thresholds of mCherry intensity using a CytoFLEX flow cytometer.

### 10. Flow-cytometry data analysis

All collected flow-cytometry data were analyzed in Python using the FlowCytometryTools package (https://eyurtsev.github.io/FlowCytometryTools/). Results of these analyses were visualized and plotted in R. All flow-cytometry analysis and plotting scripts are available at https://github.com/yachielab/cgRNA.

### 11. Genome-wide cgRNA screens

#### 11.1. Genome-wide recording of miRNA activity in mESCs

Three independently derived toolkit-2-intron mESC clones (Clone 3, Clone 7 and Clone 10) were screened separately. For each clonal population, 1 × 10^7^ mESCs were transduced in suspension with lentiviral particles generated from a library of cgRNA sensors containing targets for mouse miRNAs genome-wide (cgRNA library) at a transduction efficiency of 10% (approximately 500-fold coverage). Each transduced clonal population was plated in a T75 tissue-culture flask. At 24 h after transduction, the culture medium was replaced with antibiotic-selection medium containing 3 µg ml^−1^ puromycin. Puromycin selection was maintained during all subsequent medium changes and passages until 1 day prior to cell sorting; selection was removed on day 5 after transduction. On day 3 after selection, 6 × 10^6^ cells from each population were passaged and distributed across two T75 tissue-culture flasks.

On day 6 after transduction, each clonal population was resuspended in sorting buffer, and EGFP+ cells were isolated using a MoFlo Astrios cell sorter (Beckman Coulter), CytoFLEX SRT cell sorter or Cytek Aurora Cell Sorter (Cytek Biosciences). Prior to sorting, 2 × 10^6^ cells from each clonal population were set aside as an unsorted control. After sorting, genomic DNA was extracted from the sorted EGFP+ cells and unsorted control cells using the Monarch Genomic DNA Purification Kit (NEB, no. T3010S) according to the manufacturer’s protocol. genomic DNA from EGFP+ and unsorted cells was eluted in 50 µl and 100 µl of UltraPure Distilled Water (Thermo Fisher Scientific, no. 10977015), respectively.

#### 11.2. Genome-wide recording of miRNA activity during SMC differentiation

Toolkit-2-intron Clone 7 mESCs were subjected to mESC-to-SMC differentiation as described in Section 5. Independent differentiation cultures were established in triplicate for cgRNA recording initiated on day 3 or day 6 of differentiation. For each replicate, 9 × 10^6^ mESCs were distributed across three T75 tissue-culture flasks at 3 × 10^6^ cells per flask. On day 3 or day 6 of differentiation, adherent differentiating SMCs were transduced with lentiviral particles generated from the cgRNA library at a transduction efficiency of 10%. Given the starting population of 9 × 10^6^ cells and the expansive growth phase through day 6, the total library coverage was expected to remain approximately 500-fold. At 24 h after transduction, the culture medium was replaced with antibiotic-selection medium containing 3 µg ml^−1^ puromycin. Puromycin selection was maintained during all subsequent medium changes and passages until 1 day prior to cell sorting; selection was removed on day 7 after transduction.

On day 8 after transduction, each replicate population was resuspended in sorting buffer, and EGFP+ cells were isolated using a MoFlo Astrios cell sorter (Beckman Coulter), CytoFLEX SRT cell sorter or Cytek Aurora Cell Sorter. Prior to sorting, 2 × 10^6^ cells from each replicate were set aside as an unsorted control. After sorting, genomic DNA was extracted from the sorted EGFP+ cells and unsorted control cells using the Monarch Genomic DNA Purification Kit (NEB, no. T3010S) according to the manufacturer’s protocol. genomic DNA from EGFP+ and unsorted cells was eluted in 50 µl and 100 µl of UltraPure Distilled Water (Thermo Fisher Scientific, no. 10977015), respectively.

For recording in the mESC state, toolkit-2-intron Clone 7 mESCs were transduced in triplicate using the same protocol as described in Section 11.1. However, cell sorting and genomic DNA extraction were performed on day 8 after transduction rather than on day 6.

### 12. Sequencing library preparation

#### 12.1. Preparation of small RNA sequencing libraries for toolkit HeLa cells and toolkit mESCs

Total RNA was isolated from toolkit HeLa cells and toolkit mESCs using the mirVana miRNA Isolation Kit without phenol (Thermo Fisher Scientific, no. AM1561) according to the manufacturer’s protocol and eluted in nuclease-free water. RNA concentration was determined using a NanoDrop 2000 spectrophotometer (Thermo Fisher Scientific), and RNA integrity was confirmed by an RNA Quality Number (RQN) greater than 9 using the R1 Cartridge (BiOptic, no. C105210) on a Qsep100 DNA Fragment Analyzer (BiOptic). Total RNA samples were concentrated using an FD-1000 freeze-dryer (EYELA) to reduce the sample volume.

Small RNA sequencing libraries were prepared according to the AQ-seq method^65^, with modifications. Five micrograms of total RNA was resolved on a 15% denaturing polyacrylamide gel, and the 17–29-nt small RNA fraction was excised. The gel slices were fragmented by centrifugation through 0.5-ml tubes with holes in the bottom at 20,400 × g for 10 min into 2-ml collection tubes. The size-fractionated RNA was eluted from the gel fragments in 0.3 M NaCl at 25 °C with shaking at 1,500 rpm overnight. Gel debris was removed by centrifugation using Costar Spin-X centrifuge tubes (Corning, no. 8162), and the RNA was precipitated with ethanol in the presence of sodium acetate and GlycoBlue Coprecipitant (Thermo Fisher Scientific, no. AM9516).

The size-fractionated RNA was ligated to a 3′-randomized adapter at 25 °C for 16 h in a reaction mixture containing 0.25 µM 3′-randomized adapter (HO001), 200 U T4 RNA ligase 2 truncated KQ (NEB, no. M0373), 1× T4 RNA ligase reaction buffer (NEB), 20% PEG 8000 (NEB) and 10 U SUPERase•In RNase Inhibitor (Thermo Fisher Scientific, no. AM2694). The adapter-ligated RNA (40–55 nt) was size-fractionated by 15% denaturing polyacrylamide gel electrophoresis to remove unligated adapters and gel-purified as described above. The 3′-adapter-ligated RNA was then ligated to a 0.18 µM 5′-randomized adapter (HO002) with 14 U T4 RNA ligase 1 (NEB, no. M0204), 1× T4 RNA ligase reaction buffer, 20% PEG 8000, 1 mM ATP and 14 U SUPERase•In RNase Inhibitor at 37 °C for 1 h, followed by heat inactivation at 65 °C for 15 min.

The adapter-ligated RNA was reverse-transcribed using 200 U SuperScript III reverse transcriptase (Thermo Fisher Scientific, no. 18080044), 1× first-strand buffer (Thermo Fisher Scientific), 0.2 µM RT primer (HO003), 0.5 mM dNTPs (included in the TruSeq Small RNA Kit; Illumina) and 5 mM DTT (Thermo Fisher Scientific) at 50 °C for 1 h. Reverse transcriptase was heat-inactivated at 70 °C for 15 min after reverse transcription. The cDNA library was amplified using 1 U Phusion High-Fidelity DNA Polymerase (Thermo Fisher Scientific, no. F530S), 1× Phusion HF buffer (Thermo Fisher Scientific), 0.5 µM primers (RP1 forward primer and RPIX reverse primer from the TruSeq Small RNA Kit; Illumina) and 0.2 mM dNTPs (Takara Bio).

The amplified cDNA library was size-fractionated (145–160 nt) on a 6% non-denaturing polyacrylamide gel to remove adapter dimers. The size-fractionated cDNA library was gel-purified and precipitated as described above. The size of the cDNA library was evaluated using an S1 Cartridge (BiOptic, no. C105202) on a Qsep100 DNA Fragment Analyzer, and the concentration of each library was quantified using KAPA Library Quantification Kits (KAPA Biosystems, no. KK4873). The concentration of the pooled cDNA library was quantified using Qubit 1X dsDNA High Sensitivity Assay Kits (Thermo Fisher Scientific, no. Q33230). The pooled cDNA library was sequenced on a NextSeq 500 platform (Illumina) using a NextSeq 500/550 High Output Kit v2.5 (75 cycles; Illumina).

#### 12.2. Preparation of amplicon sequencing libraries from genome-wide cgRNA screens

miRNA target sequences from genome-integrated cgRNAs were amplified from genomic DNA isolated from EGFP+ and unsorted mESC and SMC populations generated in the genome-wide cgRNA screens. Sequencing libraries were prepared by two successive rounds of PCR.

##### 12.2.1. First-round amplification of miRNA targets from genome-integrated cgRNAs

miRNA target regions from genome-integrated cgRNAs were amplified by PCR from genomic DNA collected from EGFP+ and unsorted mESCs or SMCs transduced with the cgRNA library. In the first round of PCR, the miRNA target region was amplified from 50 µl of genomic DNA divided between two 50-µl reactions using a pool of staggered-length forward primers (AA112–AA118) and a single reverse primer (AA119), under the reaction and thermocycling conditions described in **Supplementary Note 2**. The first-round PCR products were then purified by column purification using the GeneJET Gel Extraction Kit (Thermo Fisher Scientific, no. K0692) according to the manufacturer’s protocol and eluted in 35 µl of UltraPure Distilled Water.

##### 12.2.2. Addition of Illumina adapters and sample indexes

In the second round of PCR, 25 µl of the purified first-round PCR product was amplified in a single 50-µl reaction under the PCR reaction and thermocycling conditions described in **Supplementary Note 2**. First-round PCR products from the mESC genome-wide screen (Section 11.1) were amplified using a common Illumina P5 adapter primer and custom i7-indexed Illumina P7 adapter primers (**Table S6**). First-round PCR products from the SMCs and mESCs in the SMC genome-wide screen (Section 11.2) were amplified using custom i5-indexed Illumina P5 adapter primers and custom i7-indexed Illumina P7 adapter primers (**Table S6**).

Second-round PCR products were size-selected by electrophoresis on a 2% agarose gel prepared in 1× Tris-borate-EDTA buffer using agarose from Thermo Fisher Scientific (no. 16500500). The products were purified by gel extraction using the GeneJET Gel Extraction Kit according to the manufacturer’s protocol, and the concentrations of the purified libraries were measured using a Qubit fluorometer (Thermo Fisher Scientific) prior to qPCR quantification.

The libraries were then quantified by qPCR using the NEBNext Library Quant Kit for Illumina (NEB, no. E7630S). Library concentrations were calculated based on the mean length of the expected amplicons (502 bp), and the libraries were pooled in equimolar amounts. For the mESC genome-wide screen (Section 11.1), the pooled sequencing library was sequenced on an Illumina NextSeq system using a NextSeq P1 100 150-cycle kit, with a 110-cycle single-end Read 1 and an 8-cycle index read for the i7 index. For the SMC genome-wide screen (Section 11.2), the same sequencer and sequencing kit were used, with a 120-cycle single-end Read 1, an 8-cycle index read for the i7 index and an 8-cycle index read for the i5 index. PhiX DNA was spiked into each run at 30% to improve sequence diversity on the flow cell.

### 13. Sequencing and computational analysis

NextSeq outputs generated BCL files, which were demultiplexed based on single or paired indexes and converted to FASTQ files using bcl2fastq2. All analysis scripts referenced in the following sections, together with their outputs, are available at https://github.com/yachielab/cgRNA.

#### 13.1. Small RNA sequencing analysis

For AQ-seq reads, the 3′ adapter sequence and 4-nt degenerate sequences at the 5′ and 3′ ends were trimmed using cutadapt^141^. Reads shorter than 18 nt were removed. Low-quality reads, defined as reads with a Phred quality score of 20 or below for more than 95% of bases or 30 or below for more than 50% of bases, as well as artifact reads, were removed using FASTX-Toolkit (http://hannonlab.cshl.edu/fastx_toolkit/). The processed reads were analyzed using the miRge2.0 pipeline^142^ to obtain miRNA read counts.

#### 13.2. Analysis of the genome-wide cgRNA screen in mESCs

##### 13.2.1. Counting miRNA target reads

miRNA target reads were counted from raw sequencing reads in Python using a modified version of a previously described barcode-counting script^143^ (cgLibV2_BC_counter.ipynb). The list of synthesized targets in the cgRNA library (**Table S2**) was used to generate a whitelist for the counting script. Adjacent bases from the 3′ end of the gRNA scaffold were appended to each target sequence such that all reference sequences were 28 nt in length before being used for target counting (mmuReferenceList.ipynb). Raw target counts were then compiled into a meta-label (replicate or condition)–miRNA target count table using multiClonal_cgRNA_matrix.ipynb (**Table S7**) for differential enrichment analysis using DESeq2 in R (**Table S3**)^82^.

##### 13.2.2. Counting miRNA target reads and gene set enrichment analysis

Enrichment of each miRNA target in EGFP+ cells relative to unsorted cells was analyzed using DESeq2. For each target, the mean log_2_-transformed fold enrichment in EGFP+ cells relative to unsorted cells was calculated, together with a Benjamini–Hochberg-adjusted *P* value to control the false discovery rate (FDR). A miRNA target was classified as an active hit when its log_2_ fold enrichment was greater than 1 and its FDR was less than 0.05.

Active miRNA hits were subjected to gene set enrichment analysis using miEAA^83^ with the *Mus musculus* library. Predicted gene targets associated with the miRNA hits were analyzed within each gene-set term category. Terms were considered significant at a Benjamini–Hochberg FDR threshold of 0.05 and were required to be associated with at least two miRNA hits.

##### 13.2.3. Classification of paired miRNA strands by cleavage activity

Mature miRNA targets derived from opposite arms of the same pre-miRNA hairpin were paired. Pairs were classified as homoactive when both strands met the active-hit criteria, heteroactive when only one strand met the criteria, and homoinactive when neither strand met the criteria.

Screen enrichment and AQ-seq expression were compared among active strands from homoactive pairs, active and inactive strands from heteroactive pairs, and inactive strands from homoinactive pairs. AQ-seq expression was analyzed both including and excluding miRNAs with zero read counts. Differences among activity categories were assessed using two-sided Mann–Whitney *U*-tests for unpaired comparisons.

##### 13.2.4. Sequence-based assignment of loading and passenger motifs to miRNA strands

A simplified sequence-scoring heuristic was used to classify each mature miRNA strand according to its resemblance to the loading motif 5′-UR_n_ or the passenger motif 5′-CY_n_, based on previously reported nucleotide-composition biases associated with miRNA strand selection^86^. Here, R denotes a purine and Y denotes a pyrimidine. These classifications represent sequence-based predictions and do not directly measure Argonaute loading.

For each mature miRNA strand, loading- and passenger-motif scores were calculated independently. For the loading-motif score, a 5′ uridine contributed 5.0 points, each purine at subsequent positions contributed 0.5 points, and a 5′ cytosine contributed –2.5 points. Conversely, for the passenger-motif score, a 5′ cytosine contributed 5.0 points, each pyrimidine at subsequent positions contributed 0.5 points, and a 5′ uridine contributed –2.5 points.

Using a threshold of 5.0 points, a strand was assigned the loading motif when its loading-motif score exceeded the threshold and its passenger-motif score fell below it. Similarly, a strand was assigned the passenger motif when its passenger-motif score exceeded the threshold and its loading-motif score fell below it. All other cases, including those with equal loading- and passenger-motif scores, were classified as ambiguous. Motif scores and assignments were calculated using importHits2File.Rmd and riscLoadingModel.ipynb (**Table S8**). Paired-strand analyses were restricted to miRNA pairs for which both strands received unambiguous motif assignments.

##### 13.2.5. Paired-strand motif analysis

Motif assignments were summarized according to strand activity and miRNA-pair class using ago2ModelResults.Rmd. For heteroactive pairs, conditional motif frequencies were calculated to assess whether the motif assignment of one strand depended on the activity status and motif assignment of the opposite strand from the same pre-miRNA.

For each comparison, one member of the pair was designated the focusing strand and the other the paired strand. The probability that the focusing strand carried a loading or passenger motif was calculated conditional on the activity status of the focusing strand and the motif assignment and activity status of the paired strand. To account for the overall excess of loading-motif assignments, each conditional probability was divided by the corresponding global frequency of that motif among all unambiguously assigned strands from heteroactive pairs. Values greater than 1 indicated enrichment relative to the global motif-assignment frequency, whereas values below 1 indicated depletion.

Heteroactive pairs were also grouped according to the combination of motif assignments carried by their active and inactive strands. AQ-seq expression values of the active and inactive strands within each group were compared using a two-sided Wilcoxon signed-rank test.

#### 13.3. Analysis of genome-wide cgRNA recording during SMC differentiation

miRNA target counts from mESCs and from day 3 and day 6 of SMC differentiation were generated from raw sequencing reads as described in Section 13.2, using cgLibV2_BC_SMC_counter.ipynb for miRNA target counting and SMC_cgRNA_matrix.ipynb for count-matrix generation.

For each recording condition, differential representation between EGFP+ and unsorted populations was analyzed independently using DESeq2. Active hits were defined as targets with a log_2_ fold enrichment greater than 1 and a Benjamini–Hochberg-adjusted *P* value below 0.05.

Targets classified as active in at least one condition were compiled into a unified table containing the enrichment value and adjusted *P* value for all three recording conditions. Targets were grouped by hierarchical clustering of their enrichment profiles. Shared and condition-specific active hits were identified and visualized using eulerr^144,145^.

### 14. Mouse engineering

#### 14.1. *In vitro* fertilization and zygote cryopreservation

Cumulus–oocyte complexes (COCs) were collected from 5-to 6-week-old B6D2F1 female mice superovulated by sequential intraperitoneal injections of 0.15 ml CARD HyperOva (Kyudo, no. F021) and 6.5 IU human chorionic gonadotropin (hCG; GONATROPIN 3000, ASKA Animal Health) administered 48 h apart. Sperm obtained from 12-week-old B6D2F1 males were preincubated in TYH medium at 37 °C under 5% CO_2_ for 2 h. COCs were then inseminated with sperm at a final concentration of 2 × 10^5^ sperm ml^−1^. At 3 h after insemination, COCs were treated with 0.33 mg ml^−1^ hyaluronidase at 37 °C in air for 10 min and washed in fresh KSOM to completely remove cumulus cells. Cumulus-free oocytes were then cultured in KSOM at 37 °C under 5% CO_2_ for an additional 3 h, after which two-pronuclear zygotes were selected for cryopreservation.

Zygotes were washed three times at room temperature in PB1 medium containing 1 M DMSO. Up to 80 zygotes in 5 µl of medium were transferred directly into a cryogenic vial and incubated at 0 °C for 5 min, followed by the addition of 45 µl of ice-cold DAP213 (ARK-Resource, no. I0BAI1100) and incubation at 0 °C for an additional 5 min. Cryovials were then mounted on cryogenic canes covered with plastic sleeves and gently immersed in liquid nitrogen for long-term storage.

#### 14.2. Synthesis and purification of piggyBac transposase mRNA

Hyperactive piggyBac transposase (hyPBase) mRNA was synthesized using the mMESSAGE mMACHINE T7 Transcription Kit (Thermo Fisher Scientific, no. AM1345), followed by addition of a poly(A) tail using the Poly(A) Tailing Kit (Thermo Fisher Scientific, no. AM1350). *In vitro*-transcribed mRNA was purified by phenol–chloroform extraction and ethanol precipitation, desalted using MicroSpin G-25 Columns (Cytiva, no. 27532501), diluted to 600 ng µl^−1^ in nuclease-free water, aliquoted in 2-µl volumes into 0.5-ml DNA LoBind Tubes (Eppendorf, no. EP0030122348), and stored at –80 °C for no longer than 3 months.

#### 14.3. Zygote thawing and electroporation

A cryovial was placed at room temperature with the cap removed for 1.5 min. Next, 0.9 ml of 0.25 M sucrose in PB1 was added, and the contents were gently mixed by pipetting 10 times. Zygotes were immediately recovered and washed in fresh KSOM to remove the cryoprotectant. A 2-µl aliquot of hyPBase mRNA was thawed at room temperature and diluted with 38 µl of Opti-MEM to a final concentration of 30 ng µl^−1^. The resulting 40-µl mRNA solution was loaded into the electroporation chamber of a CUY520P5 petri dish (NEPA GENE), and the impedance was confirmed to be between 490 and 590 Ω. After incubation at 37 °C under 5% CO_2_ for 30 min, zygotes were washed three times in fresh Opti-MEM, transferred to the electroporation chamber, and aligned in two parallel rows perpendicular to the direction of the electroporation pulse. Following electroporation, zygotes were washed three times in KSOM and cultured at 37 °C under 5% CO_2_ for 3 h before microinjection.

#### 14.4. Plasmid purification, pronuclear microinjection, and embryo transfer

Before microinjection, piggyBac plasmids were freshly purified by phenol–chloroform extraction and ethanol precipitation and passed through Ultrafree-MC Centrifugal Filters (Millipore Sigma, no. UFC30GV0S). Plasmids were then diluted to 6 ng µl^−1^ in T_10_E_0.1_ buffer. The base-editing reporter plasmid (pAA109) was mixed with an equal volume of either an sgRNA/Target-AIDmax plasmid containing three introns (pAA111) or a cgRNA/Target-AIDmax plasmid containing three introns (pAA110).

Holding and injection capillaries were freshly prepared using a P-1000 Micropipette Puller (Sutter Instrument) and bent to 20° using an MF-3 microforge (Narishige). Electroporated zygotes were washed three times in FHM and transferred to an FHM drop covered with mineral oil. The male pronucleus was injected with the plasmid mixture at room temperature using a Narishige micromanipulation system equipped with a manual CellTram 4r Air microinjector (Eppendorf) for the holding pipette and a semiautomatic FemtoJet 4i microinjector (Eppendorf) for the injection pipette. Following microinjection, zygotes were washed three times in KSOM and cultured overnight at 37 °C under 5% CO_2_.

The following day, embryos that had developed to the 2-cell stage were transferred into the oviductal ampullae of 0.5-dpc pseudopregnant ICR females (Japan SLC).

#### 14.5. Embryo extraction and imaging

E17.5 embryos were dissected from the uteri of pseudopregnant females and kept on ice. The abdominal wall was removed to expose the internal organs. Fluorescence images were captured using a Keyence BZ-X800 microscope.

### 15. Detailed molecular cloning methods

Unless otherwise specified, PCR primers and synthetic oligonucleotides are listed in **Table S6**, and plasmids are listed in **Table S4**. The common procedures described in the Supplementary Notes were used throughout the construction of individual plasmids. Plasmid-construction-specific templates, primers, restriction enzymes, annealing temperatures, extension times, fragment ratios and deviations are specified in the corresponding subsections.

#### 15.1. MICR gRNAs

MICR gRNAs were produced by PCR amplification, double restriction enzyme digestion and ligation. First, 15 µl of the PB4-CAGGS-17T-MICR backbone was digested with EcoRI-HF (NEB, no. R3101L) and BamHI-HF (NEB, no. R3136L) for 2 h at 37 °C using the common restriction enzyme digestion protocol described in **Supplementary Note 1**. The backbone band was size-selected by electrophoresis on a 1% agarose gel and gel-purified using the GeneJET Gel Extraction Kit according to the manufacturer’s protocol.

MICR gRNA fragments were amplified from PB4-CAGGS-17T-MICR using the primer pairs listed in **Table S6** and the Phusion PCR protocol described in **Supplementary Note 1**, with an annealing temperature of 64 °C and an extension time of 30 s. Amplified fragments were size-selected by electrophoresis on a 2% agarose gel and gel-purified using the GeneJET Gel Extraction Kit according to the manufacturer’s protocol.

The gel-purified MICR gRNA PCR products were then digested with EcoRI-HF and BamHI-HF for 1 h at 37 °C using the common restriction enzyme digestion protocol described in **Supplementary Note 1**, PCR-purified using the GeneJET Gel Extraction Kit according to the manufacturer’s protocol, and ligated between the EcoRI and BamHI sites of the digested PB4-CAGGS-17T-MICR backbone at a 3:1 insert-to-backbone molar ratio using T4 DNA Ligase (NEB, no. M0202L) in 1× T4 DNA Ligase Buffer with ATP (NEB, no. B0202L) at 25 °C for 10 min.

#### 15.2. cgRNAs and sgRNAs

cgRNAs and sgRNAs were generated using the common Golden Gate assembly protocol described in **Supplementary Note 3**, with DNA oligonucleotide pairs and backbones listed in **Table S6**.

#### 15.3. cgRNA library

The protocols for amplifying miRNA targets from a Twist Bioscience-synthesized oligonucleotide pool and inserting the amplified targets into a cgRNA backbone are described in detail in **Supplementary Note 4**.

#### 15.4. piggyBac expression vectors for gRNA units and early Target-AID reporter configuration

##### 15.4.1. PB4 cgRNA backbones

*cgRNA V1.* To produce PB4-cgRNA-V1 (cloning backbone), the PB4-CAGGS-17T-MICR backbone was first digested with KpnI-HF (NEB, no. R3142L) and BamHI-HF for 2 h at 37 °C using the common restriction enzyme digestion protocol described in **Supplementary Note 1**. The backbone band was size-selected by electrophoresis on a 1% agarose gel and gel-purified using the GeneJET Gel Extraction Kit according to the manufacturer’s protocol. The hU6-cgRNA-V1 transcriptional unit was amplified from a double-stranded DNA gene block purchased from Twist Bioscience (GID001) using AA001/AA002 and the Phusion PCR protocol described in **Supplementary Note 1**, with an annealing temperature of 66 °C and an extension time of 30 s. The amplified fragment was size-selected by electrophoresis on a 2% agarose gel and gel-purified using the GeneJET Gel Extraction Kit according to the manufacturer’s protocol. The gel-purified hU6-cgRNA-V1 PCR product was then digested with KpnI-HF and BamHI-HF for 1 h at 37 °C using the common restriction enzyme digestion protocol described in **Supplementary Note 1**, PCR-purified using the GeneJET Gel Extraction Kit according to the manufacturer’s protocol, and ligated between the KpnI and BamHI sites of the digested PB4-CAGGS-17T-MICR backbone at a 3:1 insert-to-backbone molar ratio using T4 DNA Ligase in 1× T4 DNA Ligase Buffer with ATP at 25 °C for 10 min.

*cgRNA V2.* To produce PB4-cgRNA-V2 (cloning backbone), the PB4-cgRNA-V1 cloning backbone was first digested with MluI-HF (NEB, no. R3198L) and BamHI-HF for 2 h at 37 °C using the common restriction enzyme digestion protocol described in **Supplementary Note 1**. The backbone band was size-selected by electrophoresis on a 1% agarose gel and gel-purified using the GeneJET Gel Extraction Kit according to the manufacturer’s protocol. The hU6-5′-ribozyme and cgRNA-3′-ribozyme fragments were amplified from double-stranded DNA gene blocks purchased from Twist Bioscience (GID002 and GID003) using AA001/AA003 and AA004/AA005, respectively, and the Phusion PCR protocol described in **Supplementary Note 1**, with an annealing temperature of 64 °C and an ET of 30 s. The amplified fragments were size-selected by electrophoresis on a 2% agarose gel and gel-purified using the GeneJET Gel Extraction Kit according to the manufacturer’s protocol. The purified hU6-5′-ribozyme and cgRNA-3′-ribozyme fragments and the digested and purified PB4-cgRNA-V1 backbone were then mixed at a molar ratio of 3:3:1 (insert:insert:backbone) and assembled by Gibson assembly at 50 °C for 1 h.

*cgRNA V3.* To produce PB4-cgRNA-V3 (cloning backbone), the PB4-cgRNA-V1 cloning backbone was first digested with MluI-HF and BbsI-HF (NEB, no. R3539L) for 2 h at 37 °C using the common restriction enzyme digestion protocol described in **Supplementary Note 1**. The backbone band was size-selected by electrophoresis on a 1% agarose gel and gel-purified using the GeneJET Gel Extraction Kit according to the manufacturer’s protocol. The hU6 promoter was amplified from PB4-cgRNA-V1 using AA001/AA006, and the cgRNA-V3 scaffold and 3′-bGH poly(A) signal fragments were amplified from PB4-CAGGS-17T-MICR using AA007/AA008 and AA009/AA010, respectively, using the Phusion PCR protocol described in **Supplementary Note 1**, with an annealing temperature of 61 °C and an extension time of 30 s. The amplified fragments were size-selected by electrophoresis on a 2% agarose gel and gel-purified using the GeneJET Gel Extraction Kit according to the manufacturer’s protocol. The purified hU6 promoter, cgRNA-V3 scaffold and 3′-bGH poly(A) signal fragments and the digested and purified PB4-cgRNA-V1 backbone were then mixed at a molar ratio of 3:3:3:1 (insert:insert:insert:backbone) and assembled by Gibson assembly at 50 °C for 1 h.

##### 15.4.2. PB4 sgRNA

To produce PB4-US1-sgRNA and PB4-Scr-sgRNA, the same PB4-cgRNA-V1 cloning-backbone digest used to produce PB4-cgRNA-V2 was used for assembly. The hU6-US1-sgRNA and hU6-Scr-sgRNA fragments were generated by fusion PCR. The hU6-US1 and US1-sgRNA fragments and the hU6-Scr and Scr-sgRNA fragments were first amplified from a gene block synthesized by Twist Bioscience (GID004) using AA001/AA011 and AA012/AA013, and AA001/AA014 and AA015/AA013, respectively, using the Phusion PCR protocol described in **Supplementary Note 1**, with an annealing temperature of 61 °C and an extension time of 30 s. The amplified fragments were size-selected by electrophoresis on a 2% agarose gel and gel-purified using the GeneJET Gel Extraction Kit according to the manufacturer’s protocol.

The purified hU6-US1 and US1-sgRNA fragments were mixed at equal volumes (5 µl each), and 1 µl of the mixture was amplified using AA001/AA013 and the Phusion PCR protocol described in **Supplementary Note 1**, with an annealing temperature of 65 °C and an extension time of 30 s. The resulting hU6-US1-sgRNA fusion PCR product was size-selected by electrophoresis on a 2% agarose gel and gel-purified using the GeneJET Gel Extraction Kit according to the manufacturer’s protocol. The same procedure was performed with the hU6-Scr and Scr-sgRNA fragments to generate the hU6-Scr-sgRNA fragment.

The gel-purified hU6-US1-sgRNA and hU6-Scr-sgRNA PCR products were then digested with MluI-HF and BamHI-HF for 1 h at 37 °C using the common restriction enzyme digestion protocol described in **Supplementary Note 1**, PCR-purified using the GeneJET Gel Extraction Kit according to the manufacturer’s protocol, and ligated between the MluI and BamHI sites of the digested PB4-cgRNA-V1 backbone at a 3:1 insert-to-backbone molar ratio using T4 DNA Ligase in 1× T4 DNA Ligase Buffer with ATP at 25 °C for 10 min.

##### 15.4.3. Single piggyBac vector for base-editing reporter and Target-AID

To produce PB-CAG-US1-eGFP-EF1-Target-AID, the pRS290 backbone plasmid was first digested with MfeI-HF (NEB, no. R3589L) and AvrII (NEB, no. R0174L) for 2 h at 37 °C using the common restriction enzyme digestion protocol described in **Supplementary Note 1**. The backbone band was size-selected by electrophoresis on a 1% agarose gel and gel-purified using the GeneJET Gel Extraction Kit according to the manufacturer’s protocol. The full-length EF1a promoter sequence was amplified from pSI-172-pcDNA3.1-EF1a using AA016/AA017 and the Phusion PCR protocol described in **Supplementary Note 1**, with an annealing temperature of 62 °C and an extension time of 60 s. The amplified fragment was size-selected by electrophoresis on a 2% agarose gel and gel-purified using the GeneJET Gel Extraction Kit according to the manufacturer’s protocol. The gel-purified full-length EF1a promoter PCR product was then digested with MfeI-HF and AvrII for 1 h at 37 °C using the common restriction enzyme digestion protocol described in **Supplementary Note 1**, PCR-purified using the GeneJET Gel Extraction Kit according to the manufacturer’s protocol, and ligated between the MfeI and AvrII sites of the digested pRS290 backbone at a 3:1 insert-to-backbone molar ratio using T4 DNA Ligase in 1× T4 DNA Ligase Buffer with ATP at 25 °C for 10 min.

The resulting intermediate vector was confirmed by Sanger sequencing for correct insertion and then digested with NotI-HF (NEB, no. R3189L) and AgeI-HF (NEB, no. R3552L) for 2 h at 37 °C using the common restriction enzyme digestion protocol described in **Supplementary Note 1**. The backbone band was size-selected by electrophoresis on a 1% agarose gel and gel-purified using the GeneJET Gel Extraction Kit according to the manufacturer’s protocol. The mutated EGFP fragment containing the US1 spacer target (US1-EGFP) was amplified from pRS290 using AA018/AA019 and the Phusion PCR protocol described in **Supplementary Note 1**, with an annealing temperature of 65 °C and an extension time of 30 s. The amplified fragment was size-selected by electrophoresis on a 1% agarose gel and gel-purified using the GeneJET Gel Extraction Kit according to the manufacturer’s protocol. The gel-purified US1-EGFP PCR product was then digested with NotI-HF and AgeI-HF for 1 h at 37 °C using the common restriction enzyme digestion protocol described in **Supplementary Note 1**, PCR-purified using the GeneJET Gel Extraction Kit according to the manufacturer’s protocol, and ligated between the NotI and AgeI sites of the digested intermediate vector at a 3:1 insert-to-backbone molar ratio using T4 DNA Ligase in 1× T4 DNA Ligase Buffer with ATP at 25 °C for 10 min.

To produce additional base-editing reporter targets, PB-CAG-US1-eGFP-EF1-Target-AID was first digested with NotI-HF (NEB, no. R3189L) and AgeI-HF (NEB, no. R3552L) for 2 h at 37 °C using the common restriction enzyme digestion protocol described in **Supplementary Note 1**. The backbone band was size-selected by electrophoresis on a 1% agarose gel and gel-purified using the GeneJET Gel Extraction Kit according to the manufacturer’s protocol. Target-EGFP fragments were generated by amplifying the start-codon-mutated EGFP coding sequence from PB-CAG-US1-eGFP-EF1-Target-AID using the target-containing forward primers listed in **Table S6** and the common reverse primer AA019, according to the Phusion PCR protocol described in **Supplementary Note 1**, with an annealing temperature of 65 °C and an extension time of 30 s. The amplified fragments were size-selected by electrophoresis on a 1% agarose gel and gel-purified using the GeneJET Gel Extraction Kit according to the manufacturer’s protocol. The gel-purified Target-EGFP PCR products were then digested with NotI-HF and AgeI-HF for 1 h at 37 °C using the common restriction enzyme digestion protocol described in **Supplementary Note 1**, PCR-purified using the GeneJET Gel Extraction Kit according to the manufacturer’s protocol, and ligated between the NotI and AgeI sites of the digested PB-CAG-US1-eGFP-EF1-Target-AID backbone at a 3:1 insert-to-backbone molar ratio using T4 DNA Ligase in 1× T4 DNA Ligase Buffer with ATP at 25 °C for 10 min.

##### 15.4.4. piggyBac vectors expressing mCherry and cgRNA V3 or sgRNA controls

To produce PB-CAG-mCherry-cgRNA-V3 (cloning backbone) and PB-CAG-sgRNA-BB, the PB-CAG-US1-eGFP-EF1-Target-AID backbone was first digested with NotI-HF and AgeI-HF for 2 h at 37 °C using the common restriction enzyme digestion protocol described in **Supplementary Note 1**. The backbone band was size-selected by electrophoresis on a 1% agarose gel and gel-purified using the GeneJET Gel Extraction Kit according to the manufacturer’s protocol. The mCherry fragment was amplified from pSI-173-CMVp-MCS-mCherry using AA020/AA021 and the Phusion PCR protocol described in **Supplementary Note 1**, with an annealing temperature of 65 °C and an extension time of 30 s. The amplified fragment was size-selected by electrophoresis on a 1% agarose gel and gel-purified using the GeneJET Gel Extraction Kit according to the manufacturer’s protocol. The gel-purified mCherry PCR product was then digested with NotI-HF and AgeI-HF for 1 h at 37 °C using the common restriction enzyme digestion protocol described in **Supplementary Note 1**, PCR-purified using the GeneJET Gel Extraction Kit according to the manufacturer’s protocol, and ligated between the NotI and AgeI sites of the digested PB-CAG-US1-eGFP-EF1-Target-AID backbone at a 3:1 insert-to-backbone molar ratio using T4 DNA Ligase in 1× T4 DNA Ligase Buffer with ATP at 25 °C for 10 min.

The resulting intermediate vector was confirmed by Sanger sequencing for correct insertion and then digested with MfeI-HF and ClaI (NEB, no. R0197L) for 2 h at 37 °C using the common restriction enzyme digestion protocol described in **Supplementary Note 1**. The backbone band was size-selected by electrophoresis on a 1% agarose gel and gel-purified using the GeneJET Gel Extraction Kit according to the manufacturer’s protocol. The hU6-cgRNA-V3 and hU6-sgRNA fragments were amplified from PB4-cgRNA-V3 and a gene block synthesized by Twist Bioscience (GID004), respectively, using AA022/AA023 and AA022/AA024, respectively, and the Phusion PCR protocol described in **Supplementary Note 1**, with an annealing temperature of 64 °C and an extension time of 30 s. The amplified fragments were size-selected by electrophoresis on a 1% agarose gel and gel-purified using the GeneJET Gel Extraction Kit according to the manufacturer’s protocol. The gel-purified hU6-cgRNA-V3 and hU6-sgRNA PCR products were then digested with MfeI-HF and ClaI for 1 h at 37 °C using the common restriction enzyme digestion protocol described in **Supplementary Note 1**, PCR-purified using the GeneJET Gel Extraction Kit according to the manufacturer’s protocol, and ligated between the MfeI and ClaI sites of the digested intermediate vector at a 3:1 insert-to-backbone molar ratio using T4 DNA Ligase in 1× T4 DNA Ligase Buffer with ATP at 25 °C for 10 min.

#### 15.5. Insulated piggyBac expression vectors for BE reporters, codon-optimized Cas9 editors and hAgo2

##### 15.5.1. piggyBac vectors expressing a base-editing reporter and G418 resistance gene

To produce PB-INS-CAG-US1-eGFP-EF1a-Neo, PB-U6insert-Puro (Addgene, no. 104537) was first digested with BamHI-HF and SalI-HF (NEB, no. R3138L) for 2 h at 37 °C using the common restriction enzyme digestion protocol described in **Supplementary Note 1**. The backbone band was size-selected by electrophoresis on a 1% agarose gel and gel-purified using the GeneJET Gel Extraction Kit according to the manufacturer’s protocol. The EF1-NeoR transcriptional unit was amplified from a double-stranded DNA gene block purchased from Twist Bioscience (GID005) using AA025/AA026 and the Phusion PCR protocol described in **Supplementary Note 1**, with an annealing temperature of 62 °C and an extension time of 60 s. The amplified fragment was size-selected by electrophoresis on a 1% agarose gel and gel-purified using the GeneJET Gel Extraction Kit according to the manufacturer’s protocol. The gel-purified EF1-NeoR PCR product was then digested with BamHI-HF and SalI-HF for 1 h at 37 °C using the common restriction enzyme digestion protocol described in **Supplementary Note 1**, PCR-purified using the GeneJET Gel Extraction Kit according to the manufacturer’s protocol, and ligated between the BamHI and SalII sites of the digested PB-U6insert-Puro backbone at a 3:1 insert-to-backbone molar ratio using T4 DNA Ligase in 1× T4 DNA Ligase Buffer with annealing temperatureP at 25 °C for 10 min.

The assembled product was confirmed by Sanger sequencing and then digested with SpeI-HF (NEB, no. R3133L) and AgeI-HF for 2 h at 37 °C using the common restriction enzyme digestion protocol described in **Supplementary Note 1**. The backbone band was size-selected by electrophoresis on a 1% agarose gel and gel-purified using the GeneJET Gel Extraction Kit according to the manufacturer’s protocol. To obtain the CAG-US1-EGFP transcriptional unit, PB-CAG-US1-EF1-Target-AID was digested with NheI-HF (NEB, no. R3131L) and AgeI-HF for 2 h at 37 °C using the common restriction enzyme digestion protocol described in **Supplementary Note 1**. The insert band was size-selected by electrophoresis on a 1% agarose gel and gel-purified using the GeneJET Gel Extraction Kit according to the manufacturer’s protocol. The digested CAG-US1-EGFP fragment was then ligated between the SpeI and AgeI sites of the digested assembly backbone at a 3:1 insert-to-backbone molar ratio using T4 DNA Ligase in 1× T4 DNA Ligase Buffer with ATP (NEB) at 25 °C for 10 min.

To produce PB-INS-CAG-US1-eGFP-EF1a-mCherry, PB-INS-CAG-US1-eGFP-EF1a-Neo was first digested with MfeI-HF and SalI-HF (NEB, no. R3138L) for 2 h at 37 °C using the common restriction enzyme digestion protocol described in **Supplementary Note 1**. The backbone band was size-selected by electrophoresis on a 1% agarose gel and gel-purified using the GeneJET Gel Extraction Kit according to the manufacturer’s protocol. The EF1-mCherry transcriptional unit was amplified from LVSIN-U6-cgRNA-V3-EF1a-mCherry (cloning backbone) using AA124/AA062 and the Phusion PCR protocol described in **Supplementary Note 1**, with an annealing temperature of 63 °C and an extension time of 60 s. The amplified fragment was size-selected by electrophoresis on a 1% agarose gel and gel-purified using the GeneJET Gel Extraction Kit according to the manufacturer’s protocol. The gel-purified EF1-mCherry PCR product was then digested with MfeI-HF and SalI-HF for 1 h at 37 °C using the common restriction enzyme digestion protocol described in **Supplementary Note 1**, PCR-purified using the GeneJET Gel Extraction Kit according to the manufacturer’s protocol, and ligated between the MfeI and SalII sites of the digested PB-INS-CAG-US1-eGFP-EF1a-mCherry backbone at a 3:1 insert-to-backbone molar ratio using T4 DNA Ligase in 1× T4 DNA Ligase Buffer with ATP at 25 °C for 10 min.

##### 15.5.2. piggyBac vector expressing codon-optimized Target-AID with mTagBFP2

To produce PB-INS-EF1a-Target-AID-T2A-mTagBFP2, PB-CAG-US1-eGFP-EF1-Target-AID was first digested with SpeI-HF and SalI-HF for 2 h at 37 °C using the common restriction enzyme digestion protocol described in **Supplementary Note 1**. The backbone band was size-selected by electrophoresis on a 1% agarose gel and gel-purified using the GeneJET Gel Extraction Kit according to the manufacturer’s protocol. The EF1a-nCas9, Cas9-pmCDA1-UGI-T2A and mTagBFP2 fragments were amplified with homologous overlapping regions from pNM1466, pNM1466 and dAAVS1_EF1a-BxB1_mTagBFP2, respectively, using AA027/AA028, AA029/AA030 and AA031/AA032, respectively, and the Phusion PCR protocol described in **Supplementary Note 1**, with an annealing temperature of 62 °C and an extension time of 150 s. The amplified fragments were size-selected by electrophoresis on a 1% agarose gel and gel-purified using the GeneJET Gel Extraction Kit according to the manufacturer’s protocol. The gel-purified EF1a-nCas9, Cas9-pmCDA1-UGI-T2A and mTagBFP2 fragments were then assembled into the restriction-enzyme-digested PB-CAG-US1-eGFP-EF1-Target-AID backbone by Gibson assembly at 50 °C for 1 h at a 2:2:1 molar ratio (insert:insert:backbone). The assembled product was confirmed by whole-plasmid sequencing.

To produce PB-INS-CAG-TargetAIDmax-nlsTagBFP2, PB-INS-CAG-US1-eGFP-EF1a-Neo was first digested with NotI-HF and SalI-HF for 2 h at 37 °C using the common restriction enzyme digestion protocol described in **Supplementary Note 1**. The backbone band was size-selected by electrophoresis on a 1% agarose gel and gel-purified using the GeneJET Gel Extraction Kit according to the manufacturer’s protocol. The Target-AIDmax and mTagBFP2 fragments were amplified with homologous overlapping regions from pCMV-Target-AIDmax and PB-INS-EF1a-Target-AID-T2A-mTagBFP2, respectively, using AA033/AA034 and AA035/AA032, respectively, and the Phusion PCR protocol described in **Supplementary Note 1**, with an annealing temperature of 62 °C and an extension time of 180 s. The amplified fragments were size-selected by electrophoresis on a 1% agarose gel and gel-purified using the GeneJET Gel Extraction Kit according to the manufacturer’s protocol. The gel-purified Target-AIDmax and mTagBFP2 fragments were then assembled into the restriction-enzyme-digested PB-INS-CAG-US1-eGFP-EF1a-Neo backbone by Gibson assembly at 50 °C for 1 h at a 2:2:1 molar ratio (insert:insert:backbone). The assembled product was confirmed by whole-plasmid sequencing.

##### 15.5.3. piggyBac vector expressing codon-optimized Target-AID containing endogenous mouse introns with mTagBFP2

Introns from the mouse genome were sequentially introduced into the codon-optimized Target-AID coding sequence. In the first step, PB-INS-CAG-TargetAIDmax-nlsTagBFP2 was digested with NotI-HF and PmlI-HF (NEB, no. R0532L) for 2 h at 37 °C using the common restriction enzyme digestion protocol described in **Supplementary Note 1**. The backbone band was size-selected by electrophoresis on a 1% agarose gel and gel-purified using the GeneJET Gel Extraction Kit according to the manufacturer’s protocol. The first intron of mouse *Gapdh* was amplified from 100 ng of mouse genomic DNA extracted from mESCs using AA036/AA037 and the Phusion PCR protocol described in **Supplementary Note 1**, with an annealing temperature of 64 °C and an extension time of 60 s. The amplified fragment was size-selected by electrophoresis on a 1% agarose gel and gel-purified using the GeneJET Gel Extraction Kit according to the manufacturer’s protocol. Two Target-AIDmax fragments containing homologous overlaps with the 5′ and 3′ ends of *Gapdh* intron 1 were amplified from PB-INS-CAG-TargetAIDmax-nlsTagBFP2 using AA033/AA038 and AA039/AA040, respectively, and the Phusion PCR protocol described in **Supplementary Note 1**, with an annealing temperature of 66 °C and an extension time of 120 s. The amplified fragments were size-selected by electrophoresis on a 1% agarose gel and gel-purified using the GeneJET Gel Extraction Kit according to the manufacturer’s protocol. The gel-purified 5′ Target-AIDmax, *Gapdh* intron 1 and 3′ Target-AIDmax fragments were then assembled into the restriction-enzyme-digested PB-INS-CAG-TargetAIDmax-nlsTagBFP2 backbone by Gibson assembly at 50 °C for 1 h at a 10:3:3:1 molar ratio (insert:insert:insert:backbone). The assembled product was confirmed by whole-plasmid sequencing before proceeding to the next assembly step.

To assemble PB-INS-CAG-TargetAIDmax-nlsTagBFP2 (two-introns), the one-intron assembly product from the previous step was digested with NotI-HF and PmlI-HF for 2 h at 37 °C using the common restriction enzyme digestion protocol described in **Supplementary Note 1**. The backbone band was size-selected by electrophoresis on a 1% agarose gel and gel-purified using the GeneJET Gel Extraction Kit according to the manufacturer’s protocol. The third intron of mouse *Actb* was amplified from 100 ng of mouse genomic DNA extracted from mESCs using AA041/AA042 and the Phusion PCR protocol described in **Supplementary Note 1**, with an annealing temperature of 65 °C and an extension time of 60 s. The amplified fragment was size-selected by electrophoresis on a 1% agarose gel and gel-purified using the GeneJET Gel Extraction Kit according to the manufacturer’s protocol. Two Target-AIDmax fragments containing homologous overlaps with the 5′ and 3′ ends of *Actb* intron 3 were amplified from the Intron+1 product using AA033/AA043 and AA044/AA040, respectively, and the Phusion PCR protocol described in **Supplementary Note 1**, with an annealing temperature of 64 °C and an extension time of 120 s. The amplified fragments were size-selected by electrophoresis on a 1% agarose gel and gel-purified using the GeneJET Gel Extraction Kit according to the manufacturer’s protocol. The gel-purified 5′ Target-AIDmax, *Actb* intron 3 and 3′ Target-AIDmax fragments were then assembled into the restriction-enzyme-digested one-intron backbone by Gibson assembly at 50 °C for 1 h at a 3:5:3:1 molar ratio (insert:insert:insert:backbone). The assembled product was confirmed by whole-plasmid sequencing.

To assemble PB-INS-CAG-TargetAIDmax-nlsTagBFP2 (three-introns), PB-INS-CAG-TargetAIDmax-nlsTagBFP2 (two-introns) was digested with NotI-HF and PmlI-HF for 2 h at 37 °C using the common restriction enzyme digestion protocol described in **Supplementary Note 1**. The backbone band was size-selected by electrophoresis on a 1% agarose gel and gel-purified using the GeneJET Gel Extraction Kit according to the manufacturer’s protocol. The first intron of mouse *Tuba* was amplified from 100 ng of mouse genomic DNA extracted from mESCs using AA045/AA046 and the Phusion PCR protocol described in **Supplementary Note 1**, with an annealing temperature of 65 °C and an extension time of 60 s. The amplified fragment was size-selected by electrophoresis on a 1% agarose gel and gel-purified using the GeneJET Gel Extraction Kit according to the manufacturer’s protocol. Two Target-AIDmax fragments containing homologous overlaps with the 5′ and 3′ ends of *Tuba* intron 1 were amplified from TargetAIDmax-nlsTagBFP2 (Intron+2) using AA033/AA047 and AA048/AA040, respectively, and the Phusion PCR protocol described in **Supplementary Note 1**, with an annealing temperature of 64 °C and an extension time of 120 s. The amplified fragments were size-selected by electrophoresis on a 1% agarose gel and gel-purified using the GeneJET Gel Extraction Kit according to the manufacturer’s protocol. The gel-purified 5′ Target-AIDmax, *Tuba* intron 3 and 3′ Target-AIDmax fragments were then assembled into the restriction-enzyme-digested PB-INS-CAG-TargetAIDmax-nlsTagBFP2 (two-intron) backbone by Gibson assembly at 50 °C for 1 h at a 3:3:3:1 molar ratio (insert:insert:insert:backbone). The assembled product was confirmed by whole-plasmid sequencing.

##### 15.5.4. piggyBac vectors expressing codon-optimized wild-type SpCas9 or SaCas9 with mTagBFP2

*SpCas9.* To produce PB-INS-CAG-spCas9-nlsTagBFP2, PB-INS-CAG-TargetAIDmax-nlsTagBFP2 was first digested with 1 µl of SalI-HF for 2 h at 37 °C under the common restriction enzyme digestion conditions described in **Supplementary Note 1**. The backbone band was size-selected by electrophoresis on a 1% agarose gel and gel-purified using the GeneJET Gel Extraction Kit according to the manufacturer’s protocol. The SpCas9 and mTagBFP2 fragments were amplified with homologous overlapping regions from PB-INS-CAG-TargetAIDmax-nlsTagBFP2 using AA049/AA050 and AA051/AA032, respectively, and the Phusion PCR protocol described in **Supplementary Note 1**, with an annealing temperature of 62 °C and an extension time of 180 s. The amplified fragments were size-selected by electrophoresis on a 1% agarose gel and gel-purified using the GeneJET Gel Extraction Kit according to the manufacturer’s protocol. The gel-purified SpCas9 and mTagBFP2 fragments were then assembled into the restriction-enzyme-digested PB-INS-CAG-TargetAIDmax-nlsTagBFP2 backbone by Gibson assembly at 50 °C for 1 h at a 2:2:1 molar ratio (insert:insert:backbone). The assembled product was confirmed by whole-plasmid sequencing.

*SaCas9.* To produce PB-INS-CAG-saCas9-nlsTagBFP2, PB-INS-CAG-US1-eGFP-EF1a-Neo was first digested with NotI-HF and SalI-HF (NEB) for 2 h at 37 °C using the common restriction enzyme digestion protocol described in **Supplementary Note 1**. The backbone band was size-selected by electrophoresis on a 1% agarose gel and gel-purified using the GeneJET Gel Extraction Kit according to the manufacturer’s protocol. The SaCas9 and mTagBFP2 fragments were amplified with homologous overlapping regions from pX601-AAV-CMV::NLS-SaCas9-NLS-3xHA-bGHpA;U6::BsaI-sgRNA and PB-INS-CAG-spCas9-nlsTagBFP2, respectively, using AA052/AA053 and AA054/AA032, respectively, and the Phusion PCR protocol described in **Supplementary Note 1**, with an annealing temperature of 65 °C and an extension time of 120 s. The amplified fragments were size-selected by electrophoresis on a 1% agarose gel and gel-purified using the GeneJET Gel Extraction Kit according to the manufacturer’s protocol. The gel-purified SaCas9 and mTagBFP2 fragments were then assembled into the restriction-enzyme-digested PB-INS-CAG-US1-eGFP-EF1a-Neo backbone by Gibson assembly at 50 °C for 1 h at a 2:2:1 molar ratio (insert:insert:backbone). The assembled product was confirmed by whole-plasmid sequencing.

##### 15.5.5. piggyBac vectors expressing AGO2 or mCherry control and mTagBFP2

To produce PB-CAG-hAgo2-EF1-mTagBFP2 and PB-CAG-control-EF1-mTagBFP2, the PB-CAG-US1-eGFP-EF1-Target-AID backbone was first digested with AvrII-HF and ClaI for 2 h at 37 °C using the common restriction enzyme digestion protocol described in **Supplementary Note 1**. The backbone band was size-selected by electrophoresis on a 1% agarose gel and gel-purified using the GeneJET Gel Extraction Kit according to the manufacturer’s protocol. The mTagBFP2 fragment was amplified from dAAVS1_EF1a-BxB1_mTagBFP2 using AA055/AA056 and the Phusion PCR protocol described in **Supplementary Note 1**, with an annealing temperature of 62 °C and an extension time of 30 s. The amplified fragment was size-selected by electrophoresis on a 1% agarose gel and gel-purified using the GeneJET Gel Extraction Kit according to the manufacturer’s protocol. The gel-purified mTagBFP2 PCR product was then digested with AvrII-HF and ClaI for 1 h at 37 °C using the common restriction enzyme digestion protocol described in **Supplementary Note 1**, PCR-purified using the GeneJET Gel Extraction Kit according to the manufacturer’s protocol, and ligated between the AvrII and ClaI sites of the digested PB-CAG-US1-eGFP-EF1-Target-AID backbone at a 3:1 insert-to-backbone molar ratio using T4 DNA Ligase in 1× T4 DNA Ligase Buffer with ATP at 25 °C for 10 min. The resulting intermediate vector was confirmed by Sanger sequencing for correct insertion and then digested with NotI-HF and AgeI-HF for 2 h at 37 °C using the common restriction enzyme digestion protocol described in **Supplementary Note 1**. The backbone band was size-selected by electrophoresis on a 1% agarose gel and gel-purified using the GeneJET Gel Extraction Kit according to the manufacturer’s protocol.

The N-terminal and C-terminal halves of AGO2 were amplified with homologous overlapping regions from two gene blocks synthesized by Twist Bioscience (GID006 and GID007) using AA057/AA058 and AA059/AA060, respectively, and the Phusion PCR protocol described in **Supplementary Note 1**, with an annealing temperature of 65 °C and an extension time of 90 s. The amplified fragments were size-selected by electrophoresis on a 1% agarose gel and gel-purified using the GeneJET Gel Extraction Kit according to the manufacturer’s protocol. The gel-purified AGO2 fragments were then assembled into the restriction-enzyme-digested vector from the previous step by Gibson assembly at 50 °C for 1 h at a 3:1 insert-to-vector molar ratio.

For construction of the mCherry control vector, the mCherry fragment was amplified from pSI-173-CMVp-MCS-mCherry using AA020/AA021 and the Phusion PCR protocol described in **Supplementary Note 1**, with an annealing temperature of 62 °C and an extension time of 30 s. The amplified fragment was size-selected by electrophoresis on a 1% agarose gel and gel-purified using the GeneJET Gel Extraction Kit according to the manufacturer’s protocol. The gel-purified mCherry PCR product was then digested with NotI-HF and AgeI-HF for 1 h at 37 °C using the common restriction enzyme digestion protocol described in **Supplementary Note 1**, PCR-purified using the GeneJET Gel Extraction Kit according to the manufacturer’s protocol, and ligated between the NotI and AgeI sites of the same digested vector used for the AGO2 Gibson assembly at a 3:1 insert-to-backbone molar ratio using T4 DNA Ligase in 1× T4 DNA Ligase Buffer with ATP at 25 °C for 10 min. The assembled products were confirmed by whole-plasmid sequencing.

##### 15.5.6. piggyBac vectors expressing hAgo2, Luc2 or hAgo2ΔPIWI and mCherry

To produce PB-INS-CAG-hAgo2-EF1-mCherry, PB-INS-CAG-Luc2-EF1-mCherry and PB-INS-CAG-hAgo2ΔPIWI-EF1-mCherry, the PB-INS-CAG-US1-eGFP-EF1a-Neo backbone was first digested with AvrII-HF and SalI-HF for 2 h at 37 °C using the common restriction enzyme digestion protocol described in **Supplementary Note 1**. The backbone band was size-selected by electrophoresis on a 1% agarose gel and gel-purified using the GeneJET Gel Extraction Kit according to the manufacturer’s protocol. The mCherry fragment was amplified from PB-CAG-control-EF1-mTagBFP2 using AA061/AA062 and the Phusion PCR protocol described in **Supplementary Note 1**, with an annealing temperature of 62 °C and an extension time of 30 s. The amplified fragment was size-selected by electrophoresis on a 1% agarose gel and gel-purified using the GeneJET Gel Extraction Kit according to the manufacturer’s protocol. The gel-purified mCherry PCR product was then digested with AvrII-HF and SalI for 1 h at 37 °C using the common restriction enzyme digestion protocol described in **Supplementary Note 1**, PCR-purified using the GeneJET Gel Extraction Kit according to the manufacturer’s protocol, and ligated between the AvrII and SalI sites of the digested PB-INS-CAG-US1-eGFP-EF1a-Neo backbone at a 3:1 insert-to-backbone molar ratio using T4 DNA Ligase in 1× T4 DNA Ligase Buffer with ATP at 25 °C for 10 min.

The resulting intermediate vector was validated for correct insertion by restriction-enzyme digestion with AvrII-HF and SalI-HF for 1 h at 37 °C using the common restriction enzyme digestion protocol described in **Supplementary Note 1**. The validated vector was then digested with NotI-HF and AgeI-HF for 2 h at 37 °C using the same common restriction enzyme digestion protocol. The backbone band was size-selected by electrophoresis on a 1% agarose gel and gel-purified using the GeneJET Gel Extraction Kit according to the manufacturer’s protocol. The AGO2 and AGO2ΔPIWI fragments were amplified from PB-CAG-hAgo2-EF1-mTagBFP2 using AA057/AA060 and AA057/AA063, respectively, and the Luc2 fragment was amplified from pHAGE-CMV-Luc2-IRES-ZsGreen-W using AA064/AA065. The fragments were amplified using the Phusion PCR protocol described in **Supplementary Note 1**, with an annealing temperature of 62 °C and an extension time of 90 s. The amplified fragments were size-selected by electrophoresis on a 1% agarose gel and gel-purified using the GeneJET Gel Extraction Kit according to the manufacturer’s protocol. The gel-purified AGO2, Luc2 and AGO2ΔPIWI PCR products were then assembled into the restriction-enzyme-digested vector from the previous step by Gibson assembly at 50 °C for 1 h at a 3:1 vector-to-insert molar ratio. The assembled products were confirmed by whole-plasmid sequencing.

##### 15.5.7. piggyBac vectors expressing codon-optimized wild-type SpCas9 with mTagBFP2 and Nanog- or Acta2-targeting sgRNAs

To produce PB-INS-CAG-spCas9-nlsTagBFP2-NANOG-sgRNA and PB-INS-CAG-spCas9-nlsTagBFP2-ACTA2-sgRNA, PB-INS-CAG-spCas9-nlsTagBFP2 was first digested with HindIII-HF (NEB, no. R3104L) and HpaI-HF (NEB, no. R0105L) for 2 h at 37 °C using the common restriction enzyme digestion protocol described in **Supplementary Note 1**. The backbone band was size-selected by electrophoresis on a 1% agarose gel and gel-purified using the GeneJET Gel Extraction Kit according to the manufacturer’s protocol.

The *Nanog*-sgRNA and *Acta2*-sgRNA fragments were generated by first cloning *Nanog*- and *Acta2*-targeting spacers, respectively, into PB-sgRNA-BB using the Golden Gate assembly protocol described in **Supplementary Note 3** and the corresponding DNA oligonucleotide pairs listed in **Table S6**. The resulting sgRNA constructs were confirmed by Sanger sequencing for correct insertion and used in subsequent cloning steps. The bGH poly(A)-hU6-*Nanog*-sgRNA and bGH poly(A)-hU6-*Acta2*-sgRNA fragments were then amplified from the PB-Nanog-sgRNA and PB-Acta2-sgRNA cloning products, respectively, using AA066/AA067 and the Phusion PCR protocol described in **Supplementary Note 1**, with an annealing temperature of 65 °C and an extension time of 30 s. The amplified fragments were size-selected by electrophoresis on a 1% agarose gel and gel-purified using the GeneJET Gel Extraction Kit according to the manufacturer’s protocol.

The gel-purified bGH poly(A)-hU6-*Nanog*-sgRNA and bGH poly(A)-hU6-*Acta2*-sgRNA PCR products were then digested with HindIII-HF and HpaI-HF for 1 h at 37 °C using the common restriction enzyme digestion protocol described in **Supplementary Note 1**, PCR-purified using the GeneJET Gel Extraction Kit according to the manufacturer’s protocol, and ligated between the HindIII and HpaI sites of the digested PB-INS-CAG-spCas9-nlsTagBFP2 vector at a 3:1 insert-to-backbone molar ratio using T4 DNA Ligase in 1× T4 DNA Ligase Buffer with ATP at 16 °C for 1 h. The assembled products were confirmed by whole-plasmid sequencing.

#### 15.6. Lentiviral expression vectors

##### 15.6.1. Lentiviral vectors expressing mCherry and cgRNA V3, V4, cgRNA V3-SL2 (cgRNA V5) or US1-sgRNA

To produce LVSIN-U6-cgRNA-V3-EF1a-mCherry (cloning backbone), LVSIN-U6-US1-sgRNA-EF1a-mCherry, LVSIN-U6-cgRNA-V3-SL2-EF1a-mCherry (cloning backbone) and LVSIN-U6-cgRNA-V4-EF1a-mCherry (cloning backbone), pLV-SI-112 was first digested with BamHI-HF and MluI-HF for 2 h at 37 °C using the common restriction enzyme digestion protocol described in **Supplementary Note 1**. The backbone band was size-selected by electrophoresis on a 1% agarose gel and gel-purified using the GeneJET Gel Extraction Kit according to the manufacturer’s protocol. The EF1a promoter region and mCherry coding sequence were amplified with homologous overlapping regions from PB-CAG-US1-eGFP-EF1-Target-AID and pSI-173-CMVp-MCS-mCherry, respectively, using AA068/AA069 and AA070/AA071, respectively, and the Phusion PCR protocol described in **Supplementary Note 1**, with an annealing temperature of 63 °C and an extension time of 60 s. The amplified fragments were size-selected by electrophoresis on a 1% agarose gel and gel-purified using the GeneJET Gel Extraction Kit according to the manufacturer’s protocol. The gel-purified EF1a promoter and mCherry coding-sequence fragments were then assembled into the restriction-enzyme-digested pLV-SI-112 backbone by Gibson assembly at 50 °C for 1 h at a 3:3:1 molar ratio (insert:insert:vector). The assembled product was confirmed by Sanger sequencing and then digested with BamHI-HF and ClaI for 2 h at 37 °C using the common restriction enzyme digestion protocol described in **Supplementary Note 1**. The backbone band was size-selected by electrophoresis on a 1% agarose gel and gel-purified using the GeneJET Gel Extraction Kit according to the manufacturer’s protocol. The hU6-cgRNA-V3, hU6-US1-sgRNA and hU6-cgRNA-V3-SL2 fragments were amplified from PB4-cgRNA-V3, PB4-US1-sgRNA and a gene block synthesized by Twist Bioscience (GID008), respectively, using AA072/AA073, AA072/AA074 and AA072/AA075, respectively, and the Phusion PCR protocol described in **Supplementary Note 1**, with an annealing temperature of 61 °C and an extension time of 30 s. The amplified fragments were size-selected by electrophoresis on a 1% agarose gel and gel-purified using the GeneJET Gel Extraction Kit according to the manufacturer’s protocol. hU6-cgRNA-V4 was synthesized by Twist Bioscience as a gene block (GID009) flanked by BamHI and ClaI restriction sites. The gel-purified hU6-cgRNA-V3, hU6-US1-sgRNA and hU6-cgRNA-V3-SL2 PCR products and the hU6-cgRNA-V4 gene block were then digested with BamHI-HF and ClaI for 1 h at 37 °C using the common restriction enzyme digestion protocol described in **Supplementary Note 1**, PCR-purified using the GeneJET Gel Extraction Kit according to the manufacturer’s protocol, and ligated between the BamHI and ClaI sites of the digested first-assembly vector at a 3:1 insert-to-backbone molar ratio using T4 DNA Ligase in 1× T4 DNA Ligase Buffer with ATP at 25 °C for 10 min.

To produce pLVSIN-U6-RFP-US1-cgRNA-V3-EF1a-mCherry (cloning backbone), LVSIN-U6-cgRNA-V3-EF1a-mCherry was first digested with BamHI-HF and ClaI for 2 h at 37 °C using the common restriction enzyme digestion protocol described in **Supplementary Note 1**. The backbone band was size-selected by electrophoresis on a 1% agarose gel and gel-purified using the GeneJET Gel Extraction Kit according to the manufacturer’s protocol. The hU6 and US1-cgRNA-V3 fragments were amplified from LVSIN-U6-122T-US1-cgRNA-V3-EF1a-mCherry using AA072/AA093 and AA073/AA094, respectively, and the Phusion PCR protocol described in **Supplementary Note 1**, with an annealing temperature of 69 °C and an extension time of 30 s. The amplified fragments were size-selected by electrophoresis on a 1% agarose gel and gel-purified using the GeneJET Gel Extraction Kit according to the manufacturer’s protocol. The purified hU6 and US1-cgRNA-V3 fragments were mixed at equal volumes (1 µl each), and the mixture was amplified using AA072/AA073 and the Phusion PCR protocol described in **Supplementary Note 1**, with an annealing temperature of 69 °C and an extension time of 30 s. The resulting hU6-US1-cgRNA-V3 fusion PCR product was size-selected by electrophoresis on a 1% agarose gel and gel-purified using the GeneJET Gel Extraction Kit according to the manufacturer’s protocol. The gel-purified hU6-US1-cgRNA-V3 product was then digested with BamHI-HF and ClaI for 1 h at 37 °C using the common restriction enzyme digestion protocol described in **Supplementary Note 1**, PCR-purified using the GeneJET Gel Extraction Kit according to the manufacturer’s protocol, and ligated between the BamHI and ClaI sites of the digested first-assembly vector at a 3:1 insert-to-backbone molar ratio using T4 DNA Ligase in 1× T4 DNA Ligase Buffer with ATP at 25 °C for 10 min.

The assembled product was confirmed by Sanger sequencing, after which 15 µl of the assembled vector was digested with BsmBI-v2 (NEB, no. R0739L) in 1× NEB 3.1 Buffer in a total reaction volume of 30 µl for 2 h at 55 °C. The backbone band was size-selected by electrophoresis on a 1% agarose gel and gel-purified using the GeneJET Gel Extraction Kit according to the manufacturer’s protocol. The RFP transcriptional-unit fragment was amplified from pU6-pegRNA-GG-acceptor using AA095/AA096 and the Phusion PCR protocol described in **Supplementary Note 1**, with an annealing temperature of 69 °C and an extension time of 30 s. The amplified fragment was size-selected by electrophoresis on a 1% agarose gel and gel-purified using the GeneJET Gel Extraction Kit according to the manufacturer’s protocol. The gel-purified RFP PCR product was then digested with 1 µl of BsaI-HF-v2 (NEB, no. R3733L) for 1 h at 37 °C under the common restriction enzyme digestion conditions described in **Supplementary Note 1**, PCR-purified using the GeneJET Gel Extraction Kit according to the manufacturer’s protocol, and ligated between the complementary overhangs generated by BsmBI-v2 digestion of the first-assembly vector at a 3:1 insert-to-backbone molar ratio using T4 DNA Ligase in 1× T4 DNA Ligase Buffer with ATP at 25 °C for 10 min.

##### 15.6.2. Lentiviral vector expressing mCherry and synthetic miRNAs

To produce LVSIN-U6-pri-miRNA-V3-EF1a-mCherry (cloning backbone), LVSIN-U6-cgRNA-V3-EF1a-mCherry was first digested with BamHI-HF and ClaI-HF for 2 h at 37 °C using the common restriction enzyme digestion protocol described in **Supplementary Note 1**. The backbone band was size-selected by electrophoresis on a 1% agarose gel and gel-purified using the GeneJET Gel Extraction Kit according to the manufacturer’s protocol. The hU6-pri-miRNA-scaffold was synthesized by Twist Bioscience as a gene block (GID010), digested with BamHI-HF and ClaI for 1 h at 37 °C using the common restriction enzyme digestion protocol described in **Supplementary Note 1**, PCR-purified using the GeneJET Gel Extraction Kit according to the manufacturer’s protocol, and ligated between the BamHI and ClaI sites of the digested LVSIN-U6-cgRNA-V3-EF1a-mCherry backbone at a 3:1 insert-to-backbone molar ratio using T4 DNA Ligase in 1× T4 DNA Ligase Buffer with ATP at 25 °C for 10 min.

##### 15.6.3. Lentiviral vector expressing mCherry–IRES–EGFP for assessing wild-type Cas9 activity

To produce LVSIN-EF1-mCherry-IRES-EGFP, LVSIN-U6-cgRNA-V3-EF1a-mCherry was first digested with MluI-HF and ClaI for 2 h at 37 °C using the common restriction enzyme digestion protocol described in **Supplementary Note 1**. The backbone band was size-selected by electrophoresis on a 1% agarose gel and gel-purified using the GeneJET Gel Extraction Kit according to the manufacturer’s protocol. The EF1-mCherry and IRES-EGFP fragments were amplified from double-stranded DNA gene blocks purchased from Twist Bioscience (GID011 and GID012) using AA076/AA077 and AA078/AA079, respectively, and the Phusion PCR protocol described in **Supplementary Note 1**, with an annealing temperature of 65 °C and an extension time of 90 s. The amplified fragments were size-selected by electrophoresis on a 1% agarose gel and gel-purified using the GeneJET Gel Extraction Kit according to the manufacturer’s protocol. The purified EF1-mCherry and IRES-EGFP fragments and the digested and purified LVSIN-U6-cgRNA-V3-EF1a-mCherry backbone were then mixed at a 3:3:1 molar ratio (insert:insert:backbone) and assembled by Gibson assembly at 50 °C for 1 h.

##### 15.6.4. Lentiviral vectors expressing mCherry and sgRNA, crRNA:tracrRNA, miR-122-targeted MICR and cgRNA transcriptional units with a bGH poly(A) signal for comparison with MICR

To produce LVSIN-pA30-bgH-EF1a-mCherry, LVSIN-U6-cgRNA-V3-EF1a-mCherry was first digested with BamHI-HF and ClaI for 2 h at 37 °C using the common restriction enzyme digestion protocol described in **Supplementary Note 1**. The backbone band was size-selected by electrophoresis on a 1% agarose gel and gel-purified using the GeneJET Gel Extraction Kit according to the manufacturer’s protocol. The TETp-RFP transcriptional unit and bGH poly(A) signal sequence were amplified with homologous overlapping regions from pU6-pegRNA-GG-acceptor (Addgene, no. 132777) and PB4-CAGGS-17T-sgRNA-MICR, respectively, using AA080/AA081 and AA082/AA083, respectively, and the Phusion PCR protocol described in **Supplementary Note 1**, with an annealing temperature of 60 °C and an extension time of 60 s. The amplified fragments were size-selected by electrophoresis on a 1% agarose gel and gel-purified using the GeneJET Gel Extraction Kit according to the manufacturer’s protocol. The gel-purified TETp-RFP transcriptional unit and bGH poly(A) signal sequence were then assembled into the restriction-enzyme-digested LVSIN-pA30-bgH-EF1a-mCherry backbone by Gibson assembly at 50 °C for 1 h at a 3:5:1 molar ratio (insert:insert:backbone).

The assembled product was confirmed by Sanger sequencing and then digested with HpaI-HF and NotI-HF for 2 h at 37 °C using the common restriction enzyme digestion protocol described in **Supplementary Note 1**. The backbone band was size-selected by electrophoresis on a 1% agarose gel and gel-purified using the GeneJET Gel Extraction Kit according to the manufacturer’s protocol. PB4-US1-sgRNA, PB4-Scr-sgRNA, PB4-122T-US1-MICR, PB4-122T-Scr-MICR, PB4-122T-US1-cgRNA-V1, PB4-122T-US1-cgRNA-V2, PB4-122T-US1-cgRNA-V3, PB4-122T-Scr-cgRNA-V1, PB4-122T-Scr-cgRNA-V2 and PB4-122T-Scr-cgRNA-V3 were then digested with EcoRV-HF (NEB, no. R3195L) and NotI-HF for 2 h at 37 °C using the common restriction enzyme digestion protocol described in **Supplementary Note 1**. The gRNA-containing fragments were size-selected by electrophoresis on a 1% agarose gel and gel-purified using the GeneJET Gel Extraction Kit according to the manufacturer’s protocol. The gel-purified gRNA-containing fragments were then ligated into the digested LVSIN-pA30-bgH-EF1a-mCherry backbone at a 3:1 insert-to-backbone molar ratio using T4 DNA Ligase in 1× T4 DNA Ligase Buffer with ATP at 16 °C for 1 h.

To produce the US1-crRNA:tracrRNA and Scr-crRNA:tracrRNA constructs, the hU6-US1-crRNA-mU6-TRACR and hU6-Scr-crRNA-mU6-TRACR fragments were synthesized by Twist Bioscience as double-stranded DNA gene blocks (GID013 and GID014), digested with NotI-HF (NEB) for 2 h at 37 °C under the common restriction enzyme digestion conditions described in **Supplementary Note 1**, PCR-purified using the GeneJET Gel Extraction Kit according to the manufacturer’s protocol, and ligated into the digested LVSIN-pA30-bgH-EF1a-mCherry backbone at a 3:1 insert-to-backbone molar ratio using T4 DNA Ligase in 1× T4 DNA Ligase Buffer with ATP at 16 °C for 1 h.

##### 15.6.5. Lentiviral acceptor vectors for cgRNA transcriptional units with puromycin resistance

To produce LVSIN-acceptor-pGK-Puro, pLVSIN-CMV Pur Vector (Takara Bio) was first digested with EcoRI-HF and ClaI for 2 h at 37 °C using the common restriction enzyme digestion protocol described in **Supplementary Note 1**. The backbone band was size-selected by electrophoresis on a 1% agarose gel and gel-purified using the GeneJET Gel Extraction Kit according to the manufacturer’s protocol. The TETp-RFP transcriptional unit was amplified from pU6-pegRNA-GG-acceptor using AA084/AA085 and the Phusion PCR protocol described in **Supplementary Note 1**, with an annealing temperature of 65 °C and an extension time of 30 s. The amplified fragment was size-selected by electrophoresis on a 1% agarose gel and gel-purified using the GeneJET Gel Extraction Kit according to the manufacturer’s protocol. The gel-purified TETp-RFP PCR product was then digested with EcoRI-HF and ClaI for 1 h at 37 °C using the common restriction enzyme digestion protocol described in **Supplementary Note 1**, PCR-purified using the GeneJET Gel Extraction Kit according to the manufacturer’s protocol, and ligated between the EcoRI and ClaI sites of the digested pLVSIN-CMV Pur Vector at a 3:1 insert-to-backbone molar ratio using T4 DNA Ligase in 1× T4 DNA Ligase Buffer with ATP at 25 °C for 10 min. The assembled product was confirmed by whole-plasmid sequencing.

To produce LVcHS4-acceptor-pGK-Puro, pLV.CMVenh.gp91.eGFP.cHS4 was first digested with XhoI (NEB, no. R0146L) and BsrGI-HF (NEB, no. R3575L) for 2 h at 37 °C using the common restriction enzyme digestion protocol described in **Supplementary Note 1**. The backbone band was size-selected by electrophoresis on a 1% agarose gel and gel-purified using the GeneJET Gel Extraction Kit according to the manufacturer’s protocol. The pTET-RFP-pGK-Puro transcriptional unit was amplified from LVSIN-acceptor-pGK-Puro using AA084/AA086 and the Phusion PCR protocol described in **Supplementary Note 1**, with an annealing temperature of 62 °C and an extension time of 90 s. The amplified fragment was size-selected by electrophoresis on a 1% agarose gel and gel-purified using the GeneJET Gel Extraction Kit according to the manufacturer’s protocol. The gel-purified pTET-RFP-pGK-Puro PCR product was then digested with XhoI-HF and BsrGI-HF for 1 h at 37 °C using the common restriction enzyme digestion protocol described in **Supplementary Note 1**, PCR-purified using the GeneJET Gel Extraction Kit according to the manufacturer’s protocol, and ligated between the XhoI and BsrGI sites of the digested pLV.CMVenh.gp91.eGFP.cHS4 backbone at a 3:1 insert-to-backbone molar ratio using T4 DNA Ligase in 1× T4 DNA Ligase Buffer with ATP at 25 °C for 10 min. The assembled product was confirmed by whole-plasmid sequencing.

##### 15.6.6. Lentiviral vectors expressing puromycin resistance and cgRNA V3 or US1-sgRNA

To produce LVSIN-U6-US1-sgRNA-pGK-Puro, LVSIN-U6-21T-US1-cgRNA-V3-pGK-Puro and LVSIN-U6-NCT-US1-cgRNA-V3-pGK-Puro, LVSIN-acceptor-pGK-Puro was first digested with BamHI-HF and ClaI for 2 h at 37 °C using the common restriction enzyme digestion protocol described in **Supplementary Note 1**. The backbone band was size-selected by electrophoresis on a 1% agarose gel and gel-purified using the GeneJET Gel Extraction Kit according to the manufacturer’s protocol. The hU6-21T-US1-cgRNA-V3, hU6-NCT-US1-cgRNA-V3 and hU6-US1-sgRNA fragments were amplified from LVSIN-U6-21T-US1-cgRNA-V3-EF1a-mCherry, LVSIN-U6-NCT-US1-cgRNA-V3-EF1a-mCherry and LVSIN-U6-US1-sgRNA-EF1a-mCherry, respectively, using AA072/AA073, AA072/AA073 and AA072/AA074,respectively, and the Phusion PCR protocol described in **Supplementary Note 1**, with an annealing temperature of 62 °C and an extension time of 30 s. The amplified fragments were size-selected by electrophoresis on a 1% agarose gel and gel-purified using the GeneJET Gel Extraction Kit according to the manufacturer’s protocol. The gel-purified cgRNA-V3 and sgRNA PCR products were then digested with BamHI-HF and ClaI for 1 h at 37 °C using the common restriction enzyme digestion protocol described in **Supplementary Note 1**, PCR-purified using the GeneJET Gel Extraction Kit according to the manufacturer’s protocol, and ligated between the BamHI and ClaI sites of the digested LVSIN-acceptor-pGK-Puro backbone at a 3:1 insert-to-backbone molar ratio using T4 DNA Ligase in 1× T4 DNA Ligase Buffer with ATP at 25 °C for 10 min.

To produce LVcHS4-RFP-cgRNA-V3-pGK-Puro (cloning backbone) and LVcHS4-RFP-US1-cgRNA-V3-pGK-Puro (cloning backbone), LVcHS4-acceptor-pGK-Puro was first digested with 1 µl of BamHI-HF for 2 h at 37 °C and then PCR-purified using the GeneJET Gel Extraction Kit according to the manufacturer’s protocol, with an elution volume of 26 µl. The purified digestion product was subsequently digested with 1 µl of FastDigest KflI (Thermo Fisher Scientific, no. FD2164) in 1× FastDigest Buffer for 2 h at 37 °C. The backbone band was size-selected by electrophoresis on a 1% agarose gel and gel-purified using the GeneJET Gel Extraction Kit according to the manufacturer’s protocol. For the assembly of LVcHS4-RFP-cgRNA-V3-pGK-Puro (cloning backbone), the 5′-hU6-cgRNA-V3, TETp-RFP and 3′-cgRNA-V3 fragments were amplified with homologous overlapping regions from LVSIN-U6-cgRNA-V3-EF1a-mCherry, pU6-pegRNA-GG-acceptor and LVSIN-U6-cgRNA-V3-EF1a-mCherry, respectively, using AA087/AA088, AA089/AA090 and AA091/AA092, respectively, and the Phusion PCR protocol described in **Supplementary Note 1**, with an annealing temperature of 60 °C and an extension time of 30 s. LVcHS4-RFP-US1-cgRNA-V3-pGK-Puro (cloning backbone), the 5′-hU6-cgRNA-V3, TETp-RFP-US1 and 3′-US1-cgRNA-V3 fragments were amplified with homologous overlapping regions from LVSIN-U6-cgRNA-V3-EF1a-mCherry, pU6-pegRNA-GG-acceptor and LVSIN-U6-122T-US1-cgRNA-V3-EF1a-mCherry, respectively, using AA087/AA088, AA089/AA128 and AA129/AA092, respectively, and the Phusion PCR protocol described in **Supplementary Note 1**, with an annealing temperature of 60 °C and an extension time of 30 s.

The amplified fragments were size-selected by electrophoresis on a 1% agarose gel and gel-purified using the GeneJET Gel Extraction Kit according to the manufacturer’s protocol. The gel-purified 5′-hU6-cgRNA-V3, TETp-RFP and 3′-cgRNA-V3 fragments or the 5′-hU6-cgRNA-V3, TETp-RFP-US1 and 3′-US1-cgRNA-V3 fragments were then assembled into the restriction-enzyme-digested LVcHS4-acceptor-pGK-Puro backbone by Gibson assembly at 50 °C for 1 h at a 3:3:3:1 molar ratio (insert:insert:insert:backbone).

LVcHS4-US1-sgRNA-pGK-Puro was produced exactly as described for the construction of LVSIN-U6-US1-sgRNA-pGK-Puro but the LVcHS4-RFP-cgRNA-V3-pGK-Puro (cloning backbone) was used in place of LVSIN-acceptor-pGK-Puro as the recipient backbone.

To produce LVcHS4-cgRNA-saV3-pGK-Puro (cloning backbone), LVcHS4-cgRNA-V3-pGK-Puro was first digested with BamHI-HF and ClaI for 2 h at 37 °C using the common restriction enzyme digestion protocol described in **Supplementary Note 1**. The backbone band was size-selected by electrophoresis on a 1% agarose gel and gel-purified using the GeneJET Gel Extraction Kit according to the manufacturer’s protocol. The SaCas9-cgRNA-V3 fragment was synthesized by Twist Bioscience as a double-stranded DNA gene block (GID015), digested with BamHI-HF and ClaI (NEB) for 2 h at 37 °C using the common restriction enzyme digestion protocol described in **Supplementary Note 1**, PCR-purified using the GeneJET Gel Extraction Kit according to the manufacturer’s protocol, and ligated between the BamHI and ClaI sites of the digested LVcHS4-cgRNA-V3-pGK-Puro backbone at a 3:1 insert-to-backbone molar ratio using T4 DNA Ligase in 1× T4 DNA Ligase Buffer with ATP at 25 °C for 10 min.

##### 15.6.7. Lentiviral vectors expressing puromycin resistance and cgRNA V3 sensors with extended linkers

To produce pAA072–pAA081, LVSIN-acceptor-pGK-Puro was first digested with BamHI-HF and ClaI for 2 h at 37 °C using the common restriction enzyme digestion protocol described in **Supplementary Note 1**. The backbone band was size-selected by electrophoresis on a 1% agarose gel and gel-purified using the GeneJET Gel Extraction Kit according to the manufacturer’s protocol. The hU6-cgRNA-V3 transcriptional units were amplified from pAA072–pAA081 using AA072/AA073 and the Phusion PCR protocol described in **Supplementary Note 1**, with an annealing temperature of 62 °C and an extension time of 30 s. The amplified fragments were size-selected by electrophoresis on a 1% agarose gel and gel-purified using the GeneJET Gel Extraction Kit according to the manufacturer’s protocol. The gel-purified cgRNA-V3 PCR products were then digested with BamHI-HF and ClaI for 1 h at 37 °C using the common restriction enzyme digestion protocol described in **Supplementary Note 1**, PCR-purified using the GeneJET Gel Extraction Kit according to the manufacturer’s protocol, and ligated between the BamHI and ClaI sites of the digested LVSIN-acceptor-pGK-Puro backbone at a 3:1 insert-to-backbone molar ratio using T4 DNA Ligase in 1× T4 DNA Ligase Buffer with ATP at 25 °C for 10 min.

#### 15.7. miR-FF3 gene-trap and HDR donor vectors

##### 15.7.1. piggyBac vectors expressing mCherry and synthetic intronic miRNA or miRNA clusters

To produce PB-INS-CAG-mCherry-synIntron (cloning backbone), the PB-INS-CAG-US1-eGFP-EF1a-Neo backbone was first digested with NotI-HF and HpaI-HF for 2 h at 37 °C using the common restriction enzyme digestion protocol described in **Supplementary Note 1**. The backbone band was size-selected by electrophoresis on a 1% agarose gel and gel-purified using the GeneJET Gel Extraction Kit according to the manufacturer’s protocol. The 5′-mCherry-synIntron and 3′-synIntron fragments were amplified from a gene block synthesized by Twist Bioscience (GID016) using AA097/AA098 and AA099/AA100, respectively, and the Phusion PCR protocol described in **Supplementary Note 1**, with an annealing temperature of 63 °C and an extension time of 30 s. The amplified fragments were size-selected by electrophoresis on a 1% agarose gel and gel-purified using the GeneJET Gel Extraction Kit according to the manufacturer’s protocol. The gel-purified 5′-mCherry-synIntron and 3′-synIntron fragments were mixed at equal volumes (1 µl each), and the mixture was amplified using AA097/AA0100 and the Phusion PCR protocol described in **Supplementary Note 1**, with an annealing temperature of 63 °C and an extension time of 30 s. The resulting mCherry-synIntron fusion PCR product was size-selected by electrophoresis on a 1% agarose gel and gel-purified using the GeneJET Gel Extraction Kit according to the manufacturer’s protocol. The gel-purified mCherry-synIntron product was then digested with NotI-HF and HpaI-HF for 1 h at 37 °C using the common restriction enzyme digestion protocol described in **Supplementary Note 1**, PCR-purified using the GeneJET Gel Extraction Kit according to the manufacturer’s protocol, and ligated between the NotI-HF and HpaI-HF sites of the digested PB-INS-CAG-US1-eGFP-EF1a-Neo backbone at a 3:1 insert-to-backbone molar ratio using T4 DNA Ligase in 1× T4 DNA Ligase Buffer with ATP at 25 °C for 10 min.

A single miR-FF3 sequence was cloned into the PB-INS-CAG-mCherry-synIntron backbone by first amplifying miR-FF3 from LVSIN-U6-pri-miRNA-FF3-EF1a-mCherry using AA101/AA106 and the Phusion PCR protocol described in **Supplementary Note 1**, with an annealing temperature of 60 °C and an extension time of 30 s. The amplified fragment was size-selected by electrophoresis on a 1% agarose gel and gel-purified using the GeneJET Gel Extraction Kit according to the manufacturer’s protocol. The gel-purified miR-FF3 fragment was then assembled into the PB-INS-CAG-mCherry-synIntron-filler backbone in a 25-µl Golden Gate assembly reaction at a 3:1 insert-to-vector molar ratio using 50 ng of vector. The reaction contained 1 µl of T4 DNA Ligase, 0.5 µl of PaqCI and 0.25 µl of PaqCI activator (NEB, no. R0745L) in 1× T4 DNA Ligase Buffer with ATP. The reaction was cycled 30 times between 37 °C for 1 min and 16 °C for 1 min, followed by a final incubation at 37 °C for 30 min.

A cluster of three miR-FF3 sequences was assembled into the PB-INS-CAG-mCherry-synIntron-filler backbone by first amplifying miR-FF3 from LVSIN-U6-pri-miRNA-FF3-EF1a-mCherry using AA101/AA102, AA103/AA104 and AA105/AA106 and the Phusion PCR protocol described in **Supplementary Note 1**, with an annealing temperature of 60 °C and an extension time of 30 s. The amplified fragments were size-selected by electrophoresis on a 1% agarose gel and gel-purified using the GeneJET Gel Extraction Kit according to the manufacturer’s protocol. The gel-purified miR-FF3 fragments were then assembled into the PB-INS-CAG-mCherry-synIntron-filler backbone at a 3:3:3:1 molar ratio (insert:insert:insert:backbone) using the Golden Gate assembly conditions described above.

##### 15.7.2. piggyBac vectors expressing gene-trap mCherry and a synthetic miRNA cluster

To produce PB-SA-mCherry-synIntron-3xFF3, the PB-INS-CAG-mCherry-cgRNA-V3 backbone was first digested with NheI-HF, BamHI-HF and ClaI-HF for 2 h at 37 °C using the common restriction enzyme digestion conditions described in **Supplementary Note 1**. The backbone band was size-selected by electrophoresis on a 1% agarose gel and gel-purified using the GeneJET Gel Extraction Kit according to the manufacturer’s protocol. The spliceAcceptor-mCherry-synIntron-3xFF3 fragment was amplified from PB-INS-CAG-mCherry-synIntron-3x-FF3 using AA107/AA108 and the Phusion PCR protocol described in **Supplementary Note 1**, with an annealing temperature of 60 °C and an extension time of 60 s. The amplified fragment was size-selected by electrophoresis on a 1% agarose gel and gel-purified using the GeneJET Gel Extraction Kit according to the manufacturer’s protocol. The gel-purified spliceAcceptor-mCherry-synIntron-3xFF3 PCR product was then digested with NheI-HF and ClaI for 1 h at 37 °C using the common restriction enzyme digestion protocol described in **Supplementary Note 1**, PCR-purified using the GeneJET Gel Extraction Kit according to the manufacturer’s protocol, and ligated between the NheI and ClaI sites of the digested PB-INS-CAG-mCherry-cgRNA-V3 backbone at a 3:1 insert-to-backbone molar ratio using T4 DNA Ligase in 1× T4 DNA Ligase Buffer with ATP at 25 °C for 10 min.

##### 15.7.3. HDR templates for knock-in of mCherry and a synthetic miRNA cluster

To produce pNANOG-HDR-donor-BB and pACTA2-HDR-donor-BB, the PB-INS-CAG-mCherry-cgRNA-V3 backbone was first digested with PsiI-v2-HF (NEB, no. R0744L) and MluI-HF for 2 h at 37 °C using the common restriction enzyme digestion protocol described in **Supplementary Note 1**. The backbone band was size-selected by electrophoresis on a 1% agarose gel and gel-purified using the GeneJET Gel Extraction Kit according to the manufacturer’s protocol. The NANOG-HDR-donor-BB and ACTA2-HDR-donor-BB gene blocks synthesized by Twist Bioscience (GID017 and GID018) were digested with 1 µl of MluI-HF for 2 h at 37 °C under the common restriction enzyme digestion conditions described in **Supplementary Note 1**, PCR-purified using the GeneJET Gel Extraction Kit according to the manufacturer’s protocol, and ligated between the PsiI and MluI sites of the digested PB-INS-CAG-mCherry-cgRNA-V3 backbone at a 3:1 insert-to-backbone molar ratio using T4 DNA Ligase in 1× T4 DNA Ligase Buffer with ATP at 16 °C for 1 h.

To produce pNANOG-HDR-mCherry-synIntron-3x-FF3 and pACTA2-HDR-mCherry-synIntron-3x-FF3, pNANOG-HDR-donor-BB and pACTA2-HDR-donor-BB were digested with AgeI-HF and EcoRI-HF for 2 h at 37 °C using the common restriction enzyme digestion protocol described in **Supplementary Note 1**. The backbone bands were size-selected by electrophoresis on a 1% agarose gel and gel-purified using the GeneJET Gel Extraction Kit according to the manufacturer’s protocol. The mCherry-synIntron-3xFF3 fragment was amplified from PB-INS-CAG-mCherry-synIntron-3x-FF3 using AA109/AA100 and the Phusion PCR protocol described in **Supplementary Note 1**, with an annealing temperature of 60 °C and an extension time of 60 s. The amplified fragment was size-selected by electrophoresis on a 1% agarose gel and gel-purified using the GeneJET Gel Extraction Kit according to the manufacturer’s protocol. The gel-purified mCherry-synIntron-3xFF3 PCR product was then digested with AgeI-HF and EcoRI-HF for 1 h at 37 °C using the common restriction enzyme digestion protocol described in **Supplementary Note 1**, PCR-purified using the GeneJET Gel Extraction Kit according to the manufacturer’s protocol, and ligated between the AgeI and EcoRI sites of the digested pNANOG-HDR-mCherry-synIntron-3x-FF3 and pACTA2-HDR-mCherry-synIntron-3x-FF3 vector products at a 3:1 insert-to-backbone molar ratio using T4 DNA Ligase in 1× T4 DNA Ligase Buffer with ATP at 25 °C for 10 min.

#### 15.8. Insulated piggyBac expression vectors for mouse engineering

##### 15.8.1. piggyBac vector expressing a base-editing reporter and mCherry

To produce PB-INS-CAG-US1-eGFP-EF1a-mCherry, PB-INS-CAG-US1-eGFP-EF1a-Neo was first digested with MfeI-HF and SalI-HF (NEB, no. R3138L) for 2 h at 37 °C using the common restriction enzyme digestion protocol described in **Supplementary Note 1**. The backbone band was size-selected by electrophoresis on a 1% agarose gel and gel-purified using the GeneJET Gel Extraction Kit according to the manufacturer’s protocol. The EF1-mCherry transcriptional unit was amplified from LVSIN-U6-cgRNA-V3-EF1a-mCherry (cloning backbone) using AA124/AA062 and the Phusion PCR protocol described in **Supplementary Note 1**, with an annealing temperature of 63 °C and an extension time of 60 s. The amplified fragment was size-selected by electrophoresis on a 1% agarose gel and gel-purified using the GeneJET Gel Extraction Kit according to the manufacturer’s protocol. The gel-purified EF1-mCherry PCR product was then digested with MfeI-HF and SalI-HF for 1 h at 37 °C using the common restriction enzyme digestion protocol described in **Supplementary Note 1**, PCR-purified using the GeneJET Gel Extraction Kit according to the manufacturer’s protocol, and ligated between the MfeI and SalII sites of the digested PB-INS-CAG-US1-eGFP-EF1a-mCherry backbone at a 3:1 insert-to-backbone molar ratio using T4 DNA Ligase in 1× T4 DNA Ligase Buffer with ATP at 25 °C for 10 min.

##### 15.8.2. piggyBac vectors expressing miR-122-targeted cgRNA V3 or US1-sgRNA and Target-AIDmax (three introns)

To produce PB-INS-hU6-miR122-cgRNA-CAG-eAIDmax-BFP and PB-INS-hU6-US1-sgRNA-CAG-eAIDmax-BFP, PB-INS-CAG-TargetAIDmax-nlsTagBFP2 (+3 introns) was first digested with SfiI (NEB, no. R0123L) and SpeI-HF for 2 h at 37 °C using the common restriction enzyme digestion protocol described in **Supplementary Note 1**. The backbone band was size-selected by electrophoresis on a 1% agarose gel and gel-purified using the GeneJET Gel Extraction Kit according to the manufacturer’s protocol.

The hU6-miR122-cgRNA and hU6-US1-sgRNA transcriptional units were amplified from LVSIN-U6-122T-US1-cgRNA-V3-EF1a-mCherry and LVSIN-U6-US1-sgRNA-EF1a-mCherry, respectively, using AA125/AA126 and AA125/AA127, respectively, and the Phusion PCR protocol described in **Supplementary Note 1**, with an annealing temperature of 60 °C and an extension time of 30 s. The amplified fragments were size-selected by electrophoresis on a 1% agarose gel and gel-purified using the GeneJET Gel Extraction Kit according to the manufacturer’s protocol.

The gel-purified hU6-miR122-cgRNA and hU6-US1-sgRNA PCR products were then digested with SfiI and SpeI-HF for 1 h at 37 °C using the common restriction enzyme digestion protocol described in **Supplementary Note 1**, PCR-purified using the GeneJET Gel Extraction Kit according to the manufacturer’s protocol, and ligated between the SfiI and SpeI sites of the digested PB-INS-CAG-TargetAIDmax-nlsTagBFP2 (+3 introns) backbone at a 3:1 insert-to-backbone molar ratio using T4 DNA Ligase in 1× T4 DNA Ligase Buffer with ATP at 25 °C for 10 min.

## Supplementary Note 1. General PCR and restriction enzyme digestion conditions

### Phusion High-Fidelity DNA Polymerase PCR

Unless otherwise specified, PCR amplification was performed using Phusion High-Fidelity DNA Polymerase (NEB, no. M0530L).

#### Reaction setup

PCR reactions were assembled in a total volume of 25 µl as follows:

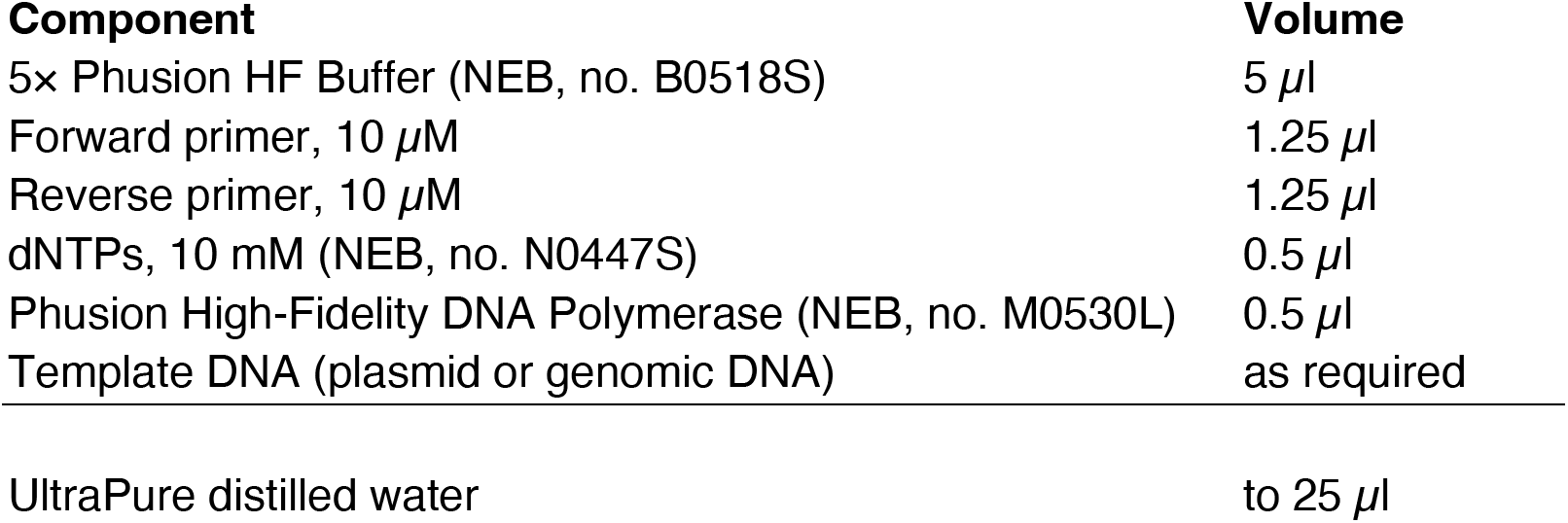

**Thermocycling protocol**

1. Initial denaturation: 98 °C for 30 s.
2. 30 cycles of:

`o` Denaturation: 98 °C for 10 s.
`o` Annealing: primer-specific annealing temperature for 30 s.
`o` Extension: 72 °C for 30 s per kb of expected product.
3. Final extension: 72 °C for 300 s.
4. Hold: 4 °C.

The annealing temperature and extension time were varied according to the primer pair and expected amplicon length, as specified in the corresponding subsection.

### Restriction enzyme digestion

Unless otherwise specified, double restriction enzyme digestions using enzymes from NEB were assembled in a total volume of 30 µl as follows:

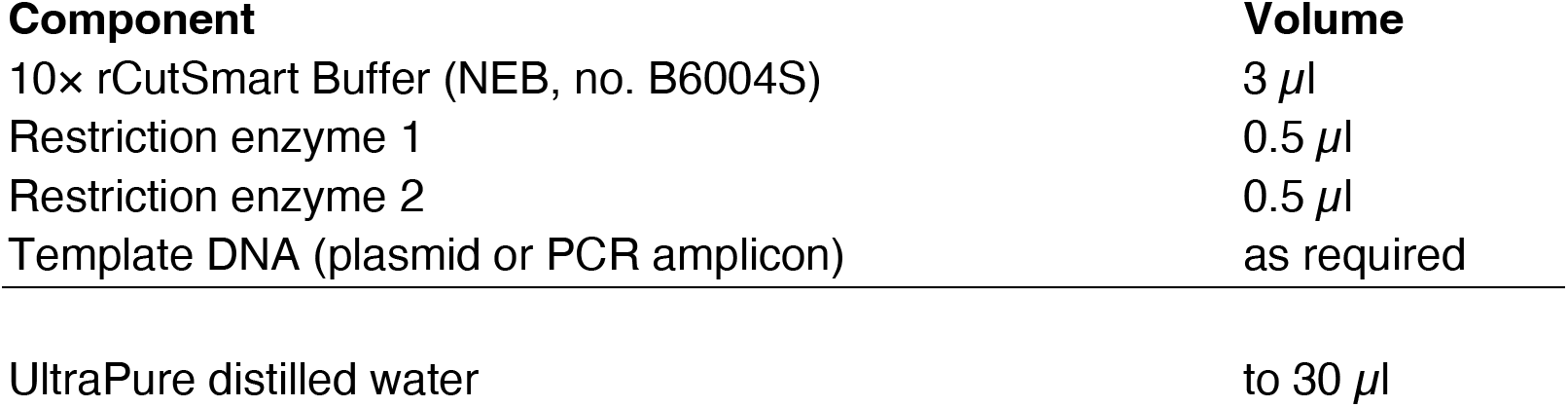

Digestion enzymes, incubation temperatures and incubation times were specified in the corresponding plasmid-construction subsections.

## Supplementary Note 2. Amplicon sequencing library generation

Illumina sequencing libraries of miRNA targets were generated from genomic DNA isolated from cells transduced with the cgRNA library using a two-step PCR protocol.

### PCR round 1: Amplification of miRNA target sequences

#### Reaction setup

PCR round 1 reactions were assembled in a total volume of 50 µl as follows:

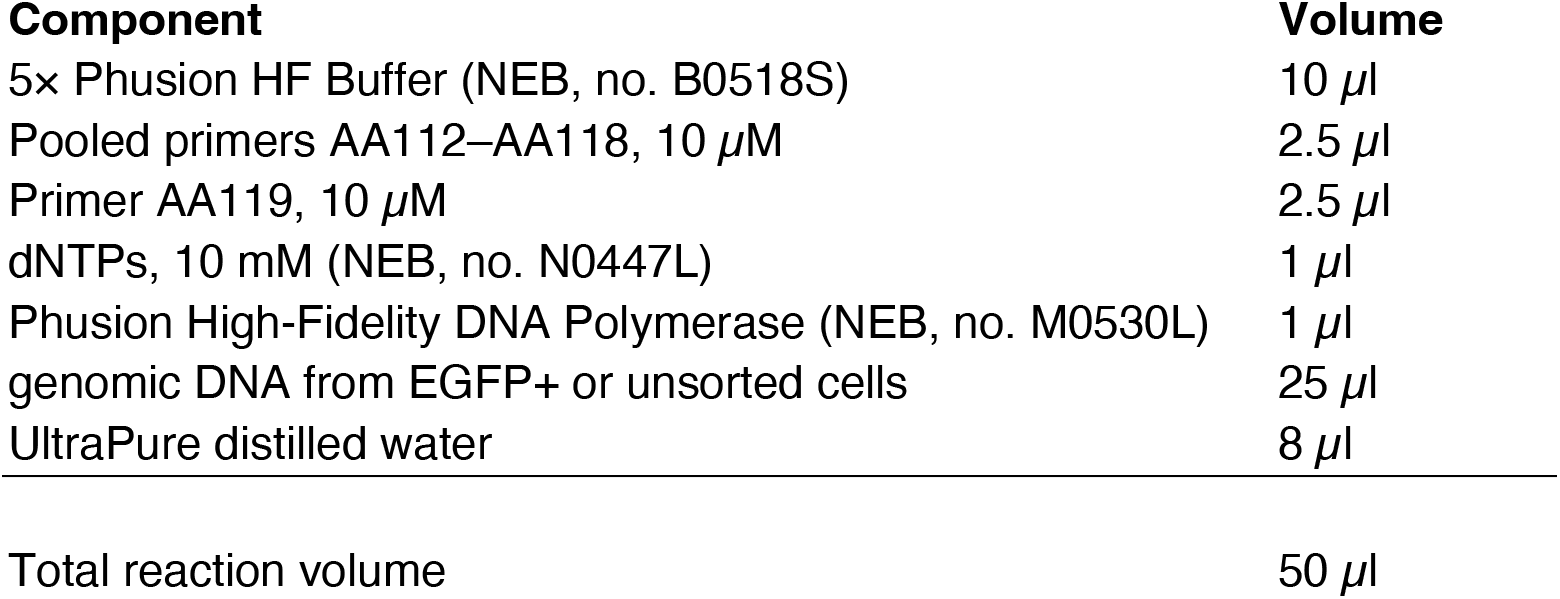

**Thermocycling protocol**

1. Initial denaturation: 98 °C for 30 s.
2. 20 cycles of:

`o` Denaturation: 98 °C for 10 s.
`o` Annealing: 63 °C for 30 s.
`o` Extension: 72 °C for 30 s.
3. Final extension: 72 °C for 300 s.
4. Hold: 4 °C.

The two PCR round 1 reactions for each population were pooled and purified using the GeneJET Gel Extraction Kit according to the manufacturer’s protocol, with a final elution volume of 35 µl in UltraPure distilled water.

### PCR round 2: Addition of Illumina adapters and sample-specific indexes

#### Reaction setup

PCR round 2 reactions were assembled in a total volume of 50 µl as follows:

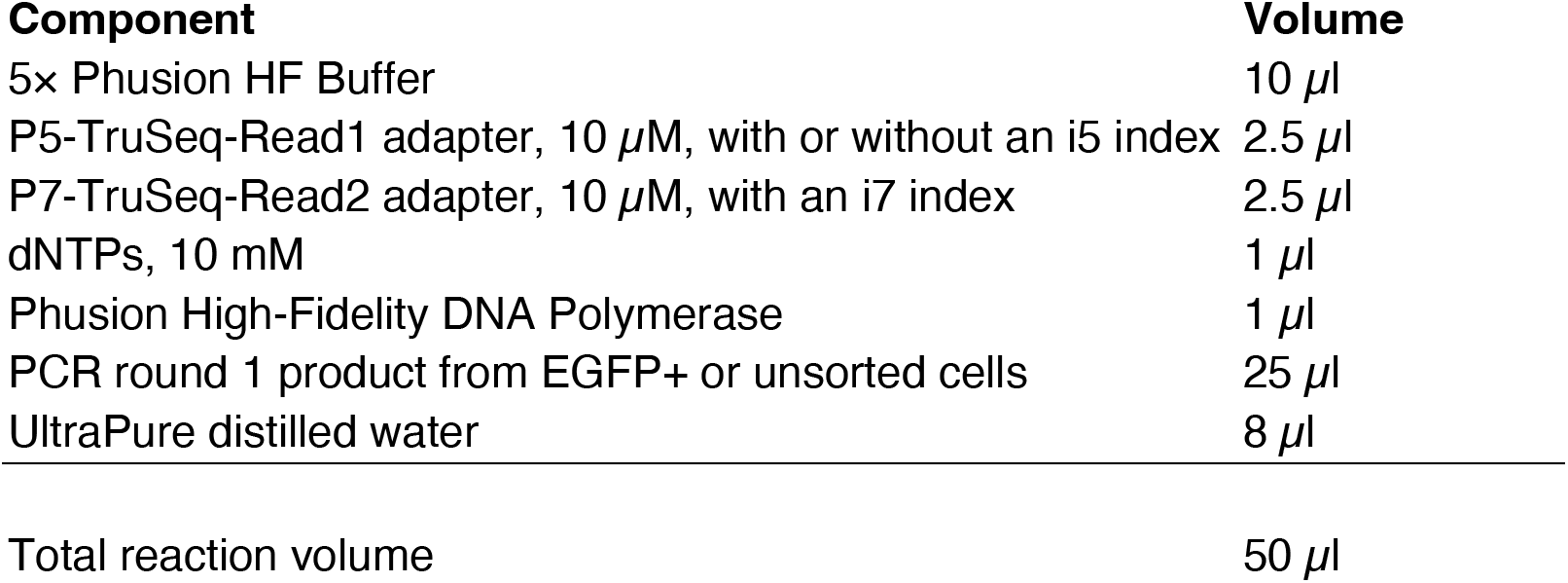

**Thermocycling protocol**

1. Initial denaturation: 98 °C for 30 s.
2. 20 cycles of:

`o` Denaturation: 98 °C for 10 s.
`o` Annealing: 65 °C for 30 s.
`o` Extension: 72 °C for 30 s.
3. Final extension: 72 °C for 300 s.
4. Hold: 4 °C.

PCR round 2 products were resolved on a 2.5% agarose gel for size selection and purified using the GeneJET Gel Extraction Kit, with a final elution volume of 50 µl in UltraPure distilled water.

## Supplementary Note 3. Golden Gate assembly cloning of cgRNAs

cgRNA sensor constructs were cloned from pairs of complementary DNA oligonucleotides using the following three-step procedure:

1. **Oligonucleotide phosphorylation and annealing:** Top- and bottom-strand oligonucleotides were phosphorylated and annealed in a single reaction using T4 polynucleotide kinase (T4 PNK; NEB, no. M0201L).
2. **Golden Gate assembly:** The annealed and phosphorylated oligonucleotide duplex was diluted 1:10 and incorporated into a Golden Gate assembly reaction with the corresponding cgRNA backbone plasmid using either BsmBI-v2 or PaqCI.
3. **Transformation and clone validation:** Assembly products were transformed into chemically competent NEB Stable Competent *E. coli* (NEB, no. C3040I). Ampicillin-resistant colonies were isolated, expanded and confirmed for correct insert incorporation by Sanger sequencing.

### Step 1. Oligonucleotide phosphorylation and annealing

#### Reaction setup

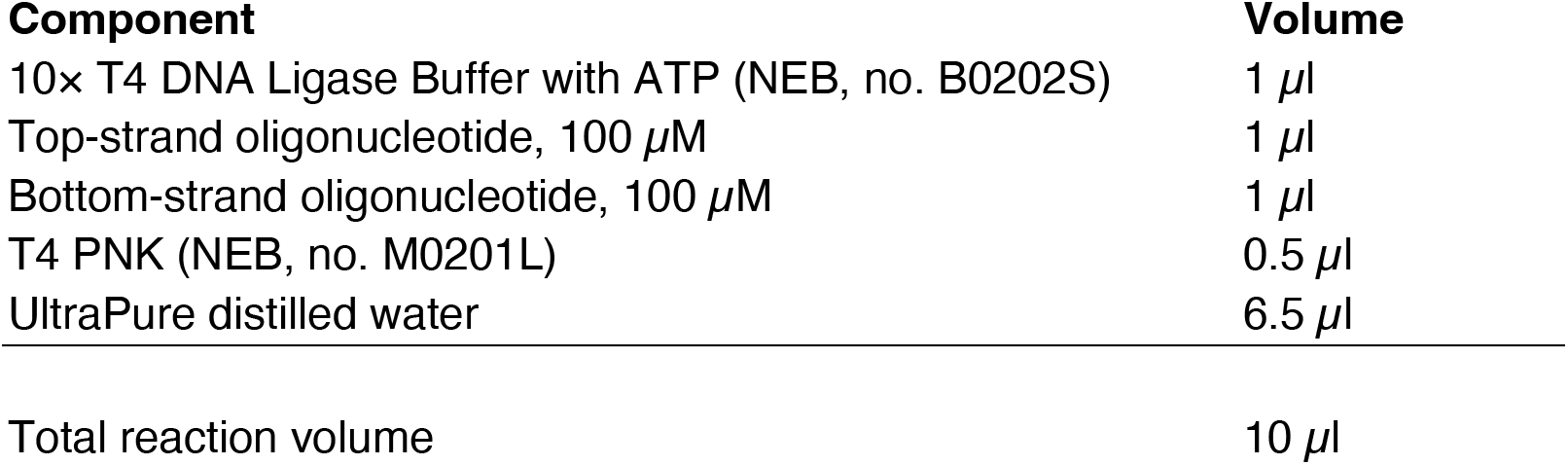

**Annealing protocol**

1. Incubate at 37 °C for 30 min.
2. Heat to 95 °C for 5 min.
3. Cool gradually to 25 °C at a rate of –1 °C every 12 s.

The annealed and phosphorylated oligonucleotide duplex was diluted 1:10 before Golden Gate assembly.

### Step 2a. Golden Gate assembly using BsmBI-v2

#### Reaction setup

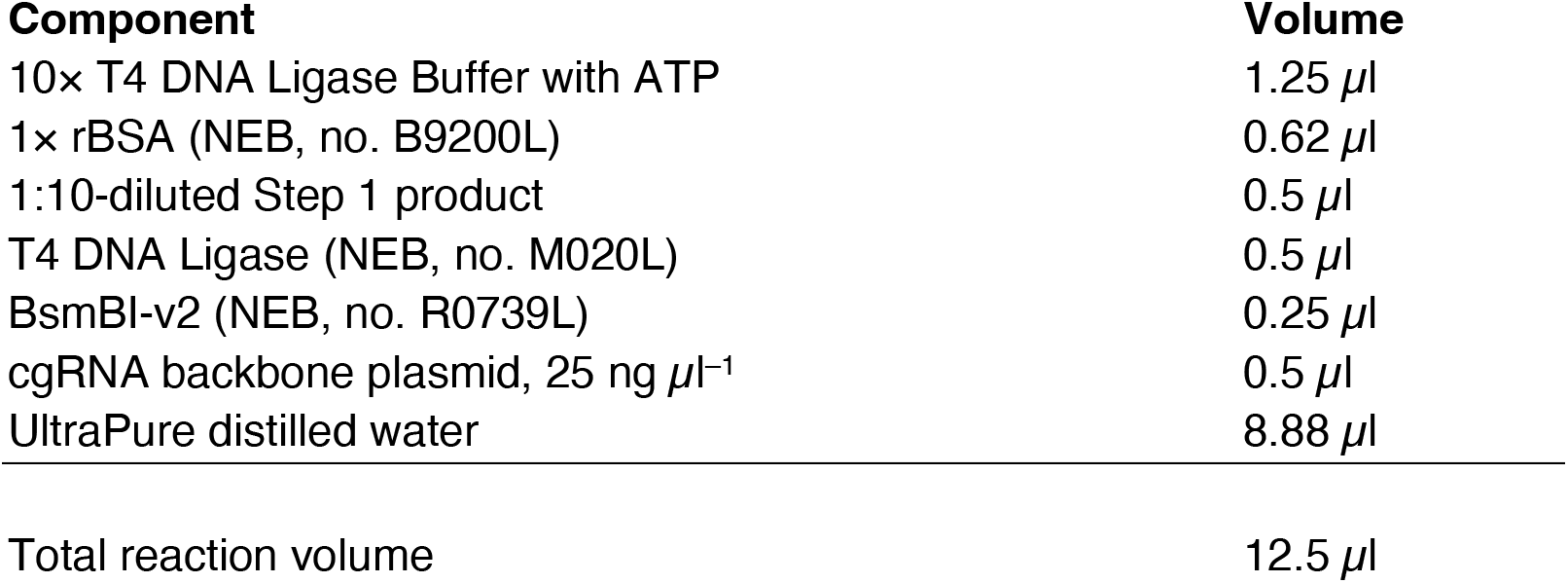

**Cycling protocol**

1. 30 cycles of:

`o` 42 °C for 1 min.
`o` 16 °C for 1 min.
2. Incubate at 55 °C for 30 min.
3. Hold at 4 °C.

### Step 2b. Golden Gate assembly using PaqCI

#### Reaction setup

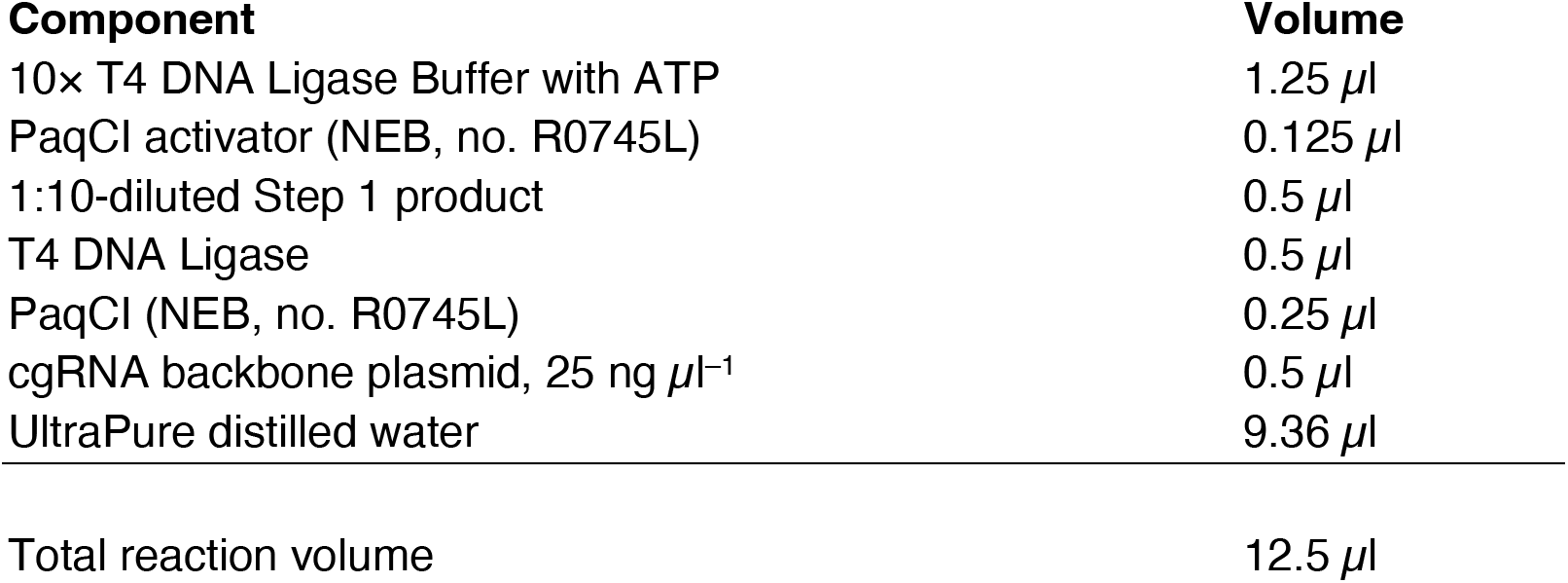

**Cycling protocol**

1. 30 cycles of:

`o` 37 °C for 1 min.
`o` 16 °C for 1 min.
2. Incubate at 60 °C for 30 min.
3. Hold at 4 °C.

### Step 3. Transformation and clone isolation

For transformation, 2 µl of the Golden Gate assembly product was added to 10 µl of chemically competent NEB Stable Competent *E. coli* (NEB, no. C3040I).

**Transformation protocol**

1. Incubate the DNA–bacteria mixture on ice for 30 min.
2. Heat shock at 42 °C for 30 s.
3. Recover on ice for 2 min.
4. Add NEB Stable Outgrowth Medium (NEB, no. B9035) and incubate for 1 h at 30 °C.
5. Plate the outgrowth on LB agar containing 100 µg ml^−1^ ampicillin and incubate overnight at 30 °C.
6. Pick individual colonies and expand them overnight at 30 °C in LB medium containing ampicillin.
7. Confirm correct insert incorporation by Sanger sequencing.

## Supplementary Note 4. Cloning of the mouse cgRNA library

A library of mouse miRNA targets was cloned into a lentiviral cgRNA backbone using the following three-step procedure:

1. **PCR amplification of the miRNA target library:** A Twist Bioscience-synthesized DNA oligonucleotide pool containing mouse miRNA targets was amplified by PCR.
2. **Golden Gate assembly:** The amplified miRNA targets were inserted into the cgRNA V3 lentiviral backbone LVcHS4-RFP-US1-cgRNA-V3-pGK-Puro.
3. **Transformation and library expansion:** The assembly product was transformed into chemically competent NEB Stable *E. coli* cells, followed by liquid-culture expansion and plasmid preparation.

### Step 1. PCR amplification of the mouse miRNA target library

#### Reaction setup

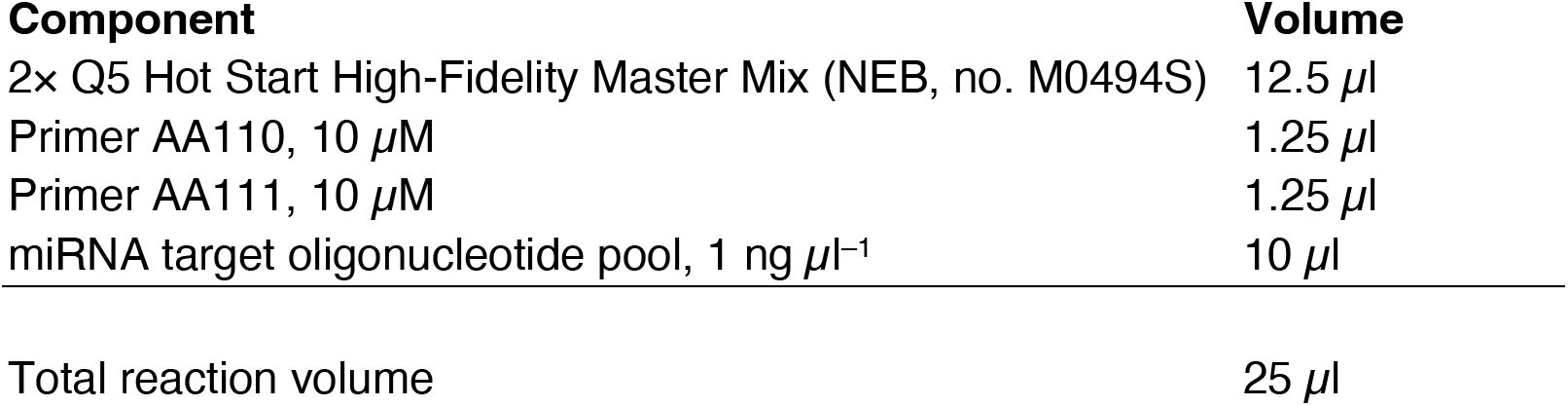

Two replicate PCR reactions were performed.

**Thermocycling protocol**

1. Initial denaturation: 98 °C for 30 s.
2. 12 cycles of:

- Denaturation: 98 °C for 10 s.
- Annealing: 71 °C for 30 s.
- Extension: 72 °C for 30 s.
3. Final extension: 72 °C for 300 s.
4. Hold: 4 °C.

The two PCR replicates were pooled and purified using the GeneJET Gel Extraction Kit (Thermo Fisher Scientific), with a final elution volume of 30 µl in ddH_2_O. To confirm the expected product size, 5 µl of the purified product was resolved on a 2% agarose gel at 100 V for 25 min before proceeding to Golden Gate assembly.

### Step 2. Golden Gate assembly of the mouse cgRNA library

#### Reaction setup

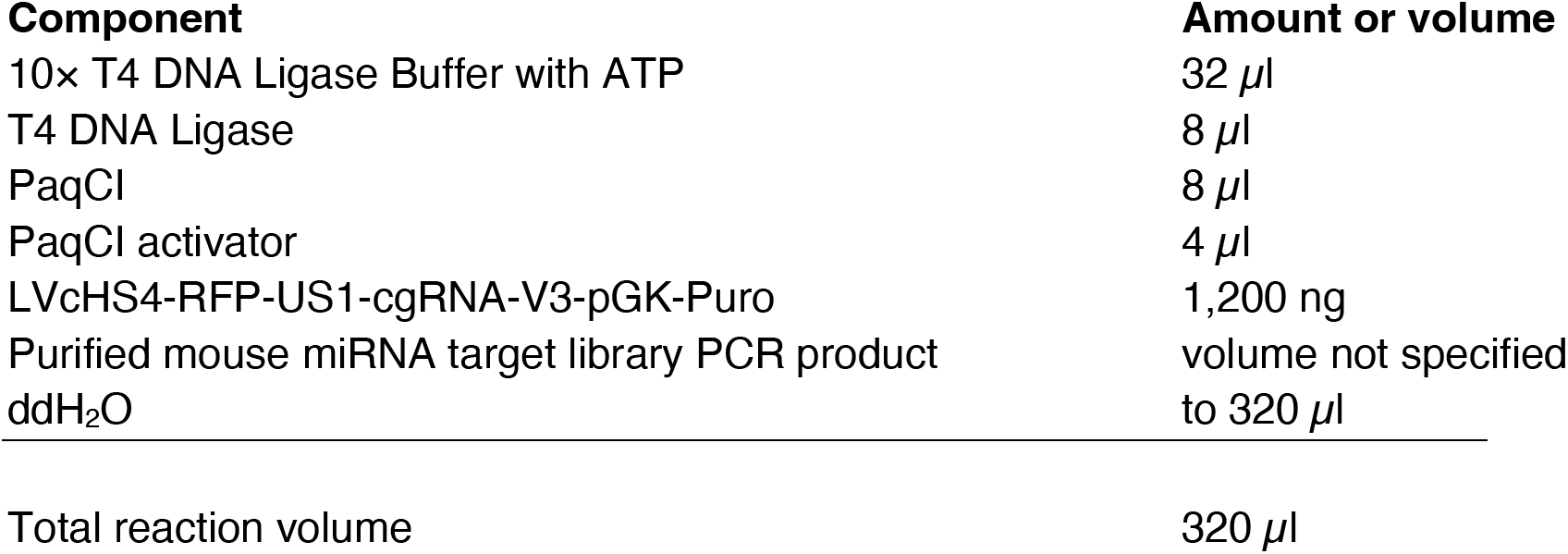

The reaction was distributed across eight PCR tubes at 40 µl per tube.

**Cycling protocol**

1. 30 cycles of:

a. 37 °C for 1 min.
b. 16 °C for 1 min.
2. Incubate at 60 °C for 30 min.
3. Hold at 4 °C.

The eight reactions were pooled and purified using the GeneJET Gel Extraction Kit, with a final elution volume of 50 µl in ddH_2_O.

### Step 3. Transformation and library expansion

For transformation, 50 µl of the purified Golden Gate assembly product was added to 200 µl of chemically competent NEB Stable *E. coli* cells.

**Transformation and expansion protocol**

1. Recover the transformed cells in 6 ml of NEB Stable Outgrowth Medium for 2 h at 30 °C with shaking under aerobic conditions.
2. Pellet the cells at 1,000 rpm.
3. Resuspend the pellet in 1 ml of NEB Stable Outgrowth Medium.
4. Transfer the resuspended cells into 200 ml of LB-ampicillin medium and culture overnight at 30 °C with shaking.
5. Divide the library outgrowth into 50-ml aliquots.
6. Pellet the aliquots by centrifugation at 4,000 rpm and either store the pellets at –80 °C or prepare plasmid DNA using the Qiagen Plasmid Midi Kit (Qiagen, no. 12143) according to the manufacturer’s protocol.

---

**Figure S1.**
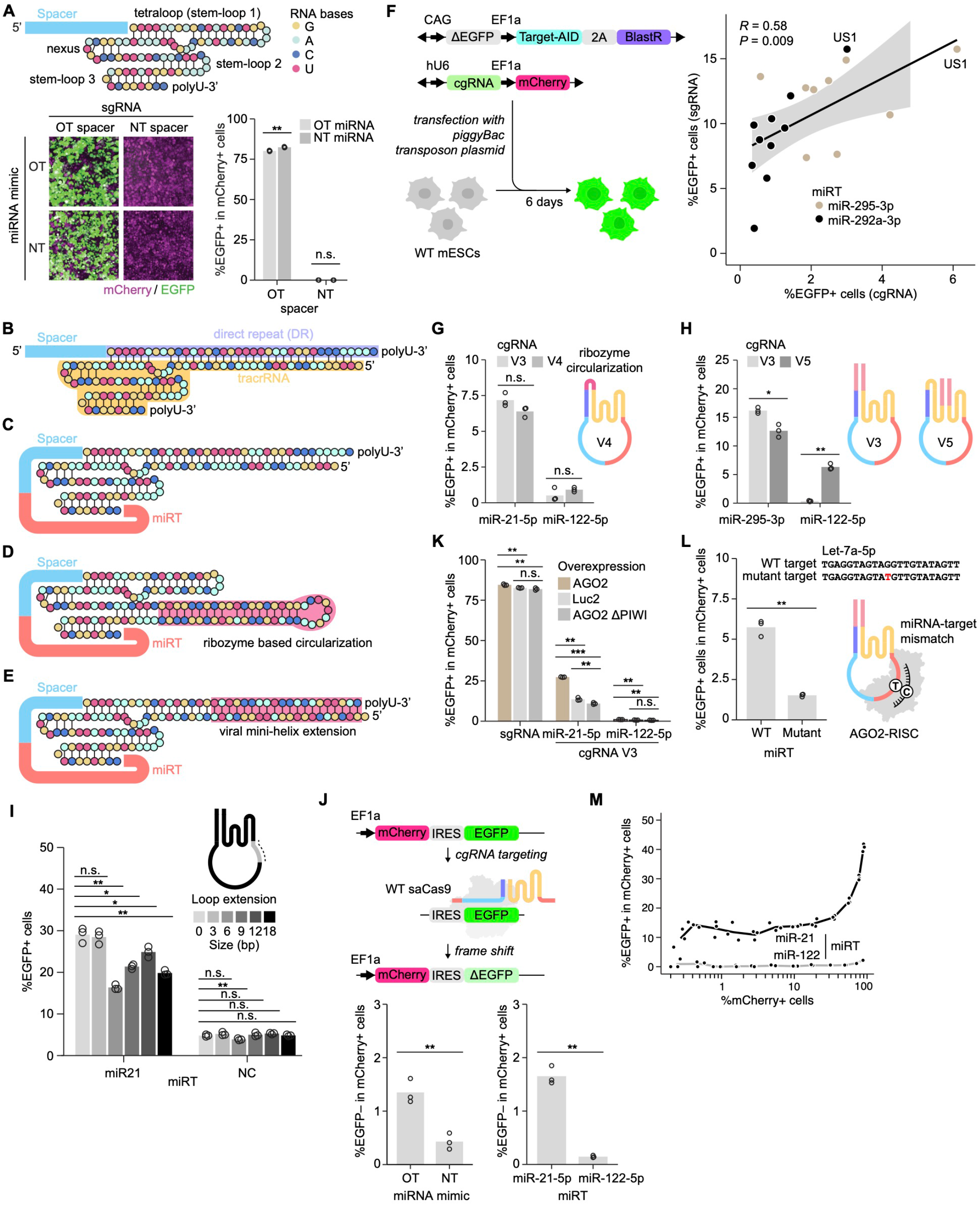
Design and functional characterization of cgRNA variants. (A) Constitutive base editing by sgRNA. Toolkit HeLa cells carrying an sgRNA with either a reporter-targeting or non-targeting spacer were transfected with miR-122 or a non-targeting miRNA mimic. Left, representative fluorescence micrographs. Scale bar, 100 μm. Right, percentage of EGFP+ cells among mCherry+ cells. Welch’s t-test compared miRNA mimic conditions within each spacer group (n = 3 replicates). (B–E) Predicted architectures of crRNA:tracrRNA (B), cgRNA V1 (C), ribozyme-circularized cgRNA V2 (D), and minihelix-extended cgRNA V3 (E). (F) Relationship between cgRNA activity and the editing efficiencies of the corresponding sgRNA spacers. mESCs were transfected with cgRNAs carrying different spacers and targets for miR-295-3p or miR-292a-3p. Each point represents the mean of two replicate measurements for one guide configuration. The line indicates a linear regression fit, gray shading indicates the 95% confidence interval, and Pearson’s *R* and associated *P* value are shown. (G) Comparison of cgRNA V3 and ribozyme-circularized cgRNA V4 in HeLa cells using targets for endogenous miR-21-5p or non-expressed miR-122-5p. Welch’s t-test compared V3 and V4 for each miRNA target (n = 3 replicates). (H) Comparison of cgRNA V3 and second-stem-circularized cgRNA V5 in mESCs using targets for endogenous miR-295-3p or non-expressed miR-122-5p. Welch’s t-test compared V3 and V5 for each miRNA target (n = 3 replicates). (I) Effect of increasing loop length on cgRNA activity in HeLa cells. cgRNAs targeting endogenous miR-21-5p or a negative-control sequence contained the indicated linker extensions between the guide RNA scaffold and miRNA target. Welch’s t-test compared each extension with the unextended construct within the corresponding target group (n = 3 replicates). (J) miRNA-dependent genome editing by cgRNA coupled to wild-type SaCas9. Left, editing following transfection with miR-122 or a non-targeting miRNA mimic. Right, editing by cgRNAs targeted by endogenous miR-21-5p or non-expressed miR-122-5p. The percentage of EGFP− cells among mCherry+ cells was quantified by flow cytometry. Welch’s t-test; n = 3 replicates. (K) Effect of AGO2 abundance on endogenous miRNA-dependent cgRNA activity. HeLa reporter cells expressing AGO2, firefly luciferase (Luc2), or AGO2ΔPIWI were transduced with sgRNA or cgRNAs targeting miR-21-5p or miR-122-5p. Welch’s t-test was used for the indicated pairwise comparisons (n = 3 replicates). (L) Effect of a central mismatch at the predicted cleavage site of the let-7a-5p target on cgRNA activity in mESCs. Welch’s t-test compared the fully complementary and mutant targets (n = 3 replicates). (M) Relationship between lentiviral transduction rate, estimated from the percentage of mCherry+ cells, and cgRNA-mediated base editing in HeLa cells for cgRNAs targeting miR-21-5p or miR-122-5p. n.s., not significant; \**P* < 0.05; \*\**P* < 0.01; \*\*\**P* < 0.001.

**Figure S2.**
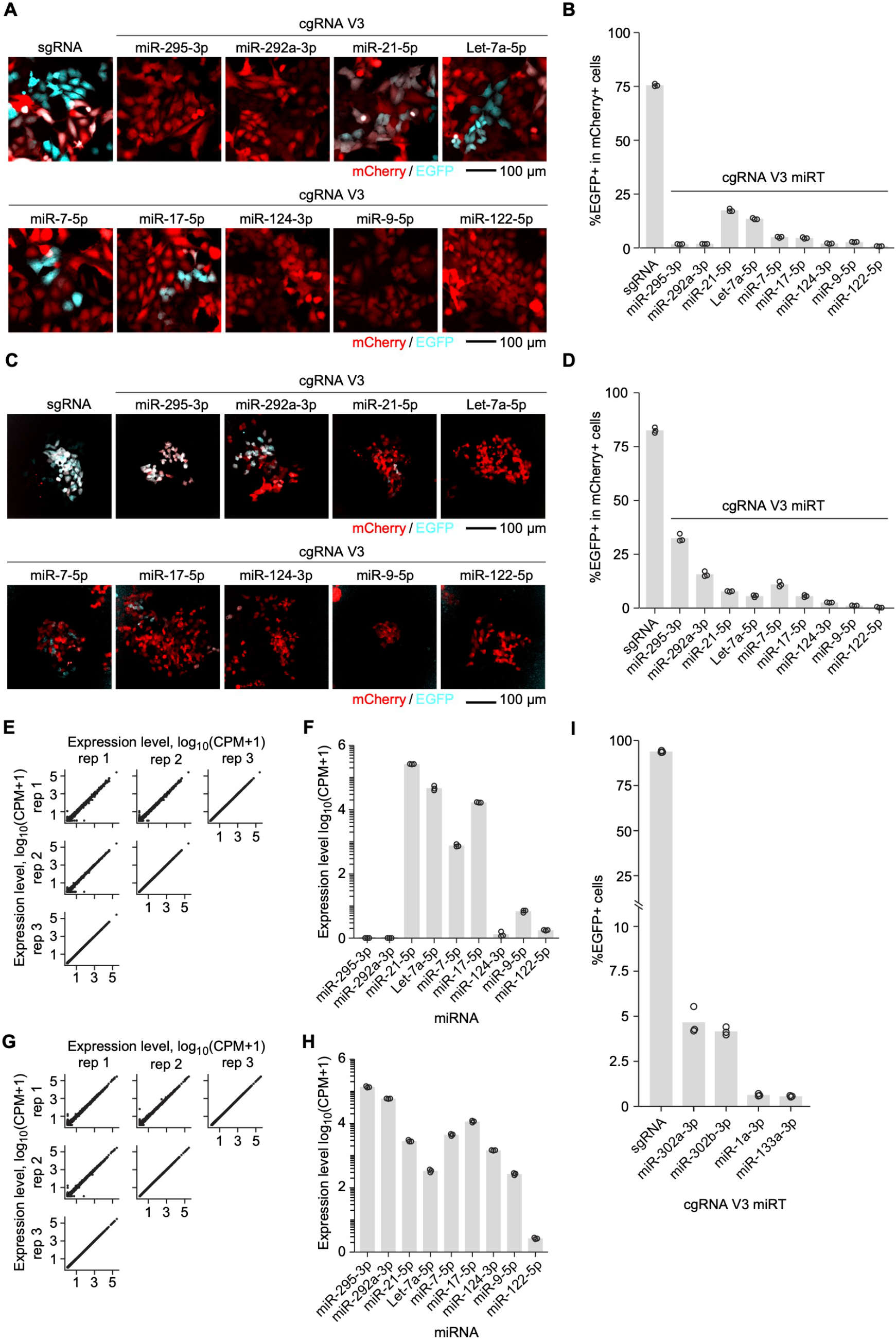
Endogenous miRNA-dependent cgRNA activity across mammalian cell types. (A, B) Endogenous miRNA-dependent base editing in toolkit HeLa cells transduced with sgRNA or cgRNA V3 targeted by indicated miRNAs. (A) Representative fluorescence micrographs showing mCherry and EGFP expression. Scale bar, 100 μm. (B) Percentage of EGFP+ cells among mCherry+ cells quantified by flow cytometry. Bars indicate means, and points represent individual replicate measurements (n = 3). (C, D) Corresponding analysis in toolkit mESCs. (C) Representative fluorescence micrographs. Scale bar, 100 μm. (D) Percentage of EGFP+ cells among mCherry+ cells quantified by flow cytometry. Bars indicate means, and points represent individual replicate measurements (n = 3). (E, F) AQ-seq analysis of miRNA abundance in HeLa cells. (E) Pairwise comparisons of log_10_-transformed counts per million plus one [log10(CPM + 1)] across three replicate libraries. (F) Mean abundance of the miRNAs targeted by the cgRNA panel. Bars indicate means, and points represent individual replicate measurements (n = 3). (G, H) Corresponding AQ-seq analysis in mESCs. (G) Pairwise comparisons of log_10_(CPM + 1) across three replicate libraries. (H) Mean abundance of the miRNAs targeted by the cgRNA panel. Bars indicate means, and points represent individual replicate measurements (n = 3). (I) Base editing in toolkit hiPSCs transduced with sgRNA or cgRNA V3 targeted by miR-302a-3p, miR-302b-3p, miR-1a-3p, or miR-133a-3p. The percentage of EGFP+ cells was quantified by flow cytometry. Bars indicate means, and points represent individual replicate measurements (n = 3).

**Figure S3.**
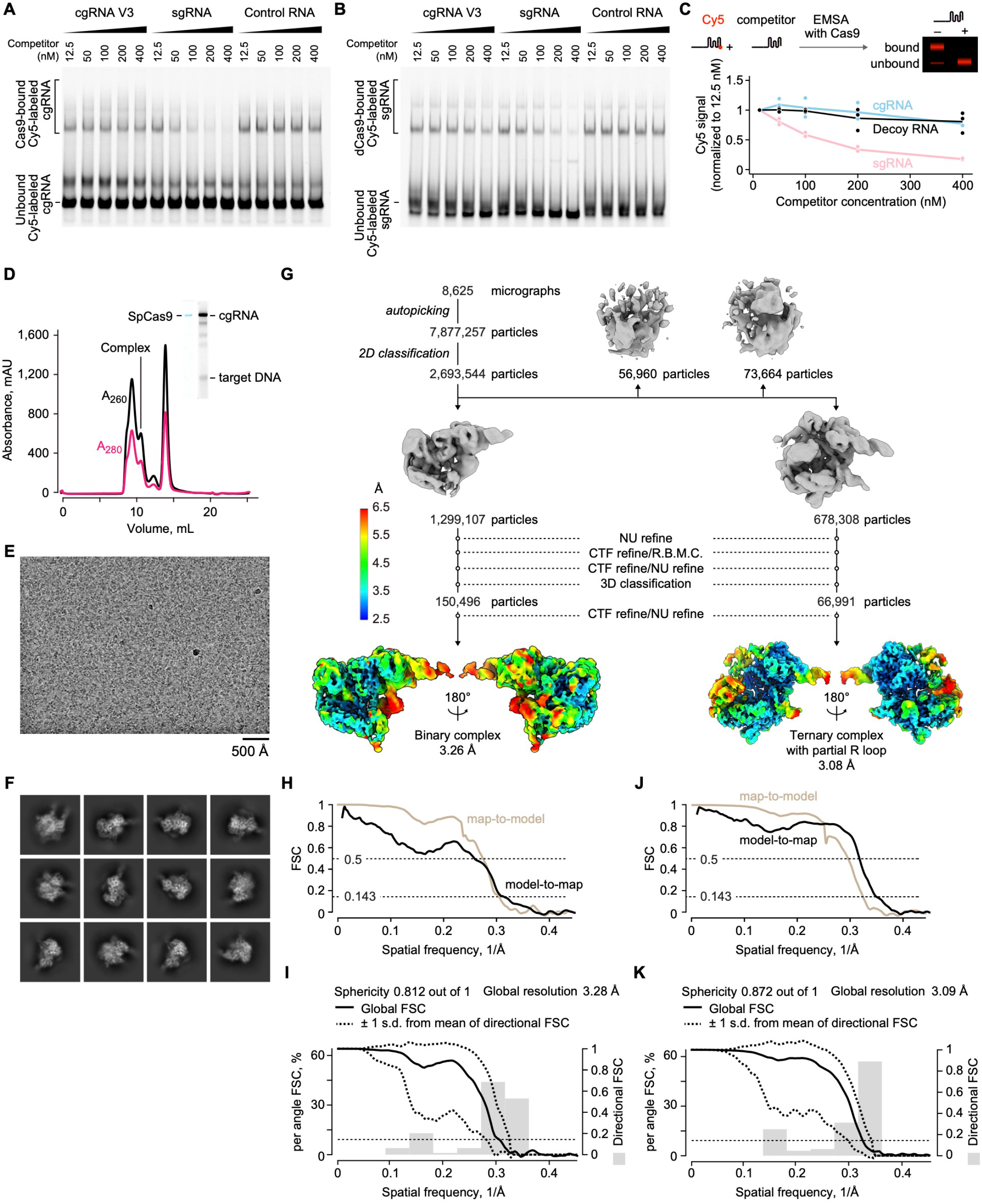
Biochemical and cryo-EM characterization of SpCas9–cgRNA complexes. (A, B) Competitive electrophoretic mobility shift assays using Cy5-labeled cgRNA (A) or sgRNA (B) in the presence of increasing concentrations of unlabeled cgRNA, sgRNA, or nonspecific competitor RNA. (C) Quantification of the Cas9-bound Cy5 signals in B. Values were normalized to the lowest concentration of the corresponding competitor RNA. (D) Size-exclusion chromatography of SpCas9 assembled with cgRNA and target DNA. The indicated complex-containing fraction was evaluated by SDS–PAGE and urea–PAGE and used for cryo-EM analysis. (E) Representative cryo-EM micrograph recorded using a 300-kV Titan Krios microscope equipped with a K3 detector. Scale bar, 500 Å. (F) Representative two-dimensional class averages. (G) Single-particle cryo-EM processing workflow yielding the SpCas9–cgRNA binary complex and the SpCas9–cgRNA–target DNA ternary complex containing a partial R-loop. Final density maps are colored according to local resolution. (H, I) Map-to-model and model-to-map Fourier shell correlation (FSC) curves (H) and directional FSC analysis (I) for the SpCas9–cgRNA binary complex. (J, K) Corresponding FSC curves (J) and directional FSC analysis (K) for the SpCas9–cgRNA– target DNA ternary complex. The global resolutions and map sphericity values are indicated in I and K.

**Figure S4.**
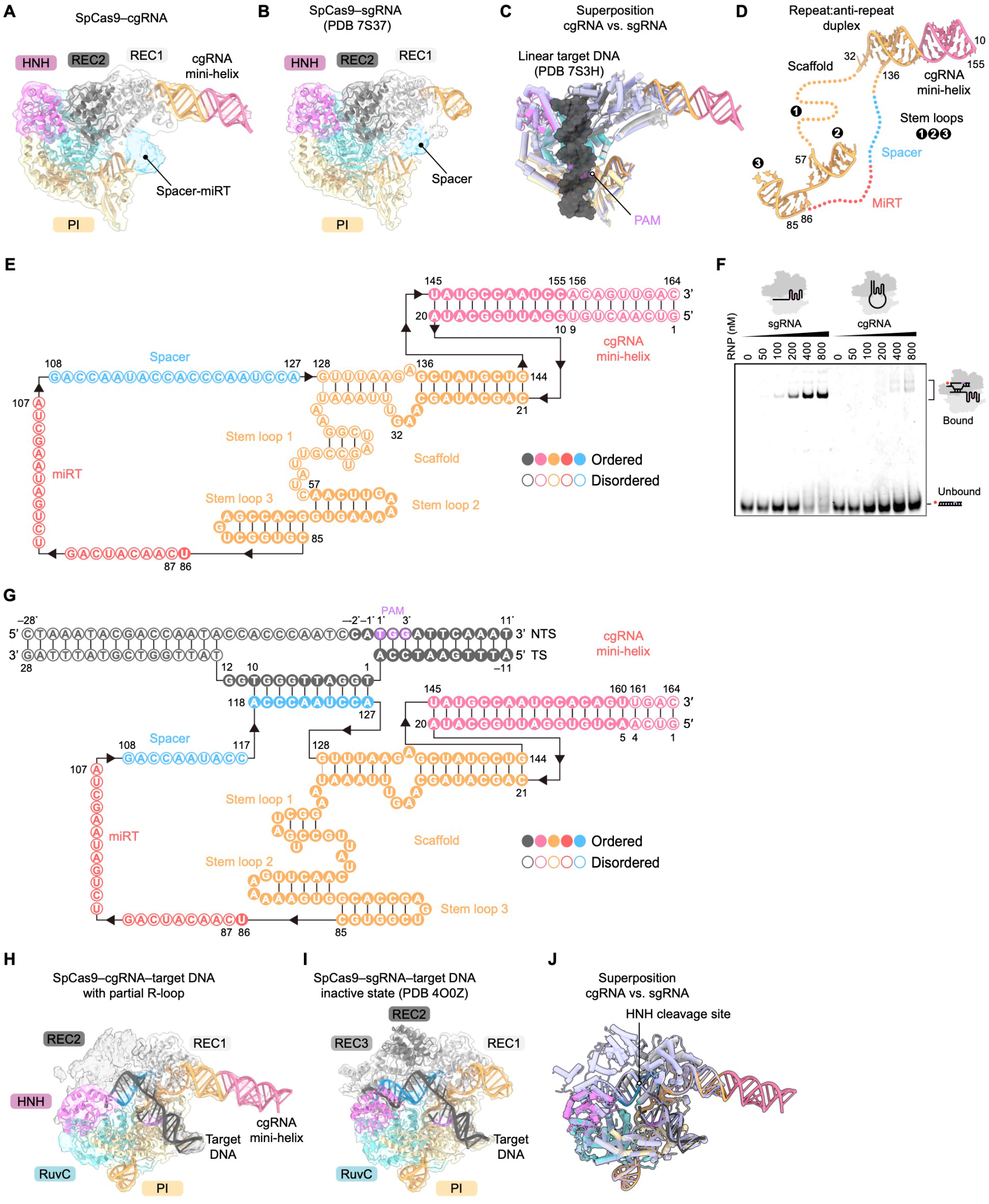
Structural comparison of cgRNA- and sgRNA-bound SpCas9 complexes. (A, B) Structural models of the SpCas9–cgRNA binary complex (A) and the SpCas9–sgRNA binary complex (B; PDB 7S37). The SpCas9–cgRNA model is overlaid with the corresponding cryo-EM density map. (C) Superposition of the SpCas9–cgRNA complex and the SpCas9–sgRNA complex. Linear target DNA from the PAM-bound SpCas9–sgRNA structure (PDB 7S3H) is shown for reference. RMSD, 1.15 Å across 739 equivalent Cα atoms. (D) Domain organization of cgRNA in the SpCas9–cgRNA binary complex. Ordered and disordered RNA regions are indicated by solid and dotted lines, respectively. (E) Secondary-structure model of cgRNA in the binary complex. Watson–Crick and noncanonical base pairs are indicated by black and gray lines, respectively; filled and open circles denote ordered and disordered nucleotides. (F) Electrophoretic mobility shift assay measuring binding of Cy5-labeled target DNA by SpCas9 assembled with sgRNA or cgRNA. The experiment was repeated three times with similar results. (G) Secondary-structure model of cgRNA and target DNA in the SpCas9–cgRNA–target DNA ternary complex containing a partial R-loop. TS, target strand; NTS, non-target strand. Base-pairing and nucleotide-order conventions are as in E. (H, I) Structural models of the SpCas9–cgRNA–target DNA ternary complex containing a partial R-loop (H) and the inactive SpCas9–sgRNA–target DNA complex (I; PDB 4O0Z). The cgRNA complex is overlaid with the corresponding cryo-EM density map. (J) Superposition of the complexes shown in H and I. RMSD, 0.80 Å across 991 equivalent Cα atoms.

**Figure S5.**
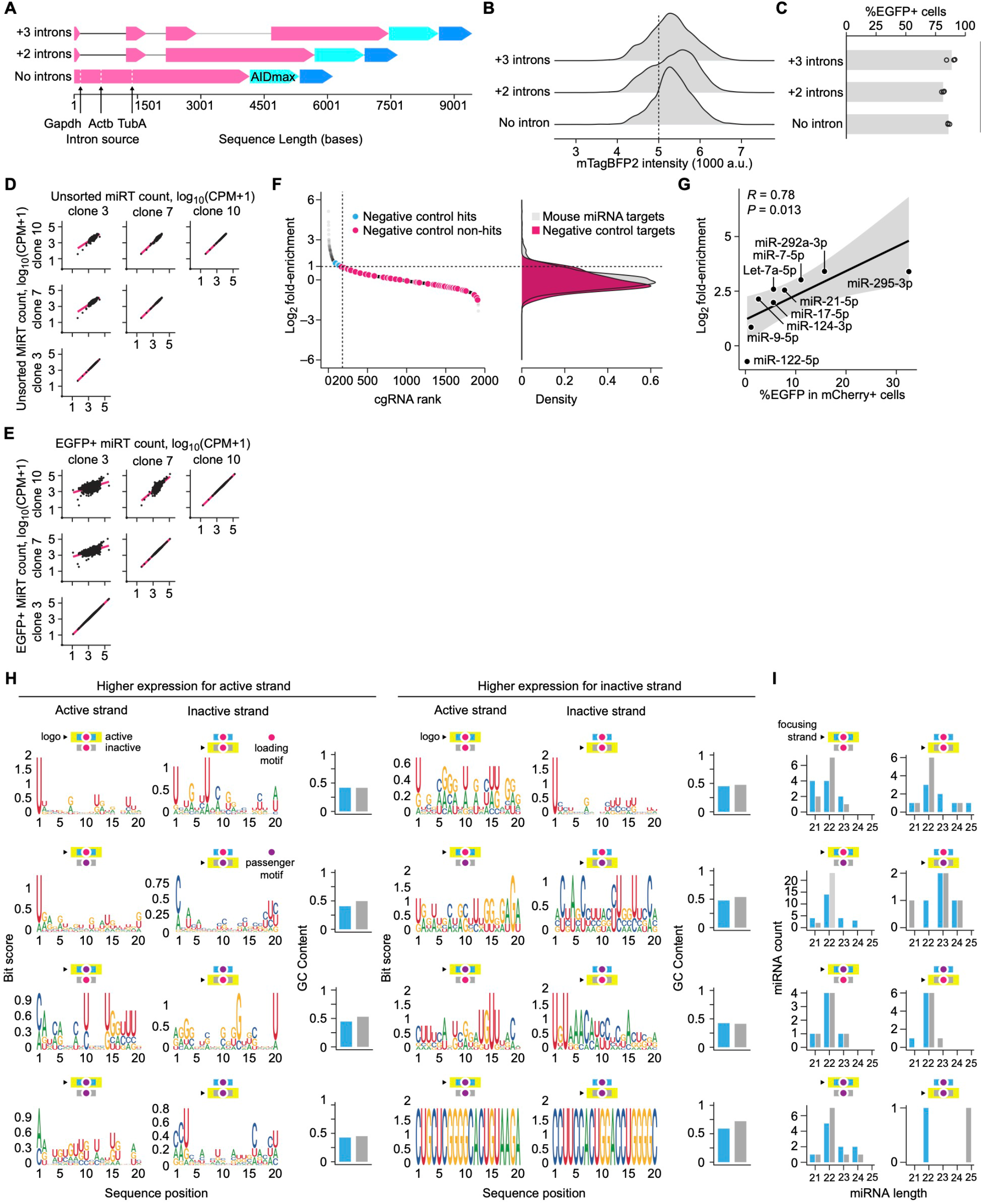
mESC pooled cgRNA screen and sequence features associated with miRNA strand activity. (A) The architecture of Target-AIDmax containing zero, two, or three mouse introns. Colors indicate the genes from which the introns were derived. (B) mTagBFP2 fluorescence distributions in the corresponding mESC reporter lines. (C) Base-editing activity of the reporter lines following sgRNA transduction. The percentage of EGFP+ cells was quantified by flow cytometry. Welch’s t-test was used to compare each intronic Target-AIDmax line with the no-intron control (n = 3 replicates). (D, E) Pairwise comparisons of log_10_-transformed cgRNA miRNA-target abundance [log10(CPM + 1)] among Clones 3, 7, and 10 in the unsorted (D) and EGFP+ (E) populations. (F) Ranked enrichment and density distributions of mouse miRNA targets and negative-control targets in the pooled screen. Negative controls meeting the hit criteria are distinguished from negative-control non-hits. The horizontal dashed line denotes log_2_ fold enrichment = 1.0. (G) Relationship between cgRNA enrichment in the pooled screen and the percentage of EGFP+ cells measured in individual cgRNA assays. Each point represents one miRNA target. The line indicates a linear regression fit, gray shading indicates the 95% confidence interval, and Pearson’s *R* and associated *P* value are shown. (H) Sequence logos and GC content of active and inactive strands from heteroactive miRNA pairs, stratified by whether the active or inactive strand exhibited higher AQ-seq abundance and by their combinations of loading- and passenger-motif assignments. Yellow outlines and arrows identify the focusing strand represented in each sequence logo. Blue and gray bars indicate the GC contents of the active and inactive strands, respectively. (I) miRNA-length distributions for the corresponding heteroactive-pair categories shown in H. Yellow outlines and arrows identify the focusing strand, and blue and gray bars indicate active and inactive strands, respectively.

**Figure S6.**
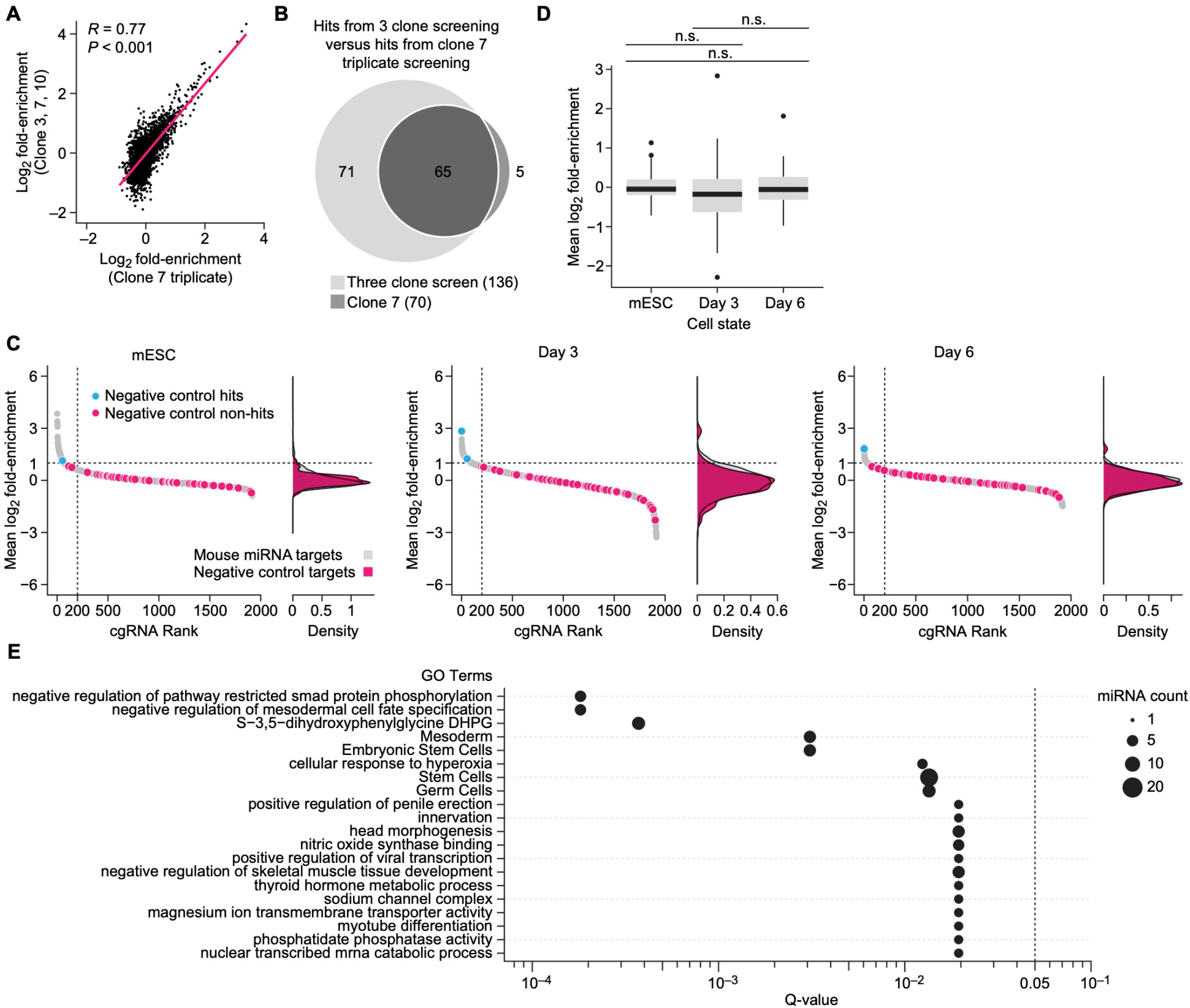
Validation of genome-wide cgRNA screening during SMC differentiation. (A) Correlation between mean cgRNA enrichment in the Clone 7 triplicate mESC screen and the three-clone mESC screen shown in Figure 3. Each point represents one miRNA target. The line indicates a linear regression fit, and Pearson’s *R* and associated *P* value are shown. (B) Overlap of miRNA targets identified as hits in the Clone 7 and three-clone mESC screens. Hits were defined by log2 fold enrichment ≥ 1.0 and FDR < 0.05. (C) Ranked enrichment and density distributions of mouse miRNA targets and negative-control targets in the Clone 7 screens performed in the mESC state and on Days 3 and 6 of SMC differentiation. Negative controls meeting the hit criteria are distinguished from negative-control non-hits. The horizontal dashed line denotes log_2_ fold enrichment = 1.0. (D) Distributions of negative-control cgRNA enrichment across the mESC, Day 3, and Day 6 screens. Mann–Whitney two-sided U-tests were used for the indicated pairwise comparisons. (E) Gene set enrichment analysis of miRNAs identified as hits in the Clone 7 mESC screen. Bubble size represents the number of miRNAs associated with each term, and the dashed line denotes Q = 0.05. n.s., not significant.

